# XMAP: Cross-population fine-mapping by leveraging genetic diversity and accounting for confounding bias

**DOI:** 10.1101/2023.03.30.534832

**Authors:** Mingxuan Cai, Zhiwei Wang, Jiashun Xiao, Xianghong Hu, Gang Chen, Can Yang

**Affiliations:** Department of Biostatistics, City University of Hong Kong; Guangzhou HKUST Fok Ying Tung Research Institute, Guangzhou 511458, China; Department of Mathematics, The Hong Kong University of Science and Technology; Shenzhen Research Institute of Big Data, Shenzhen 518172, China; The WeGene Company

## Abstract

Fine-mapping prioritizes risk variants identified by genome-wide association studies (GWASs), serving as a critical step to uncover biological mechanisms underlying complex traits. However, several major challenges still remain for existing fine-mapping methods. First, the strong linkage disequilibrium among variants can limit the statistical power and resolution of fine-mapping. Second, it is computationally expensive to simultaneously search for multiple causal variants. Third, the confounding bias hidden in GWAS summary statistics can produce spurious signals. To address these challenges, we develop a statistical method for cross-population fine-mapping (XMAP) by leveraging genetic diversity and accounting for confounding bias. By using cross-population GWAS summary statistics from global biobanks and genomic consortia, we show that XMAP can achieve greater statistical power, better control of false positive rate, and substantially higher computational efficiency for identifying multiple causal signals, compared to existing methods. Importantly, we show that the output of XMAP can be integrated with single-cell datasets, which greatly improves the interpretation of putative causal variants in their cellular context at single-cell resolution.

## Introduction

Genome-wide association studies (GWASs) have reported hundreds of thousands of associations between single-nucleotide polymorphisms (SNPs) and various phenotypes [1], but most reported SNPs reside in non-coding regions [2, 3, 4]. As the cell type and cellular process in which the identified SNPs are active remains largely unknown, the GWAS findings remain hard to interpret. Fine-mapping seeks to prioritize the causal SNPs underlying complex traits and diseases. Recent progress shows that, by integrating fine-mapping results and single-cell data, it becomes feasible to identify disease/trait-relevant cell types and cell states [5, 6]. Therefore, fine-mapping is a critical step to interpret GWAS findings by elucidating their biological mechanisms of identified risk variants, and fine-mapping results will offer an invaluable resource for precision medicine [7].

Despite the great promise of fine-mapping, efforts toward reliable prioritization of causal SNPs have been hampered by three key challenges. First, when GWAS samples come from a single population, SNPs in a local genomic region can be highly correlated due to the low recombination rates in that region. It is very difficult for statistical methods to distinguish the causal variants from a set of SNPs in strong linkage disequilibrium (LD). Second, genetic signals at trait-associated regions are commonly conferred by many variants acting together. A very recent study of 744 human expression quantitative trait loci (eQTLs) reported that 17.7% of the eQTLs harbour more than one variant with major effects on gene expression levels, emphasizing the importance of identifying multiple genetic variants within an associated locus [8, 9]. For example, an eQTL associated with *ERPA2* and Crohn’s disease was found to be driven by 13 separate variants [9]. However, it becomes computationally expensive to simultaneously search for multiple SNPs by enumerating causal combinations. Third, the unadjusted socioeconomic status [10] and geographic clustering [11, 12] in GWAS samples can induce confounding bias in GWAS estimates [13]. These confounding factors cannot be fully corrected through linear mixed models (LMMs) [14, 15] or principal component analysis (PCA) [16]. Fine-mapping without correcting the confounding bias in GWAS data can yield spurious results.

While many efforts have been devoted to the development of fine-mapping methods, existing methods only partially addressed the above major challenges. The classical fine-mapping methods [17, 18] rely on an exhaustive search for all possible causal configurations of vari-ants. They become computationally unaffordable when searching for more than three causal associations among thousands of variants. More efficient methods have been developed based on approximated inference, including CAVIARBF [19], FINEMAP [20], and DAP-G [21, 22]. A very recent method, SuSiE [23, 24], introduces a novel framework by assuming the overall genetic effects can be decomposed as a sum of single effects. The model structure of SuSiE enables an efficient algorithm to detect multiple causal SNPs with minor computational over-head. Despite their improvement in computational efficiency, the statistical power of these methods is usually limited because it is difficult for them to distinguish the causal variants from the highly correlated variants in the single population setting. To boost the statistical power of fine-mapping, several methods were developed to leverage different LD patterns with cross-population GWASs, including trans-ethnic PAINTOR [25] and MsCAVIAR [26]. Although these methods allow a locus to harbour multiple causal variants in principle, they require enumerating all causal combinations of variants, hence become too time-consuming to search for more than three causal variants. Furthermore, existing fine-mapping methods do not account for confounding bias in GWAS summary statistics, leading to spurious results.

In this paper, we develop a statistical method for cross-population fine-mapping (XMAP) by leveraging genetic diversity and accounting for confounding bias (Figure 1). The success of XMAP relies on its three unique features. First, XMAP can leverage distinct LD structures from genetically diverged populations. It is known that individuals from different population backgrounds usually have different LD structures. For example, individuals from the African (AFR) population are known to have narrower LD compared to those from the European (EUR) population [27]. By jointly analyzing cross-population GWASs, XMAP can effectively improve the power and resolution of fine-mapping. Second, XMAP can identify multiple causal signals with a linear computational cost, while many existing fine-mapping methods are too time-consuming to identify multiple causal signals. Third, XMAP can correct the confounding bias in GWAS summary data to avoid false positive findings and improve reproducibility.

**Figure 1:**
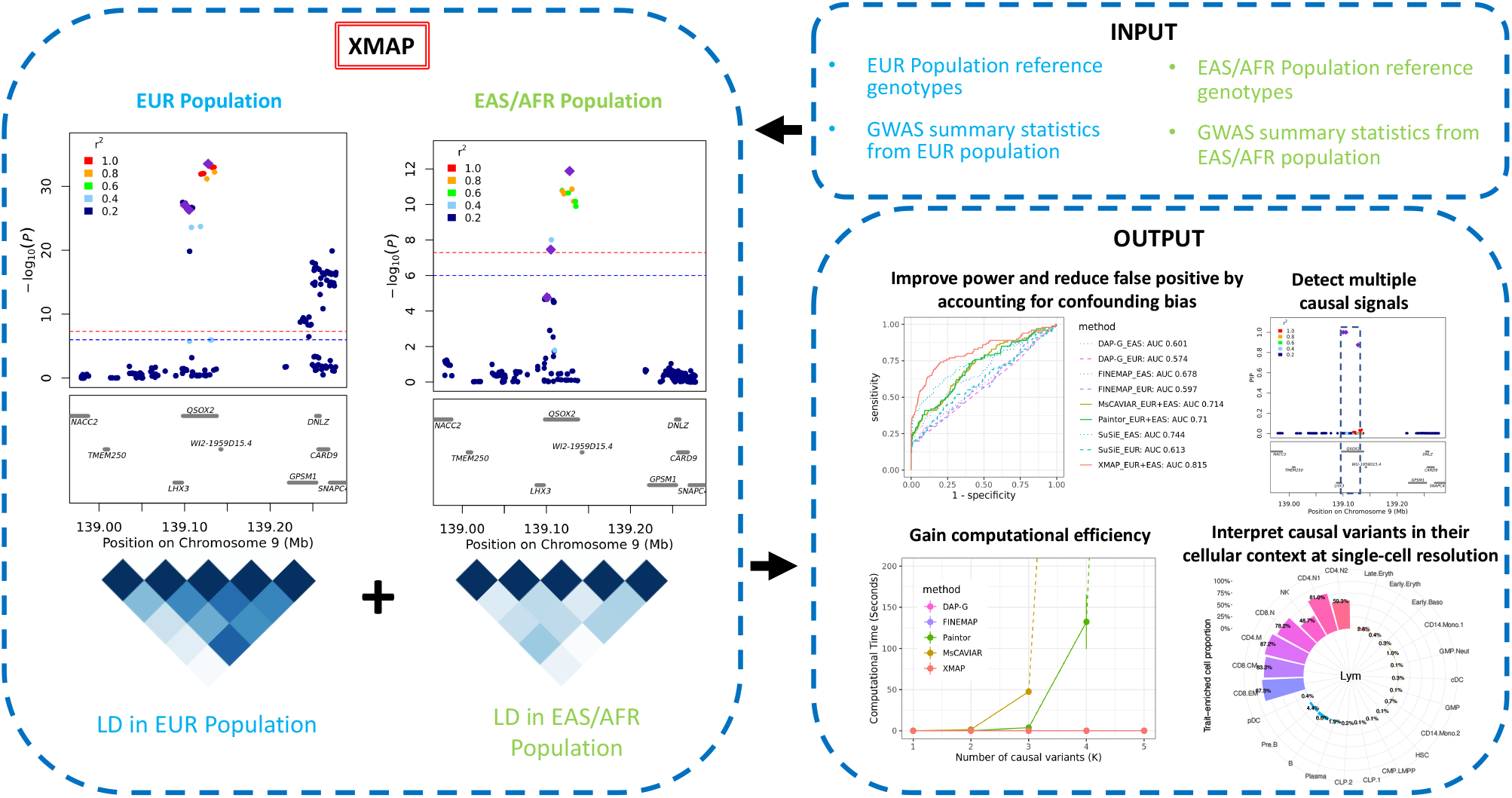
XMAP overview. XMAP takes the summary statistics and reference genotypes from multiple populations as inputs. XMAP can improve the statistical power of fine-mapping by leveraging the distinct LD pattern across populations while reducing false positives by accounting for confounding bias in GWAS summary statistics. Paired with a fast algorithm, XMAP is able to efficiently identify multiple causal signals. The fine-mapped SNPs can be integrated with single-cell datasets to identify trait-relevant cells.

Through comprehensive simulation studies, we show that XMAP not only improves the statistical accuracy of fine-mapping but also offers a substantial computational advantage over existing methods. The evidence from real data analysis indicates that XMAP achieves substantial power gain with high reproducibility. By combining the GWASs of low-density lipoprotein (LDL) from East Asian (EAS), African, and European, XMAP identifies three times more putative causal SNPs than SuSiE. These SNPs are strongly enriched in the eQTL of the liver, suggesting their important roles underlying the biological process of LDL. Furthermore, using the height GWAS as an example, we show that XMAP can effectively correct confounding bias and substantially improve reproducibility. Lastly but importantly, XMAP results can be integrated with single-cell data to identify trait-relevant cell populations at single-cell resolution, maximizing the utility of single-cell data for the inference of the pathological mechanisms. We apply XMAP to 12 blood traits and perform integrative analyses of the XMAP results and single-cell profiles of 23 hematopoietic cell populations. The analysis results suggest that XMAP enables the identification of the trait-relevant cell types in which putative causal SNPs are active. For example, SNPs identified by XMAP show a significant enrichment of the mean corpuscular volume in 99.3% of late-stage erythroid cells, which is very helpful to interpret GWAS results.

## Results

### Method overview

XMAP is a computationally efficient and statistically accurate method for fine-mapping causal variants using GWAS summary statistics. With innovations in its model and algorithm design, XMAP has three features: (i) It can better distinguish causal variants from a set of associated variants by leveraging different LD structures of genetically diverged populations. (ii) By jointly modeling SNPs with putative causal effects and polygenic effects, XMAP allows a linear-time computational cost to identify multiple causal variants, even in the presence of an over-specified number of causal variants. (iii) It further corrects confounding bias hidden in the GWAS summary statistics to reduce false positive findings and improve replication rates. The fine-mapping results given by XMAP can be further used for downstream analysis to illuminate the causal mechanisms at different cascades of biological processes, including tissues, cell populations, and individual cells. In particular, XMAP results can be effectively integrated with single-cell datasets to identify disease/trait-relevant cells. We provide the implementation of XMAP in an efficient and freely available R package at https://github.com/YangLabHKUST/XMAP. The technical details of XMAP are described in the Methods section.

### Simulation study

We conducted comprehensive simulation studies to compare the performance of XMAP with several related fine-mapping methods, including DAP-G, FINEMAP, SuSiE, PAINTOR and MsCAVIAR. To mimic realistic LD patterns in different populations, we used genotypes of EUR samples from UKBB and genotypes of EAS samples from a Chinese cohort [28, 29]. We considered a region between the base pair position 45,202,602 and 45,435,202 in chromosome 22 (GRCH37), which comprises *p* = 500 SNPs. To demonstrate the benefit of leveraging genetic diversity in different populations, we selected three candidate SNPs that satisfy the following properties: (i) In EUR population, they are in high LD (i.e., with absolute correlation *>* 0.9) with at least three non-causal SNPs. (ii) In EAS population, they are weakly correlated with non-causal SNPs (i.e., have an absolute correlation *>* 0.6 with less than two non-causal SNPs). The heat maps in Figure 2 B show the absolute correlation between the three candidate causal SNPs and their neighboring SNPs. We investigated *K*_*true*_ causal SNPs, where *K*_*true*_ ∈ {1, 2, 3}, we randomly sampled *K*_*true*_ from the three candidate SNPs as the causal ones. To mimic the unbalanced composition of GWAS samples in global populations, we considered *n*_2_ = 20, 000 samples from the EUR population and explored different sample sizes *n*_1_ from the EAS population: 5,000, 10,000, 15,000, and 20,000. For reference LD matrices, we used the EUR LD matrix estimated with 337,491 British UKBB samples provided in a recent study [30] and estimated the EAS LD matrix with 35,989 EAS samples from the Chinese cohort [28]. We designed our simulations in two scenarios. First, we illustrated the benefit of cross-population fine-mapping by generating GWAS data without confounding bias. In the second scenario, we examined the effectiveness of XMAP in correcting confounding bias by simulating GWAS summary data with unadjusted sample structure.

**Figure 2:**
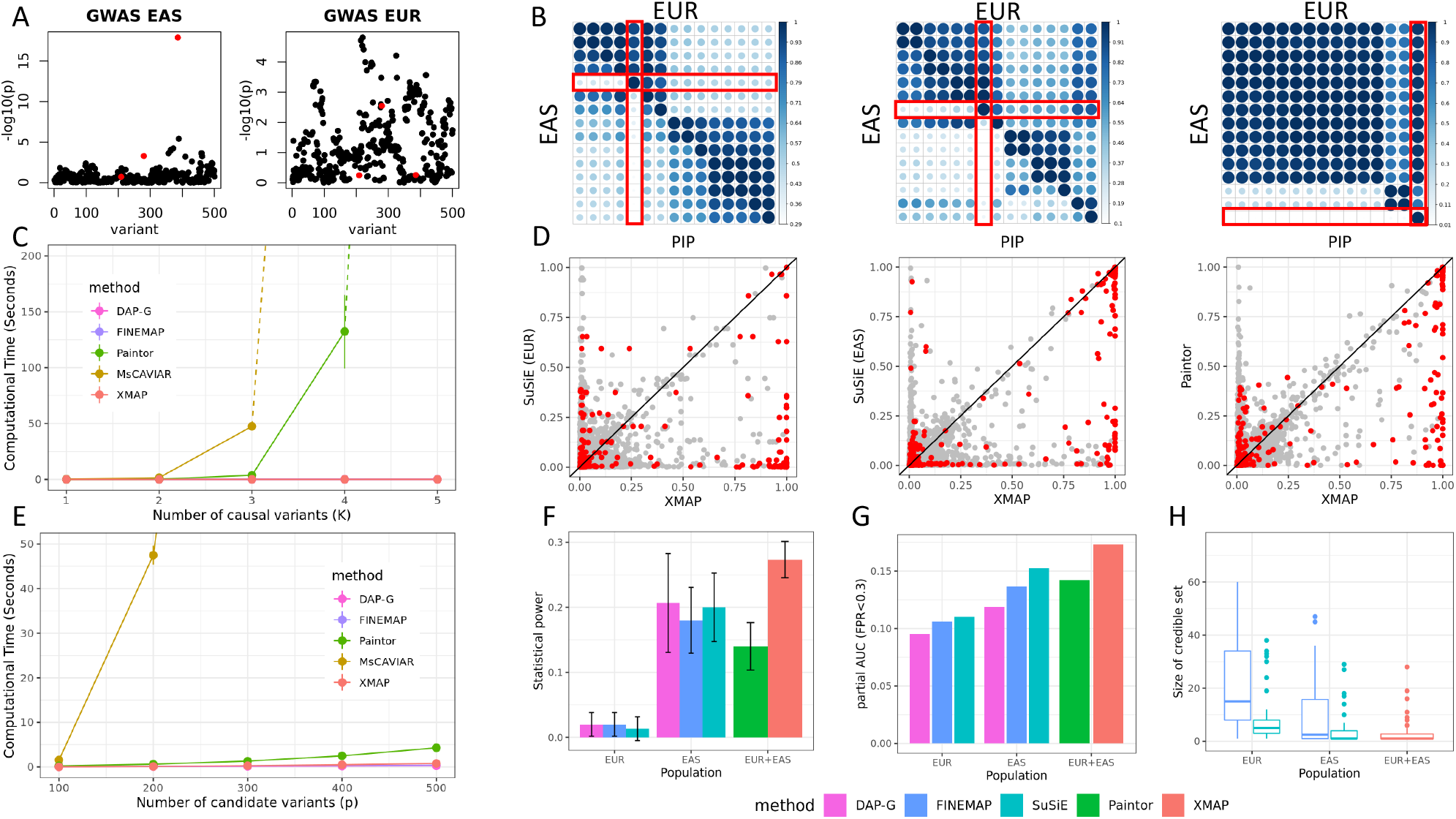
Comparisons of fine-mapping approaches in GWAS without confounding bias. (A) Manhattan plots of a simulated GWAS data in EAS (left) and EUR (right). (B) Heat maps showing the absolute correlations between the three causal SNPs (highlighted with rectangles) and their nearby SNPs in EAS and EUR populations. (C) CPU timings of XMAP, MsCAVIAR, PAINTOR, FINEMAP, and DAP-G are shown for increasing *K* with *p* = 100. Solid lines are CPU time recorded in our experiments and dashed lines represent predicted CPU time based on the time complexity of corresponding approaches. (D) Comparisons of PIP between XMAP and SuSiE, and between XMAP and PAINTOR. Red points represent true causal SNPs, and gray points represent SNPs with no effect. (E) CPU timings are shown for increasing *p* with *K* = 2. (F-H) Comparisons of statistical power (F), partial AUC with false positive rate*<* 0.3 (G), and level-95% credible set size (H) with *n*_1_ = *n*_2_ = 20, 000 and *K*_*true*_ = 3. Results are summarized from 50 replications.

We first consider the scenario in the absence of confounding bias. Specifically, we generated the polygenic effects with 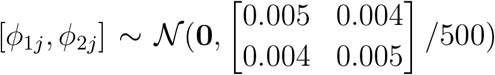 for *j* = 1, …, 500*di*, where 0.005 is the total heritability contributed by polygenic effects of the 500 SNPs in the locus, with a per-SNP heritability 0.005*/*500 = 10^−5^ and a genetic correlation 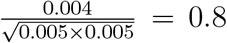between two populations. Then, we simulated the causal effects in the two populations with 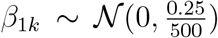 and 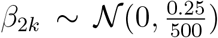 for *k* = 1, …, *K*_*true*_. This specification means that each causal SNP has a 0.25*/*0.005 = 50 fold per-SNP heritability enrichment compared to non-causal SNPs, and the effect sizes of SNP *k* are not necessarily the same across the two populations. The *K*_*true*_ causal SNPs jointly contribute 0.25*/*500 × *K*_*true*_ = 5 × 10^−4^ × *K*_*true*_ heritability. We obtained the standardized genotype matrices 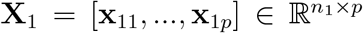 and 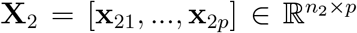, whose columns have zero mean and unit variance. Given the genotypes and effect sizes, we generated quantitative phenotypes in the two populations with 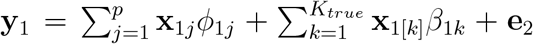 and 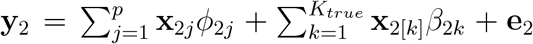, where **x**_1[*k*]_ and **x**_2[*k*]_ are the columns of **X**_1_ and **X**_2_ corresponding to the *k*-th causal SNP, and 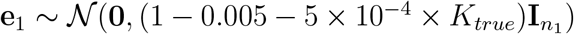 and 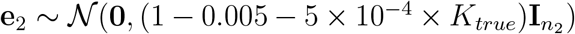 are independent noise in the two populations, respectively. Finally, we computed the GWAS summary statistics by marginally regressing the simulated phenotypes on each SNP for each population (Figure 2 A). The details of data pre-processing and parameter settings of XMAP and compared methods are given in the Supplementary Note.

Using a posterior inclusion probability (PIP) threshold of 0.9, we first evaluated the statistical power of compared methods. Figure 2 F shows the comparison of statistical power when *K*_*true*_ = 3 and *n*_1_ = *n*_2_ = 20, 000. Clearly, XMAP was the overall winner with the highest statistical power averaged across 50 replicates. In practice, we are usually more interested in the performance of fine-mapping when the false positive rate is small. Here, we evaluated the sensitivity and specificity under various PIP thresholds and generated the receiver operating characteristic (ROC) curve. As shown in Figure 2 G, DAP-G, SuSiE and FINEMAP only have a partial area under ROC curve (pAUC) around 0.1 when they were applied to EUR GWAS with the false positive rate (FPR)*<* 0.3. They achieved a higher pAUC when applied to the EAS GWAS because the causal variants were less correlated with non-causal variants in the EAS samples. For cross-population methods, we examined the performance of PAINTOR and MsCAVIAR. Because MsCAVIAR was too time-consuming to include more than two causal variants, we only applied MsCAVIAR to the setting with *K*_*true*_ ∈ {1, 2}. The results in Figure 2 F-G and Supplementary Figures 1-3 indicate that XMAP is more powerful than PAINTOR and MsCAVIAR in the existence of polygenic effects. In our additional simulation without polygenic effects (Supplementary Figures 8-13), XMAP could still achieve comparable performance with PAINTOR and MsCAVIAR because we allow the polygenic effects to be adaptively estimated from the data. To further investigate the difference in fine-mapping performance, we contrasted the PIP obtained by XMAP with those obtained by other methods (Figure 2D and Supplementary Figures 4-6). Clearly, XMAP produced substantially higher PIP for causal variants, as compared to SuSiE and PAINTOR, suggesting that XMAP could better distinguish causal SNPs from non-causal SNPs. This explains our observation that XMAP often yields higher pAUC and statistical power. We also assessed resolution of fine-mapping by evaluating the size of credible sets. The smaller credible sets, the higher resolution of fine-mapping. Here we consider XMAP, FINEMAP and SuSiE because they are the only methods that can provide credible sets for individual causal signals. As summarized in Figure 2 H, XMAP and SuSiE were the only two methods that could produce level-95% credible sets with a median size of two when they were applied to EAS GWAS. We used *K* = 5 for XMAP in the main results and investigated *K* = 10 in the Supplementary Figure 1-3. Under both settings, XMAP had consistent performance and steadily outperformed compared methods, suggesting its robustness to the specification of *K*. More comparisons under different settings of *n*_1_, *n*_2_ and *K*_*true*_ are provided in the Supplementary Figures 1-3.

To investigate the computational efficiency, we evaluated the CPU time of compared methods under different setting of *K* and *p*. As shown in Figure 2 D, the computational cost of MsCAVIAR and PAINTOR increases exponentially with both *K* and *p*. When analyzing a locus with *p* = 100 SNPs, MsCAVIAR could only include *K* ≤ 4 causal signals and PAINTOR could only include *K* ≤ 5 causal signals. It took more than one week for them to finish the analysis when more signals were included. By contrast, the computational cost of XMAP is linear to *K*, which makes it highly efficient when applied to locus with multiple causal SNPs. To identify multiple causal signals, the computational efficiency of XMAP allows us to set *K* to a large value (e.g., *K* = 10) when *K*_*true*_ is unknown. While DAP-G and FINEMAP had CPU times comparable to XMAP, they could not leverage cross-population GWASs to improve fine-mapping. This benchmark was evaluated using a Linux computing platform with 20 CPU cores of Intel (R) Xeon (R) Gold 6152 CPU at 2.10 GHz processor.

In the second set of simulations, we focus on fine-mapping of GWAS data in the presence of uncorrected confounding bias. We introduced sample structures to GWAS data by using the genotype principal components following a previous work [31]. Specifically, we first performed PCA on the genotypes of EAS and EUR samples separately and extracted the first principal components from the two populations as representations of sample structures, denoted as 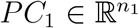 and 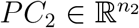, respectively. We re-scaled *PC*_1_ to have mean zero and variance 0.05 and re-scaled *PC*_2_ to have mean zero and variance 0.2. These variance values were selected to introduce proper level of inflation in the summary statistics. Next, we generated quantitative phenotypes with 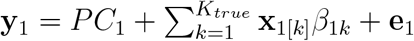 and 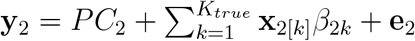, where the generating distributions of *β*_1*k*_ and *β*_2*k*_ are the same as those in the first scenario and the independent errors were generated with 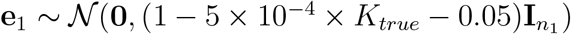 and 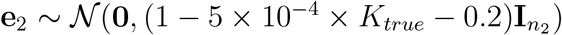. Finally, we simulated GWAS summary data by regressing phenotype vectors on each SNP without including the PCs as covariates. Figure 3 B shows the inflation constants in the simulated GWASs of the two populations evaluated btuy estimated LDSC intercepts *ĉ*_1_ and *ĉ*_2_. The inflation constants were substantially larger than one, indicating strong confounding bias. The confounding bias became stronger when the sample size increased, suggesting an exacerbated inflation in GWAS summary statistics. By accounting for the confounding bias, XMAP achieved the best overall performance across different PIP thresholds among compared methods. For example, when *K*_*true*_ = 2, *n*_1_ = 5, 000 and *n*_2_ = 20, 000, XMAP produced the highest AUC (0.784), as shown in Figure 3 C. When we focus on the the ROC curve with FPR*<* 0.3 (Figure 3 A), XMAP also achieved the highest pAUC. These results suggest that XMAP can improve statistical power while controlling the false positive rate. The pAUC evaluated under other simulation settings are summarized in the Supplementary Figure 7. Here we showed an concrete example with a single causal signal in Figure 3 D as an illustration. With uncorrected confounding bias, the GWAS *p*-values were inflated in the left regions of the locus (top panels of Figure 3 D). Without accounting for the confounding bias, SuSiE produced a false positive signal (SNPs in blue circles in the middle right panel of Figure 3 D) and assigned a high PIP≈ 0.6 for a null SNP. By adjusting the estimation error of GWAS effects based on inflation constants *ĉ*_1_ and *ĉ*_2_, XMAP effectively reduced the PIP of SNPs related to the false positive signal and correctly excluded the false positive signal from level-95% credible sets (left region in the bottom right panel of Figure 3 D). When we forced XMAP to ignore the inflation by setting *ĉ*_1_ = *ĉ*_2_ = 1, the false positive signal appeared in the output (bottom left panel of Figure 3 D), indicating the confounding bias was not properly adjusted. This observation implies the effectiveness of using the inflation constants to correct confounding bias in GWAS.

**Figure 3:**
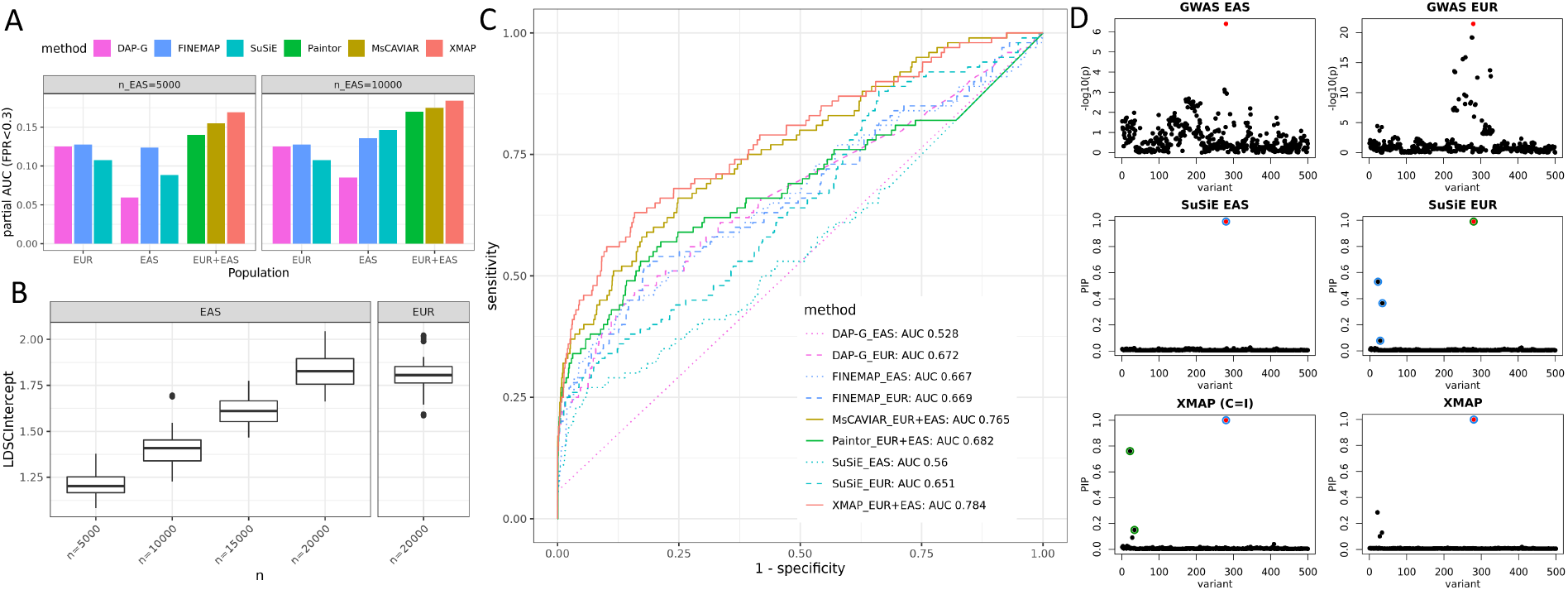
Comparisons of fine-mapping approaches in the presence of confounding bias. (A) Comparison of pAUC (FPR*<* 0.3) of fine-mapping among XMAP, PAINTOR, MsCAVIAR, SuSiE, FINEMAP, and DAP-G with *K*_*true*_ = 2 and sample size *n*_1_ ∈ {5, 000, 10, 000} in EAS. (B) Estimated LDSC intercepts *ĉ*_1_ (EAS) and *ĉ*_2_ (EUR) with sample size *n*_2_ = 20, 000 in EUR and *n*_1_ ∈ {5, 000, 10, 000, 15, 000, 20, 000} in EAS. (C) ROC curves of XMAP, PAINTOR, MsCAVIAR, SuSiE, FINEMAP, and DAP-G with *K*_*true*_ = 1, *n*_1_ = 5, 000, *n*_2_ = 20, 000. (D) An illustrative example generated by simulation. The first row shows the − log_10_(*p*)-value in the GWAS of EAS(left) and EUR (right). The second row shows the PIP obtained by applying SuSiE to the training data of EAS (left) and EUR (right). The third row shows the PIP obtained from XMAP by setting *ĉ* = *ĉ*_2_ = 1 (left) and estimating *c*_1_ and *c*_2_ from the data (right). Red dots represent causal SNPs. Circles in the same color represent SNPs in the level-95% credible sets of a causal signal. Results are summarized from 50 replications.

### Real data analysis

We performed fine-mapping to identify putative causal SNPs of complex traits with cross-population GWASs. First, by applying XMAP to LDL GWASs, where the magnitude of confounding bias was ignorable, we illustrated XMAP’s superior performance in improving fine-mapping power and resolution. Second, to investigate the ability of XMAP in correcting confounding bias, we applied XMAP to combine height GWASs from an EAS cohort [28] and the British cohort in UKBB, which was known to be affected by population structure [11, 12]. Through replication analysis, we compared the credibility of XMAP fine-mapped SNPs with related methods. Third, with the confounding bias properly corrected, we showed that XMAP enables the identification of multiple causal signals within a locus. Lastly but importantly, we integrated the fine-mapping output of XMAP in blood traits with single-cell data. With the improved fine-mapping results, we can have a better interpretation of risk variants in their relevant cellular context, gaining biological insights of causal mechanisms at single-cell resolution.

### XMAP improves fine-mapping by leveraging genetic diversity

We first applied XMAP to analyze LDL by combining GWASs form EUR, EAS, and AFR. As discovery cohorts, we used the GWASs of AFR and EAS released by the Global Lipids Genetics Consotium (GLGC), which were obtained based on 92,934 AFR samples and 71,150 EAS samples, respectively. For EUR, we considered two GWAS datasets: the UKBB GWAS summary data released by the Neale Lab with a sample size of 343,621, and the EUR GWAS data from GLGC with a sample size of 664,450. These GWAS summary statistics included 11,569,928-35,328,891 genotyped and imputed autosomal SNPs, minimizing the risk of omitting causal variants. Details of GWAS summary statistics are summarized in Supplementary Table 1. For EAS and EUR, we used the same reference LD matrices as in our simulation studies. For AFR, we estimated the LD matrices by using 3,072 African individuals from UKBB as reference samples. We followed a previous work [30] to partition all autosomal chromosomes into 2,763 consecutive loci, each with a width of 1 million base pairs (Mbp). To fully account for LD when analyzing each 1 Mbp locus, we included all SNPs in an extended region that also covers 1 Mbp before the starting position and 1 Mbp beyond the ending position of the locus, leading to a 3 Mbp extended region. We excluded the MHC region (25.5Mbp-33.5Mbp in chromosome 6) and two other long-range LD regions (8Mbp-12Mbp in chromosome 8 and 46Mbp-57Mbp in chromosome 11) because many spurious results were reported in these regions [30]. We applied XMAP to all regions that have more than 100 SNPs after overlapping the reference LD matrices with GWAS data. Because SuSiE often achieved the best performance among single-population methods in our simulation studies, we applied SuSiE to the GWAS of each population separately, serving as a baseline for comparison. We set *K* = 10 in XMAP and SuSiE for all loci.

We first quantified the confounding bias in these GWAS data using the estimates of LDSC intercepts. As shown in Supplementary Table 1, the LDSC intercepts estimated from all LDL GWASs were not substantially different from one, suggesting ignorable confounding bias here. We then summarized the fine-mapped SNPs in Figure 4 A. By combining GWAS data from different populations, XMAP consistently identified more causal signals than SuSiE with different PIP thresholds. Specifically, XMAP identified 149 SNPs with PIP*>* 0.8 and 145 SNPs with PIP*>* 0.9 when the GWASs from all three populations were jointly analyzed, which was three times more than the number of SNPs identified by SuSiE in EUR (50 SNPs with PIP*>* 0.8 and 45 SNPs with PIP*>* 0.9). The complete fine-mapping results are available at https://github.com/YangLabHKUST/XMAP/results.

**Figure 4:**
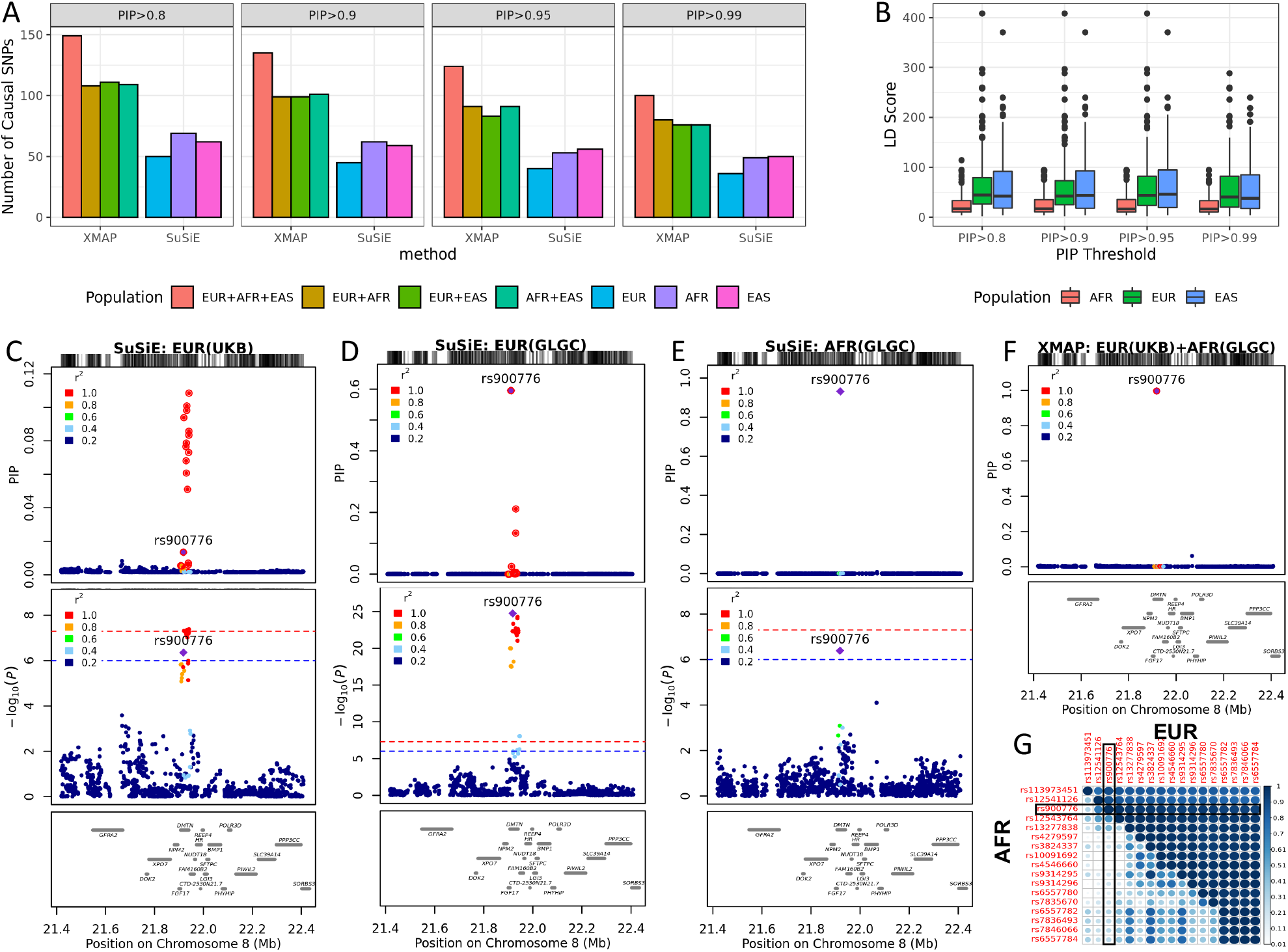
Application of XMAP and SuSiE to LDL. (A) Number of causal signals identified by XMAP and SuSiE with PIP thresholds 0.8, 0.9, 0.95, and 0.99. Colors represent different combination of GWAS training data. (B) LD score distribution of causal SNPs identified by XMAP. (C-F) Fine-mapping of locus 21.4Mbp-22.4Mbp in chromosome 8. The fine-mapping methods and training data are labelled on top of each panel. Top panels show the PIP. SNPs within 99% credible set are highlighted with red circles. Middle panels show the − log_10_(*p*−value) in GWAS. Red dashed lines represent 5 × 10^−8^. Blue dashed lines represent 1 × 10^−6^. Bottom panels annotate the position of genes in the locus. (G) Absolute correlation in EUR and AFR among the SNPs within level-99% credible set as shown in the red circles of (C). The SNP rs900776 is highlighted in the heat map.

The improved statistical power of XMAP could be atttributed to its capacity of leveraging genetic diversity. To see this, we checked the LD score which is a summation of squared correlation between a SNP and other SNPs in a population. A large LD score of a SNP means that this SNP has strong LD with many other SNPs. We observed that the XMAP fine-mapped SNPs have smallest LD scores in AFR (Figure 4 C), suggesting the power gain of XMAP could be attributed to the weak LD between causal SNPs and non-causal SNPs in AFR. As an example, rs900776 is an intronic variant in the *DMTN* region, which is highly correlated with surrounding SNPs in EUR. Because of this, SuSiE estimated the PIP of rs900776 as small as 0.002 using UKBB GWAS and produced very large 99% credible set that included 16 other SNPs for this signal. When applying SuSiE to the larger EUR GWAS data from GLGC, the PIP of SNP rs900776 increased from 0.002 to 0.6 (Figure 4 D). Different from the LD pattern in European population, rs900776 is less correlated with nearby SNPs in African population (Figure 4 G). Therefore, when SuSiE was applied to AFR GWAS, the estimated PIP of rs900776 increased to 0.9 (Figure 4 E). Unlike SuSiE that analyzes a single population at a time, XMAP enables joint analysis of EUR and AFR GWASs. XMAP successfully identified SNP rs900776 with a PIP as high as 0.99, yielding a high resolution credible set which contains rs900776 only (Figure 4 F). This indicates the improved power and resolution of XMAP by leveraging genetic diversity. We verified our findings with the expression quantitative trait loci (eQTLs) of liver obtained from the Genotype-Tissue Expression (GTEx) project [32]. As demonstrated in Supplementary Figure 14, the SNPs identified by XMAP produced a substantially higher enrichment of LDL in the liver eQTLs, as compared to SNPs identified by SuSiE using single population GWASs.

### XMAP enables the correction of confounding bias in fine-mapping

To demonstrate the effectiveness of XMAP in correction of confounding bias, we applied XMAP to the height GWASs which were well known to be affected by population structure [11, 12]. Following the previous cross-population fine-mapping pipeline [33], we first applied fine-mapping methods to discovery GWAS datasets, and then evaluated the credibility of fine-mapped SNPs in replication datasets from different population backgrounds. Here, we used the EUR GWAS from UKBB and a Chinese GWAS in our previous study [28] as discovery cohorts. For replication, we considered a recently released within-sibship GWAS from European population, which was known to be less confounded by population structure. We also included the GWAS from BBJ cohort as a replication data from EAS background. To ensure the SNP density, these GWASs were imputed to cover 3,776,576-12,515,778 variants (see Supplementary Table 1). The LDSC intercepts of UKBB GWAS and BBJ GWAS were estimated as 1.66 (s.e.=0.042) and 1.39 (s.e.=0.024), respectively, indicating the presence of strong confounding bias. The LDSC intercepts of EUR Sibship GWAS and Chinese GWAS were estimated as 1.07 (s.e.=0.0089) and 1.12 (s.e.=0.012), suggesting that the confounding bias is nearly ignorable. To investigate the ability of XMAP in accounting for confounding bias, we used UKBB and Chinese GWASs as inputs of XMAP and used SuSiE to analyze these GWAS data separately as benchmarks.

We summarized the replication rates of fine-mapped SNPs in Figure 5. Among the overlapped SNPs between the EUR Sibship GWAS and discovery cohorts, SuSiE detected 306 SNPs with PIP*>* 0.8 from UKBB GWAS. However, only 14.1% (43/306, Figure 5 A) were found to be genome-wide significant and only 13.4% of them (41/306, Figure 5 B) had PIP*>* 0.1 in the EUR Sibship replication cohort. The low replication rate suggests that these SNPs could be false positive signals due to unadjusted confounding bias. By accounting for the confounding bias, XMAP successfully reduced the number of false positive signals. For example, using PIP*>* 0.8 as a threshold, 21.4% (44/206) SNPs detected by XMAP were genome-wide significant and 21.4% (44/206) had PIP*>* 0.1 in BBJ replication cohort. A similar pattern can be observed in the BBJ replication cohort. With a PIP threshold of 0.8, only 23.9% (54/226, Figure 5 C) SNPs detected from UKBB GWAS by SuSiE were genome-wide significant and 8.8% (19/226, Figure 5 D) had PIP*>* 0.1 in BBJ GWAS. As a comparison, 42.3% (71/168) SNPs detected by XMAP were genome-wide significant and 14.9% (25/168) had PIP*>* 0.1 in BBJ replication cohort. The higher replication rate of XMAP implies its effectiveness of fine-mapping by accounting for confounding bias.

**Figure 5:**
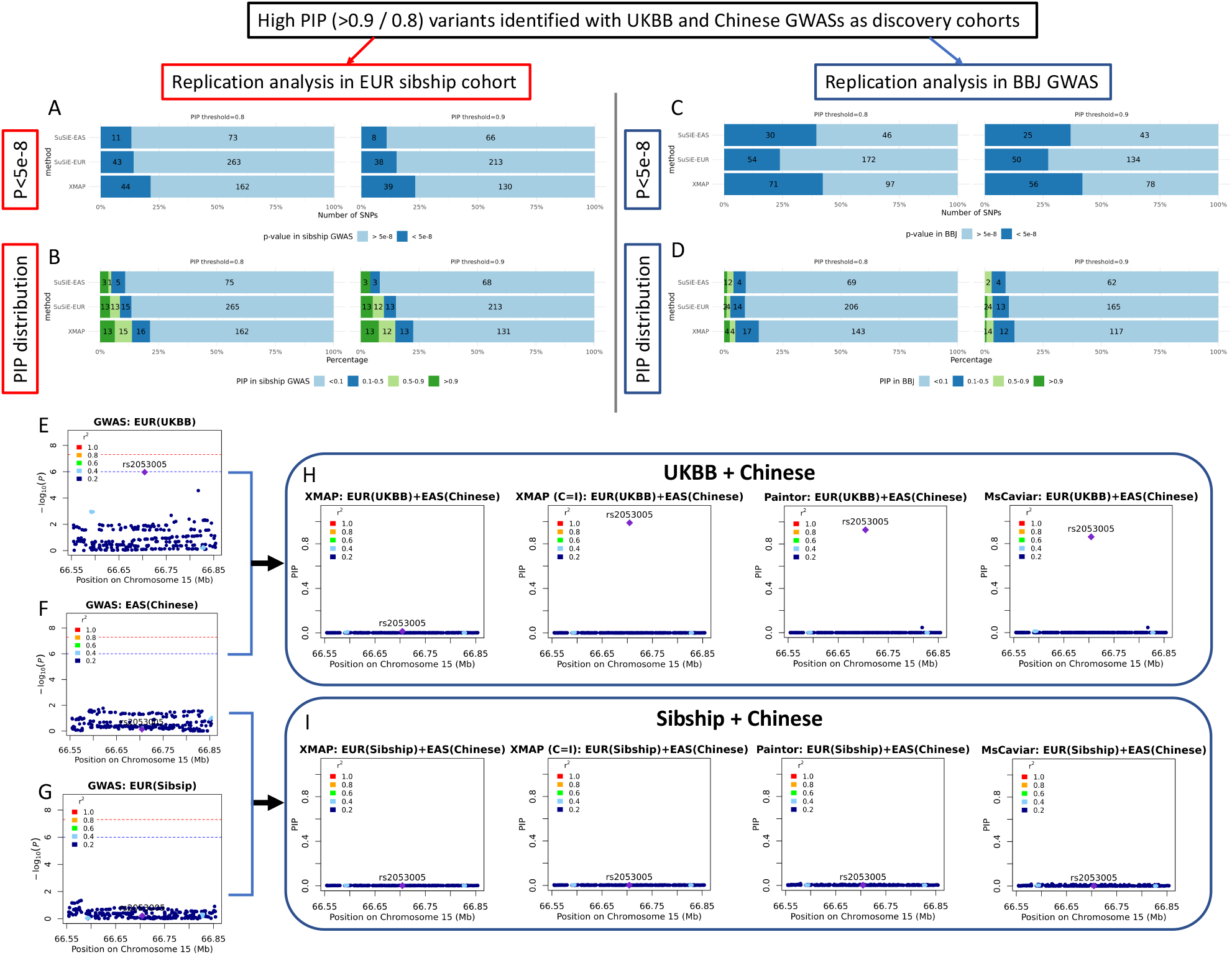
Replication analysis of XMAP and related methods on height GWASs. (A-D) Overview of replication analyses of high-PIP fine-mapped SNPs across populations: bar charts showing the fraction and number of fine-mapped SNPs with *p*-value*<* 5 × 10^−8^ in the replication cohorts of EUR Sibship GWAS (A) and BBJ (C) cohorts and bar charts showing the distribution of PIP for fine-mapped SNPs computed by SuSiE in the replication cohorts of EUR Sibship GWAS (B) and BBJ (D). (E-I) Fine-mapping of locus 66.55 Mbp-66.85Mbp in chromosome 15. The SNP rs2053005 is significant (*p*-value*<* 1 × 10^−6^) in UKBB (E), but not significant in Chinese GWAS and EUR Sibship GWAS (F and G). When UKBB and Chinese cohorts were combined for cross-population fine-mapping (H), the PIP of rs2053005 was computed to be *>* 0.8 by PAINTOR, MsCAVIAR and XMAP when we set *c*_1_ = *c*_2_ = 1 (XMAP C=I). XMAP estimated the inflation constants of UKBB and BBJ as 1.66 and 1.39, suggesting they are influenced by confounding bias. After correcting for confounding bias, this signal was excluded in XMAP with a PIP*<* 0.05, which suggests that the high PIP of the SNP could have been induced by uncorrected population stratification. To test our assumption, we combined Chinese and EUR Sibship GWASs, which are both less influenced by confounding factors (both with inflation constant estimated as 1.07). As expected, all methods consistently produced a low PIP for rs2053005 (I), which confirmed our assumption and suggested XMAP can reduce spurious signals.

Although PAINTOR and MsCAVIAR can also integrate cross-population GWASs, they are too time-consuming to analyze all loci on the genome. Here, we consider a concrete example to compare the performance of cross-population methods in the presence of confounding bias (Figure 5). For XMAP, we considered two settings: (i) the standard XMAP that used the estimated inflation constants (*c*_1_ and *c*_2_) to correct the confounding bias; (ii) a special case of XMAP forced not to correct the confounding bias by setting *c*_1_ = *c*_2_ = 1, denoted as ‘XMAP (C=I)’. In this example, the SNP rs2053005 locating at the locus 66.55 Mbp-66.85Mbp in chromosome 15 was significantly associated (*p*-value*<* 10^−6^) in UKBB GWAS (Figure 5 E), but not significant in both Chinese GWAS and EUR Sibship GWAS (Figure 5 F and G). When UKBB and Chinese cohorts were combined for cross-population fine-mapping, the PIP of rs2053005 was computed to be *>* 0.8 by PAINTOR, MsCAVIAR and XMAP (C=I) without accounting for confounding bias. After correcting for confounding bias, the PIP of this signal dramatically decreased in XMAP with a PIP*<* 0.05, which suggests that the high PIP of the SNP could have been caused by population stratification (Figure 5 H). To test our assumption, we applied cross-population methods to combine Chinese and EUR Sibship GWASs, both of which are known to be less influenced by population structure. As expected, all methods consistently yielded a low PIP for rs2053005 (Figure 5 I). This observation confirmed our assumption that rs2053005 could be a false positive and XMAP was able to exclude this signal by correcting the confounding bias.

### XMAP enables identification of multiple putative causal signals in fine-mapping

With the confounding bias properly corrected, XMAP’s efficient algorithm allows us to produce reliable PIP for identifying multiple putative causal variants in thousands of loci across the whole genome. As summarized in Figure 6 A, with a PIP threshold of 0.5, XMAP identified 55 loci harboring more than one putative causal SNPs of height by combing UKBB and Chinese GWASs, among which 6 loci harbor more than 3 causal SNPs and 2 loci harbor 5 causal variants. With a stringent threshold PIP= 0.9, XMAP identified 15 loci with 2 causal SNPs and 9 loci with 3 causal SNPs. To examine the reliability of putative causal SNPs in loci harboring multiple causal signals, we evaluated the replication rates of these SNPs using the Sibship GWAS. Figure 6 B and C compare the replication rates of XMAP and SuSiE using their putative causal SNPs with a PIP threshold of 0.9. For loci with more than one putative causal SNPs, XMAP had the best replication rate (i.e., 24/55=43.6% SNPs had *p*-values*<* 10^−6^ and 14/55=25.5% SNPs had PIP *>* 0.1). Although SuSiE can also identify multiple causal signals (Supplementary Figure 17), it had lower replication rates than XMAP because it cannot correct for confounding bias. For loci with more than two putative causal SNPs, XMAP had similar replication rate with SuSiE applied to EUR GWAS. Although PAINTOR and MsCAVIAR can also integrate cross-population GWASs, they are too time-consuming to analyze all loci on the genome when the number of causal signals are set to be larger than 2. We could only run PAINTOR by setting the number of causal signals to 1 and 2. However, PAINTOR often produced unrealistic PIP for loci containing thousands of variants (Supplementary Figures 17 and 18). Here, we compared the PIP of SNPs computed by XMAP with PAINTOR and MsCAVIAR using the locus 130.2 Mbp-130.5Mbp in chromosome 6 as an example. We first combined the GWASs of UKBB (Figure 6 B) and Chinese (Figure 6 C). Clearly, all compared methods suggest that both rs1415701 and rs6569648 had high probability to be causal (Figure 6 E). To test the robustness of compared methods, we replaced the UKBB GWAS with EUR Sibship GWAS (Figure 6 D) which has smaller sample size but is less influenced by confounding bias, and computed the PIP again (Figure 6 F). Because of the reduced sample size, the PIP of rs6569648 computed by MsCAVIAR reduced to 0.78; the PIP computed by PAINTOR substantially differed from its previous output. By contrast, XMAP was the only method that consistently produced high PIP for rs1415701 and rs6569648 (PIP*>* 0.8).

**Figure 6:**
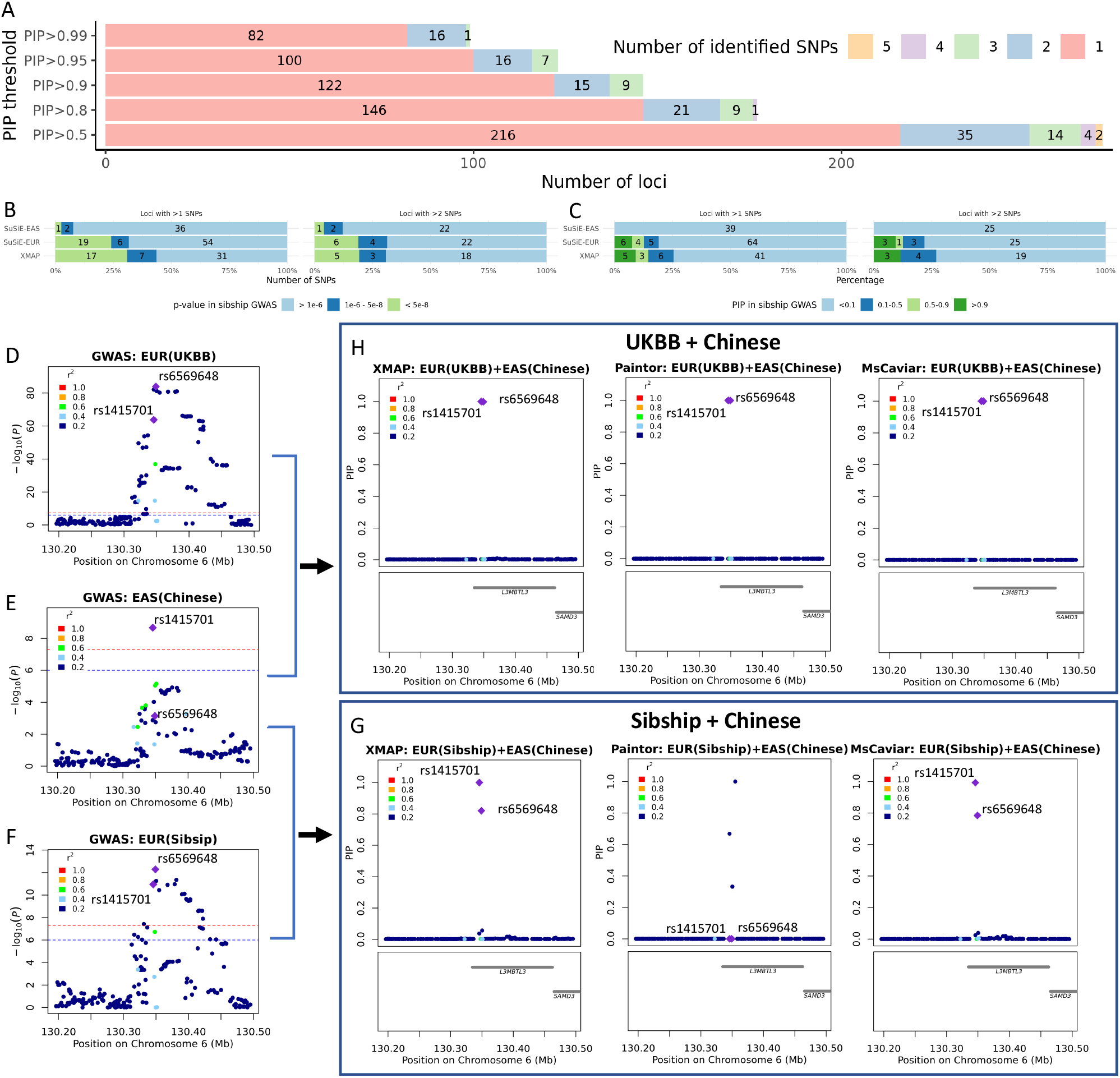
Performance of XMAP in identifying multiple causal variants for height. (A) Distributions of the number of putative causal SNPs identified by XMAP under different PIP thresholds. (B) With a PIP threshold of 0.9, the *p*-value distributions in the Sibship GWAS replication cohort are shown for putative causal SNPs within loci harboring *>* 1 and > 2 putative causal SNPs. (C) With a PIP threshold of 0.9, the PIP distributions in the Sibship GWAS replication cohort are shown for putative causal SNPs within loci harboring > 1 and > 2 putative causal SNPs. (D-G) A demonstrative example using the locus 130.2 Mbp-130.5Mbp in chromosome 6. Manhattan plots of the locus are shown for UKBB GWAS in (D), Chinese GWAS in (C), and EUR Sibship GWAS in (F). The PIP of SNPs in target locus are computed by XMAP, PAINTOR and MsCAVIAR with GWASs of UKBB+Chinese (H) and Sibship+Chinese (G).

In the main analysis, we set *K* = 10 to allow the detection of multiple causal variants. The setting *K* = 10 was supported by the analysis of height as summarized in Figure 6, where most loci had *<* 5 causal variants in height. To investigate the sensitivity of fine-mapping performance to the parameter *K*, we further considered *K* = 15 for XMAP and SuSiE. As shown in Supplementary Figure 15, the number of putative causal SNPs identified by XMAP are highly consistent under different settings of *K*. Besides, the fine-mapped SNPs could be replicated in a consistent rate under different settings of *K* (Figure 5 A-C and Supplementary Figure 16). These evidence consolidate our conclusion of the XMAP’s robustness to the setting of *K*.

### The XMAP output improves the interpretation of risk variants in their relevant cellular context at single-cell resolution

Integration of fine-mapping results with single-cell datasets is expected to offer a better interpretation of putative causal variants in their relevant cellular context at single-cell resolution [6]. However, fine-mapping of an under-presented population often lacks statistical power due to the limited sample size, making the interpretation of causal risk variants difficult. In this section, we show that cross-population fine-mapping results given by XMAP can greatly improve the interpretation of putative causal variants in their relevant cellular context by integrating single-cell datasets. To illustrate this benefit, we carried out SCAVENGE [6] analysis to quantify the enrichment of putative causal variants for 12 blood traits (summarized in Supplementary Table 1) within regions of accessible chromatin using the single-cell assay for transpose-accessible chromatin by sequencing (scATAC-seq). We employed a scATAC-seq dataset that encompasses multiple hematopoietic lineages [34], which includes 33,819 cells from 18 hematological populations (Figure 7 A). Specifically, we have a matrix of fragment counts **F** ∈ ℝ^*C*×*L*^, where *C* is the number of cells in scATAC-seq data and *L* is the number of accessible chromatin peaks. To quantify the relevance between the peaks and a phenotype, we first used the XMAP output to compute a vector of weight ***η*** ∈ ℝ^*L*^ with the *l*-th element of ***η*** being the sum of XMAP PIP for SNPs within the genomic region of peak *l*, which indicates the relative importance of a peak to the phenotype. The raw cell-trait relevance scores could be computed as **t** = **F*η***. As such, trait-related cells tend to have larger scores because more causal SNPs are located within their accessible chromatin regions. Then a *Z*-score characterizing the relationship between each pair of cell and trait can be obtained by further correcting for technical confounders, such as GC content bias and PCR amplification, using g-chromVAR [5]. To optimize the inference by leveraging relatedness across individual cells, we constructed a cell-cell similarity network and applied SCAVENGE [6] to assign a trait-relevance score (TRS) for each cell via network propagation. Finally, we simulated null distributions of TRS by using random seed cells for propagation and computed a *p*-value of trait-enrichment for each cell. The cells with *p*-value*<* 0.05 were considered as significantly enriched for the trait.

**Figure 7:**
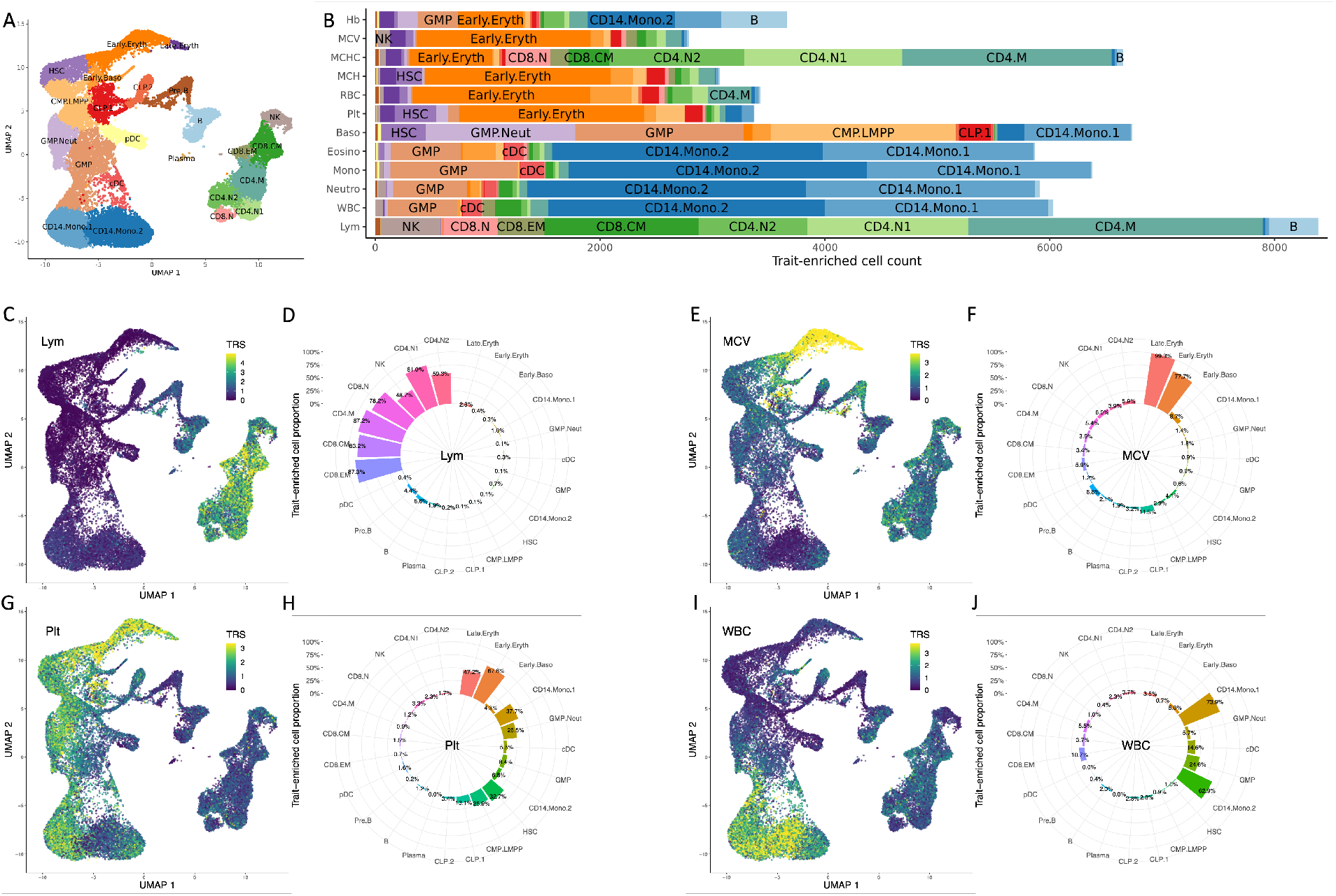
Enrichment of blood cell traits in hematological populations using XMAP fine-mapped SNPs as input. (A) The two dimensional uniform manifold approximation and projection (UMAP) plot of scATAC-seq data for 18 hematological populations. (B) The bar plots showing the number of cells significantly enriched in each of the 12 blood traits. The TRS are shown in the UMAP coordinates for four representative traits: Lym (C), MCV (E), Plt (G), and WBC (I). The proportions of significantly enriched cells within each population are shown for Lym (D), MCV (F), Plt (H), and WBC (J).

We summarized the identified trait-enriched cells and the median TRS of each cell type in Figure 7 B and Supplementary Figure 19, respectively. As we can observe, the enriched cells were highly aligned with our knowledge of cell types related to the blood traits. For example, we identified 8,388 lymphocyte count (Lym)-related cells, among which 5,021 cells were CD4 cells and 2,272 were CD8 cells. For traits related to myeloid/compound white cells, including eosinophil count (Eosino), monocyte count (Mono), neutrophil count (Neutro) and white blood cell count (WBC), we observed a substantial number of enriched cells from the CD14^+^ monocytes. For traits related to red cells such as red blood cell count (RBC), mean corpuscular hemoglobin (MCH), mean corpuscular hemoglobin concentration (MCHC), mean corpuscular volume (MCV), and hemoglobin (HB), a large amount of enriched cells were erythroid cells. These observations indicate that the biological mechanisms of putative causal SNPs identified by XMAP can be interpreted at single-cell resolution. Due to the unbalanced cell type composition in the single cell dataset, cells from rare populations can be under-represented. To rule out the influence of cell type composition on our analysis, we further investigated the proportion of trait-relevant cells within each cell type. We observed that biologically related cell types had largest proportion of enriched cells. For example, MCV was significantly enriched in 99.3% of late stage erythroid cells (Figure 7 E-F), WBC was significantly enriched in more than 60% of CD14^+^ monocytes (Figure 7 I-J), Lym was significantly enriched in a large proportion of CD4 and CD8 cells (Figure 7 C-D), and Plt was significantly enriched in the erythroid cells (Figure 7 G-H). These results suggest that the identification of trait-relevant cells is immune to the cell type composition. As shown in Supplementary Figures 20-31, we compared the trait-relevant cells obtained by using the XMAP PIP as input with those using the SuSiE PIP from single population analysis as input. Due to the relatively smaller sample size in the BBJ cohort, the trait-relevant cells were less enriched when fine-mapping was performed only using the BBJ GWASs, including GWASs of lymphocyte count (Supplementary Figure 20C-D), eosinophil count (Supplementary Figure 25C-D), and basophil count (Supplementary Figure 24C-D). Compared with the single-population fine-mapping result by SuSiE, XMAP can take the advantage of well-powered UKBB GWASs and provide a more accurate fine-mapping result (Supplementary Figures 20A-B, 25A-B, 24A-B). By integrating with single-cell datasets, the fine-mapping results given by XMAP can offer a better understanding of the putative causal variants in their cellular context at single-cell resolution.

## Discussion

In this paper, we have introduced a novel method named XMAP for cross-population fine-mapping. XMAP is able to improve the statistical power of fine-mapping by leveraging heterogeneous LD patterns across multiple populations. By correcting the hidden confounding bias in GWAS summary statistics, XMAP can effectively reduce spurious causal signals induced by sample structure. XMAP’s fast algorithm allows us to efficiently analyze loci that harbour multiple causal SNPs. Through comprehensive simulations, we showed that XMAP has greater statistical power, better control of false positive rate, and substantially higher computational efficiency for identifying multiple causal signals. We applied XMAP to fine-map causal SNPs of LDL by combining GWASs from EAS, EUR and AFR, achieving substantial gains in statistical power. Furthermore, we showed that XMAP was able to exclude spurious signals and produced reproducible results. By combining the output of XMAP for blood traits with scATAC-seq profiles of hematopoietic cells, we illustrated that the output of XMAP was particularly helpful to characterize the causal mechanism behind phenotypic variation at single-cell resolution. We believe that XMAP can serve as a powerful analytic tool of fine-mapping.

Considering the polygenic nature of complex traits, XMAP assumes that the genetic effects can be decomposed into two parts: the major causal effects and polygenic effects. For the causal effects, we assume that the total effects can be decomposed as a sum-of-single-effects [23, 24], which enables a highly efficient algorithm. While this assumption was also adopted in previous works [23, 24], they could not leverage genetic diversity to improve statistical power in the cross-population setting. For the polygenic effects, it benefits fine-mapping in two aspects. First, it captures the small genetic effects, allowing us to focus on the causal SNPs with major genetic impact that can be more biologically interesting for downstream analysis. Second, the statistical inference of causal effects are protected against over-fitting when *K* is specified larger than the ground truth. Therefore, we can safely set *K* to be a larger number, when the ground truth is unknown (Supplementary Figure 3). The parameters of the polygenic component are pre-estimated using LDSC, ensuring the model identifiability (see Methods). Because SNPs from the entire genome are used for estimation, the parameter estimates of the polygenic component are accurate and stable.

Identifying the tissue and cellular context of causal variants is a critical step to understand their biological mechanisms. Existing methods are usually limited to investigation at tissue [35, 36, 37, 38, 39, 40, 41, 42] or cell type levels [43, 44, 45, 46, 47], which do not fully utilize the rich resources of single-cell profiles. An important feature of XMAP is that it produces outputs that can be integrated with single-cell profiles to illuminate the cellular context of putative causal SNPs at single-cell resolution, offering a unique opportunity to characterize the biological mechanisms across a whole spectrum of cell functions.

Although it is convenient to work with GWAS summary statistics, fine-mapping requires a population-matched reference LD matrix as an input. The inconsistency of LD patterns between reference samples and GWAS samples can lead to false positive findings [24, 48, 49]. In our main analysis, we have used the in-sample LD references for EAS and UKBB GWAS to minimize the risk of LD mismatching. In practice, if an in-sample LD reference is not available, some diagnostic tools such as SLALOM [49] and DENTIST [48] should be carried out to validate the fine-mapping results and remove suspicious signals.

Our XMAP approach needs more investigation in the following directions. First, similar to PAINTOR and MsCAVAIR, XMAP assumes that the causal variants are shared across populations. Recent studies have reported that some causal signals could be specific to a certain population [50]. Hence, extending XMAP to handle the population-specific causal effects may yield biologically interesting discoveries. Second, causal variants are reported to be distributed disproportionately in the genome, depending on the functional context of the genomic regions [18, 25, 30, 51]. Some recent methods incorporate the information of functional annotation to improve fine-mapping [18, 25, 30]. It is interesting to incorporate functional annotations in the causal inference of XMAP, which may further boost the statistical power of fine-mapping. Third, gene-level effects can be more stably shared across populations, as compared to SNP-level effects. A recent study [52] suggests that the correlation of gene-level effects is 20% stronger than SNP-level effects across populations. Therefore, leveraging the genetic diversity at the gene-level for fine-mapping can be also an interesting direction. We will explore these potential extensions in the near future.

## Methods

### The XMAP model

We begin with the probabilistic formulation of XMAP with individual-level GWAS data. For easier introduction, we consider the case of two populations for easier introduction but note XMAP that is generally applicable to analyze multiple populations. Let {**y**_1_, **X**_1_} and {**y**_2_, **X**_2_} be the GWAS datasets collected from two different populations, where 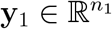 and 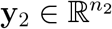 are phenotype vectors, 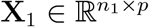 and 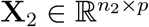 are genotype matrices, *p* is the number of SNPs in the locus of interest, and *n*_1_ and *n*_2_ are the GWAS sample sizes of populations 1 and 2, respectively. With different recombination rates, the two populations tend to have different LD patterns, i.e., the correlations among columns of **X**_1_ are usually distinct from those of **X**_2_. Without loss of generality, we assume that the columns of **X**_1_ and **X**_2_ have been standardized to have zero mean and unit variance. To relate genotypes and phenotypes, we consider the following linear models:

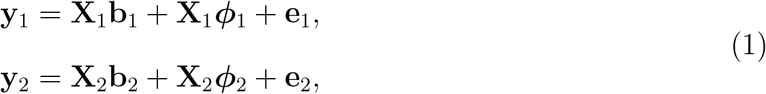

where **b**_1_ ∈ ℝ^*p*^ and **b**_2_ ∈ ℝ^*p*^ are sparse vectors of causal effects with major impact on phenotypes, ***ϕ***_1_ = [*ϕ*_11_, *ϕ*_12_, …, *ϕ*_1*p*_]^*T*^ ∈ ℝ^*p*^ and ***ϕ***_2_ = [*ϕ*_21_, *ϕ*_22_, …, *ϕ*_2*p*_]^*T*^ ∈ ℝ^*p*^ are dense vectors capturing the polygenic effects [53], and 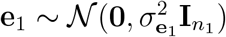 and 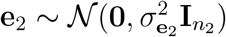 are vectors of independent noises from populations 1 and 2, respectively. Here, we assume that the covariates (e.g., sex, age, and principal components) have been adjusted. The detailed treatment of covariates follows our previous works [28, 54]. Unlike previous methods that only consider the overall genetic effects [17, 18, 19, 20, 22], we separate the genetic effects into causal and polygenic components. This decomposition allows us to focus on the causal SNPs with major genetic impact **b**_1_ and **b**_2_ that can be more biologically interesting for downstream analysis. Accumulating evidence of a shared genetic basis across populations [28, 25, 26, 55, 56] implies that **b**_1_ and **b**_2_ tend to have the same set of nonzero entries. Therefore, we expect that the different LD patterns in **X**_1_ and **X**_2_ can be helpful for fine-mapping shared causal SNPs across populations.

To leverage the cross-population GWASs for fine-mapping, we propose to specify model (1) by decomposing the causal genetic effects **b**_1_ and **b**_2_ into *K* ‘single effects’:

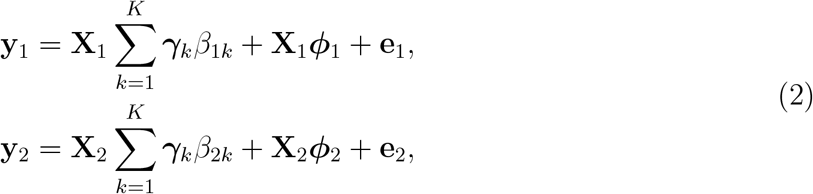

where *β*_1*k*_ and *β*_2*k*_ are effect sizes of the *k*-th causal signal in populations one and two, respectively, ***γ***_*k*_ = [*γ*_*k*1_, …, *γ*_*kp*_]^*T*^ ∈ {0, 1}^*p*^ in which only one element is 1 and the rest are 0 with *γ*_*kj*_ = 1 indicating the *j*-th variant is responsible for the *k*-th causal signal. This formulation of XMAP has three salient properties. First, through the shared causal status ***γ***_*k*_, XMAP can leverage the distinct LD patterns between **X**_1_ and **X**_2_. Meanwhile, we allow the two populations to have different effect sizes *β*_1*k*_ and *β*_2*k*_. Second, the decomposition of the causal signals into *K* single causal effects not only allows us to characterize each individual causal signal with an associated credible set [23] but also offers a computational advantage over existing methods, as we shall see later. Third, the inclusion of the polygenic component also protects the statistical inference against over-fitting when *K* is specified larger than the ground truth. With this property, we can safely set *K* to be a reasonably large number, say *K* = 10 by default, when the ground truth is unknown. To infer the causal status ***γ***_*k*_, we specify the probabilistic structures for the genetic effects in model (2) as follows:

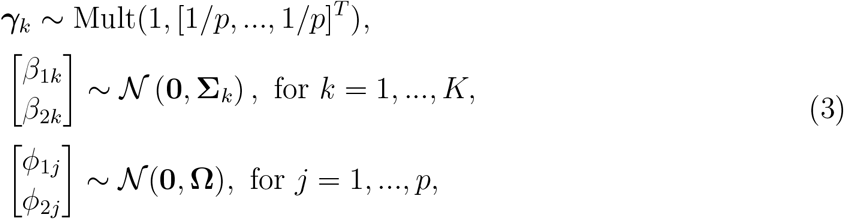

where Mult(1, [1*/p*, …, 1*/p*]^*T*^) denotes the non-informative categorical distribution of class counts drawn with class probabilities given by 1*/p* for each SNP, 𝒩 (**0, Σ**_*k*_) and 𝒩 (**0, Ω**) denote the multivariate normal distributions with mean **0** and covariance matrices 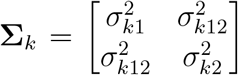 and 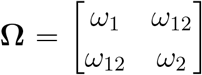, respectively. The variance components **Σ** = {**Σ**, …, **Σ**} capture the genetic covariance of the tow populations attributed to the *K* causal effects, and **Ω** captures the genetic covariance attributed to the polygenic effects.

So far, we have assumed the covariates have been adjusted. In the presence of covariates, we can extend XMAP model in Equation (2) as

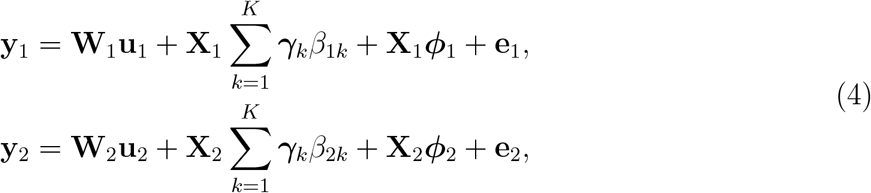

where 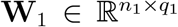 and 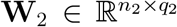 are the covariate matrices of populations 1 and 2, respectively, and 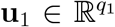 and 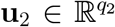 are corresponding vectors of covariate effects. To adjust the covariates, we first construct the projection matrices 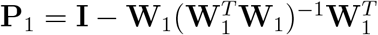 and 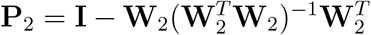. Then we multiply **P**_1_ on both sides of the first equation and **P**_2_ on both sides of the second equation in model (4). Through this projection, we can obtain a model without covariates

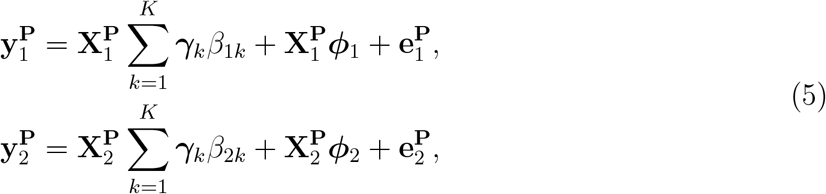

where 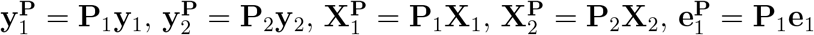, and 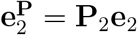. As we can observe, model (5) reduces to model (2). With this equivalence, we can work with model (2) without loss of generality.

### The XMAP model for summary-level data

Due to privacy concerns, the individual-level GWAS data may not be easily accessible. Given this situation, we consider the summary-level GWAS data 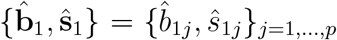 and 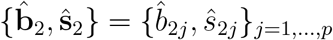 obtained from simple linear regressions:

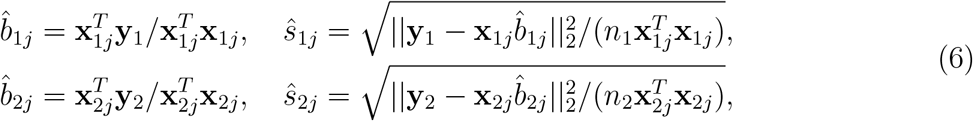

where **x**_1*j*_ ∈ ℝ^*p*^ and **x**_2*j*_ ∈ ℝ^*p*^ denote the *j*-th column of **X**_1_ and **X**_2_, respectively. To derive XMAP with summary-level data, we consider the rows of **X**_1_ and **X**_2_ as independently and identically distributed samples drawn from the two populations, respectively. Then, we define the LD matrices **R**_1_ = {*r*_1*jl*_} ∈ ℝ^*p*×*p*^ and **R**_2_ = {*r*_2*jl*_} ∈ ℝ^*p*×*p*^, where 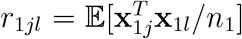 and 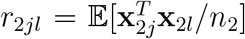 denote the correlation between variants *j* and *l* in populations 1 and 2, respectively. We can then obtain the expectation of GWAS effect sizes conditional on **b** and ***ϕ***:

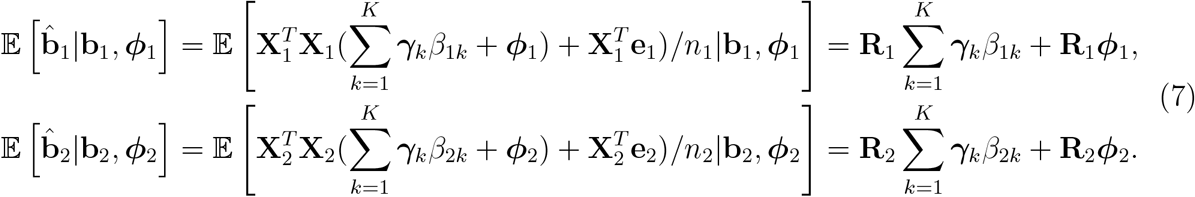

With this expression, we can connect **b** and ***ϕ*** with GWAS summary data with the following model:

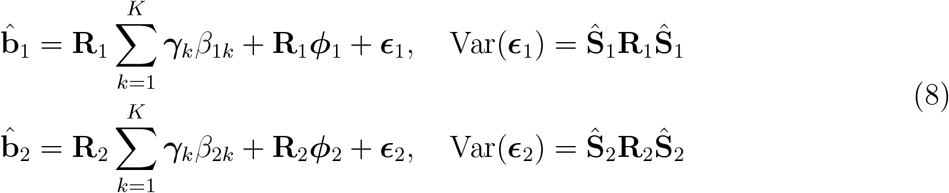

where **Ŝ**_1_ ∈ ℝ^*p*×*p*^ and **Ŝ**_2_ ∈ ℝ^*p*×*p*^ are diagonal matrices with diagonal terms given as { **Ŝ**_1_}_*jj*_ = *ŝ*_1*j*_ and { **Ŝ**_2_}_*jj*_ = *ŝ*_2*j*_ for *j* = 1, …, *p*, respectively (see Supplementary Note). To obtain a likelihood function of summary level data, we impose normal distributions for 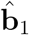 and 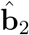, and Eq. (8) becomes the following model:

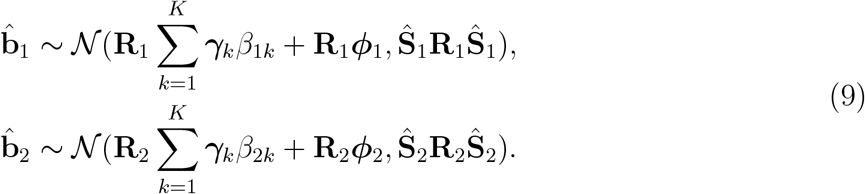

Note that model (8) or model (9) is derived by assuming that all the population structures have been properly adjusted in the GWAS summary statistics. To account for the unadjusted confounding bias hidden in GWAS summary statistics, we extend Equation (1) under the genetic drift model of LDSC [31] (see Supplementary Note). We show that model (9) is modified accordingly as

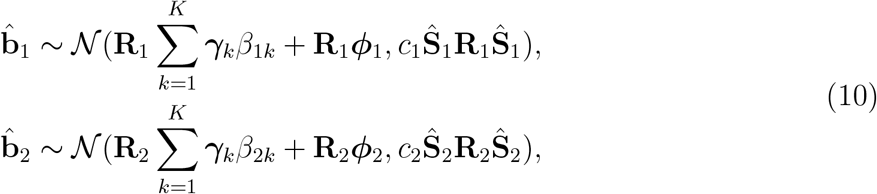

where *c*_1_ and *c*_2_ are LDSC intercepts that indicate the magnitude of inflation in GWAS effect sizes due to confounding bias. In the absence of confounding bias, the values of inflation constants *c*_1_ and *c*_2_ are close to one. As observed in biobank-scale GWASs [11, 12, 13, 54], the inflation constant is often larger than one in the presence of confounding bias. These inflation constants in the variance term of model (10) can re-calibrate the GWAS standard error based on the magnitude of confounding effects. The SNP correlation matrices **R** = {**R**_1_, **R**_2_} can be estimated with genotypes either from subsets of GWAS samples or from population-matched reference panels. Under model (3) and (10), we denote the collection of unknown parameters ***θ*** = {**Σ, Ω**, *c*_1_, *c*_2_}, and the collections of latent variables ***ϕ*** = {***ϕ***_1_, ***ϕ***_2_}, ***γ*** = {***γ***_*k*_}_*k*=1,…,*K*_ and ***β*** = {*β*_1*k*_, *β*_2*k*_}_*k*=1,…,*K*_. We shall obtain the parameter estimates 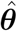 and identify causal SNPs with the posterior

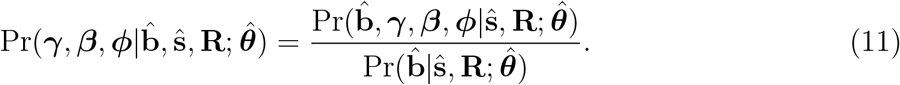

### Algorithm and parameter estimation

To ensure the model identifiability, we first apply LDSC to estimate the parameters *c*_1_, *c*_2_, and **Ω** using summary statistics across the whole genome. For **Ω**, the diagonal terms *ω*_1_ and *ω*_2_ are estimated with the per-SNP heritabilities of the corresponding populations using LDSC. The off-diagonal term *ω*_12_ is estimated by the per-SNP co-heritability obtained via bi-variate LDSC. The inflation constants *c*_1_ and *c*_2_ are estimated by the intercepts of LDSC of the two populations. Then, with the parameters 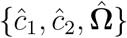 pre-fixed, we can estimate **Σ** without model identifiability issue. Traditional maximum likelihood approach estimates **Σ** by maximizing the marginal likelihood

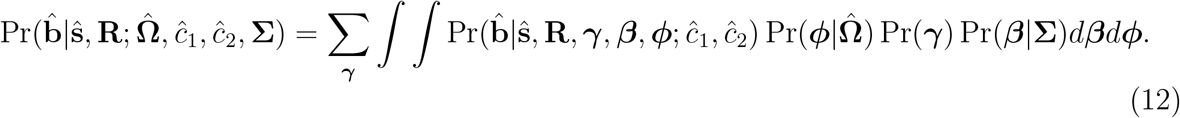

However, due to the combinatorial nature of ***γ***, the computational cost for Equation (12) grows exponentially with the number of causal signals *K*. To address this difficulty, we develop an efficient variational expectation-maximization (VEM) algorithm to estimate **Σ** and approximate the posterior (11). To achieve this, we first derive a lower bound of the logarithm of the marginal likelihood (12)

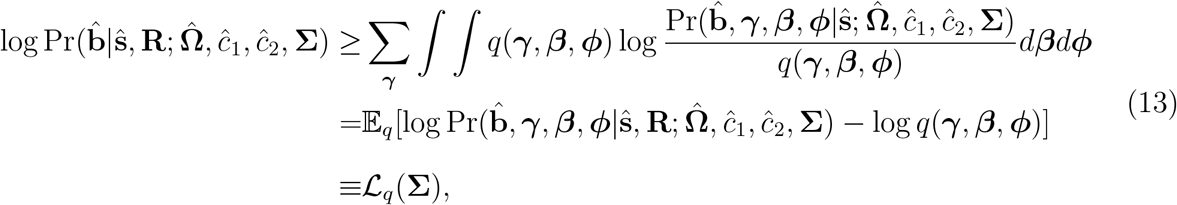

where the inequality follows Jensen’s inequality and *q*(***γ, β, ϕ***) is a variational approximation of the posterior (11). For convenience, we denote **b**_1*k*_ = ***γ***_*k*_*β*_1*k*_ and **b**_2*k*_ = ***γ***_*k*_*β*_2*k*_. By leveraging the decomposition in model (2), we propose a factorizable formulation of the variational approximation:

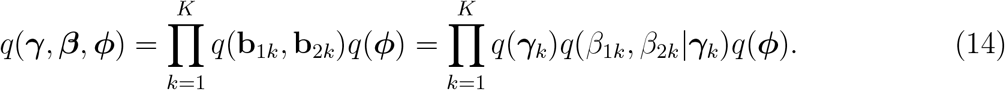

Unlike previous methods [57, 58] that require **b**_1*k*_ and **b**_2*k*_ to be fully factorizable across their *p* elements, the variational approximation in Equation (14) only requires that {**b**_11_, **b**_21_}, …, {**b**_1*K*_, **b**_2*K*_} are independent and they are independent of ***ϕ*** [23, 24], which allows flexible dependencies among the elements of **b**_1*k*_ and **b**_2*k*_. With the above factorizable approximation given by Equation (14), it turns out that both *q*(***γ, β, ϕ***) and ℒ_*q*_(**Σ**) can be analytically evaluated. We summarize the VEM algorithm in the following:

**E-step** At the *t*-th iteration, the variational distributions are given as

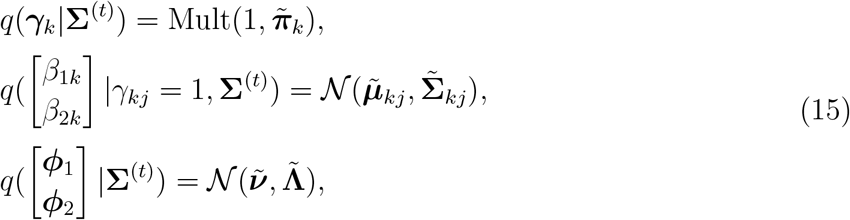

where 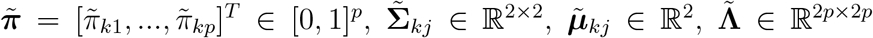, and 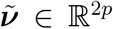 are variational parameters. The variational parameters are given as

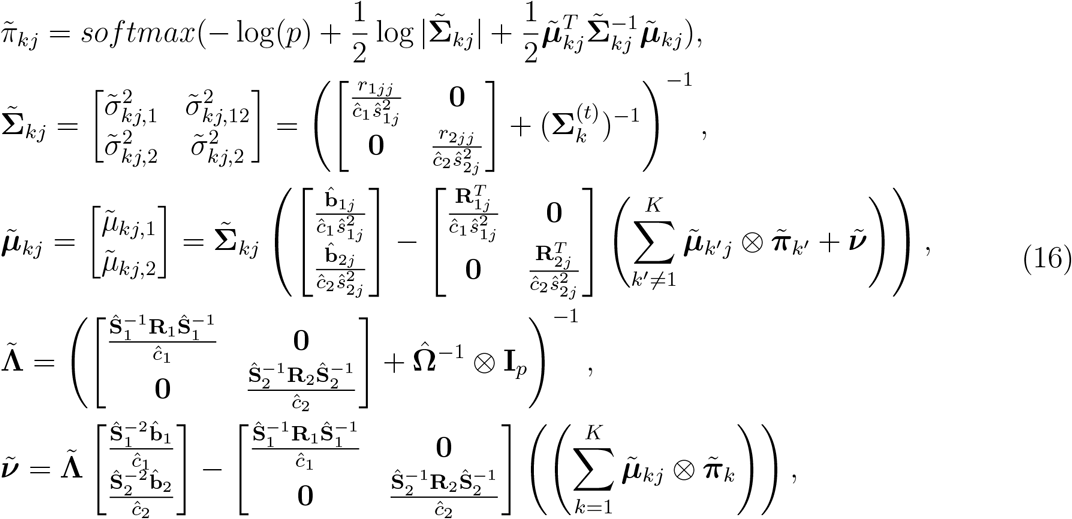

where *softmax* denotes the softmax function to make sure 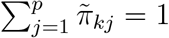 and ⊗ is the Kronecker product. By combing Equations (14,15,16), the lower bound (13) can be analytically evaluated as + constant,

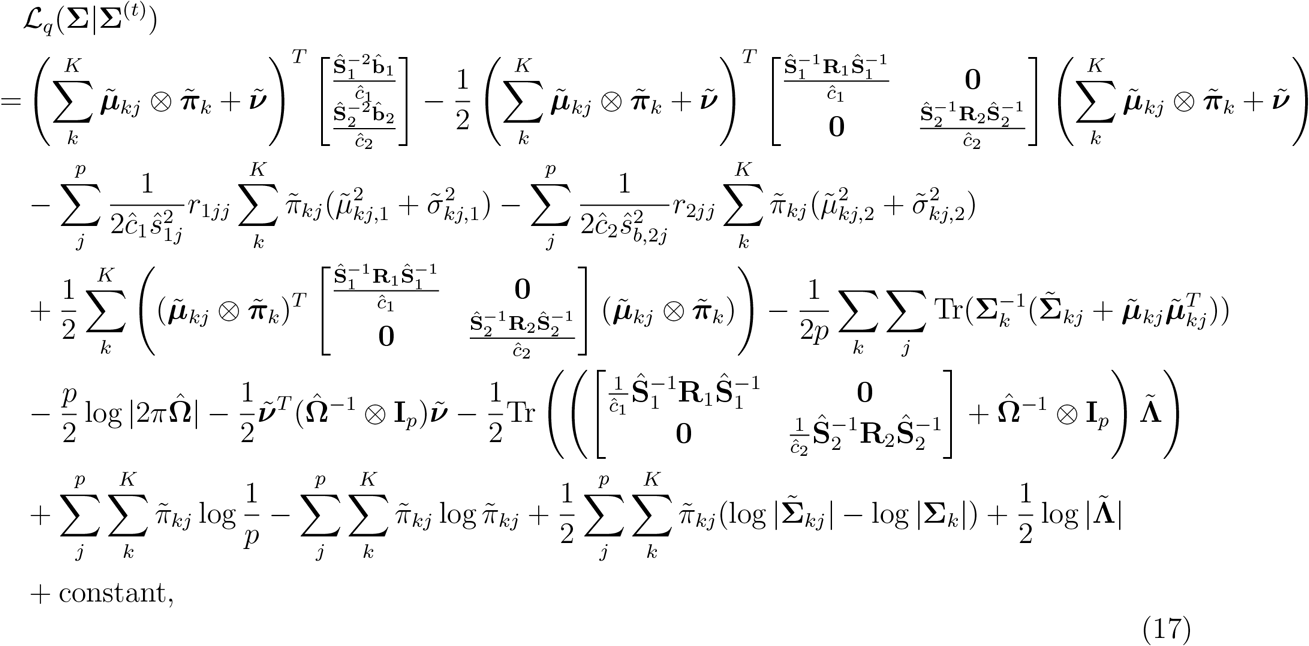

where Tr(**B**) denotes the trace of the square matrix **B**, the constant term does not involve **Σ. M-step** We solve 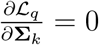 to obtain the update equation of **Σ**_*k*_:

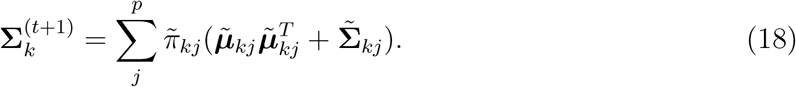

The above VEM algorithm has computational cost linear to the number of causal variants *K*, allowing for detecting multiple causal effects (e.g., *K* = 10) at a given locus.

### Identification of causal variant and construction of credible set

After the convergence of VEM algorithm, we can obtain the approximated posterior probabilities 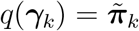, where 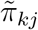 is the posterior probability that the *k*-th causal signal is contributed by the *j*-th SNP. With the variational approximation given by Equation (15), we can compute the posterior inclusion probability of SNP *j* as

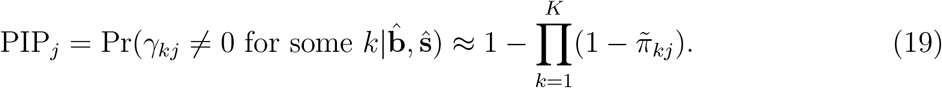

We can compute the local false discovery rate of SNP *j* as *fdr*_*j*_ = 1 − PIP_*j*_ and prioritize the causal SNPs by controlling the false discovery rate.

The decomposition of causal effects (2) offers an opportunity to characterize the set of SNPs that have high credibility to contribute to an individual causal signal. Let ℳ ⊂ {1, …, *p*} be a subset of SNPs from the target locus. A level-*α* credible set of a causal signal *k*, denoted as *CS*(*k, α*), is defined as the smallest M with 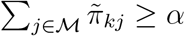. A smaller size of level-*α* credible set (e.g., *α* = 0.9) indicates a higher confidence of the identified causal variants.

### Influence and choice of *K*

The number of causal signals is usually unknown in practice. In XMAP, we do not require *K* to be the number of causal SNPs in the target locus. Instead, because the computational cost of our VEM algorithm only increases linearly with *K*, we can set *K* to a reasonably large number (e.g., *K* = 10) with minor computational overhead. When *K* is larger than the ground truth, the posterior probabilities in the excessive components will be broadly distributed across all SNPs in the locus because there is high uncertainty in the assignment of these causal effects. Importantly, the polygenic component will account for the small genetic effects, forcing the variance of excessive signals toward zero. Therefore, it has very minor influence in prioritization of causal SNPs when including extra causal effects than necessary. To exclude credible sets associated with redundant signal clusters, we follow SuSiE [23] to introduce the purity of credible sets. The purity of a credible set is defined as the smallest absolute correlation between pairs of SNPs within it. In XMAP, we consider the credible sets with purity less than 0.1 in all populations as redundant and discard the associated credible sets.

## Data and Code Availability

The publicly available GWAS summary statistics for meta-analysis were obtained from the links summarized in Supplementary Table 1. The XMAP software and source codes in this study were publicly available in GitHub repository of XMAP (https://github.com/YangLabHKUST/XMAP).

## Web resources

LDSC: https://github.com/bulik/ldsc;

XMAP: https://github.com/YangLabHKUST/XMAP;

PLINK: https://www.cog-genomics.org/plink;

BOLT-LMM: https://alkesgroup.broadinstitute.org/BOLT-LMM.

UKBB: https://www.ukbiobank.ac.uk;

SuSiE: https://github.com/stephenslab/susieR

PAINTOR: https://github.com/gkichaev/PAINTOR_V3.0

MsCAVIAR: https://github.com/nlapier2/MsCAVIAR

FINEMAP: http://www.christianbenner.com

DAP-G: https://github.com/xqwen/dap

g-chromVAR: https://github.com/caleblareau/gchromVAR

SCAVENGE: https://github.com/sankaranlab/SCAVENGE

