## Supplementary Figures and Note for "XMAP: Cross-population fine-mapping by leveraging genetic diversity and accounting for confounding bias"

### 1 Supplementary Tables

| | Cohort | Intercept | Intercept s.e. | per-SNP h2 | per-SNP h2 s.e. |
|---|---|---|---|---|---|
| **LDL-AFR** | GLGC | 1.066208 | 1.84E-02 | 3.88E-08 | 1.25E-08 |
| **LDL-EUR** | GLGC | 1.101591 | 1.47E-02 | 2.23E-08 | 5.97E-09 |
| **LDL-EUR** | UKBB | 1.094689 | 1.39E-02 | 1.93E-08 | 4.51E-09 |
| **LDL-EAS** | GLGC | 1.036577 | 7.17E-03 | 2.37E-08 | 4.50E-09 |
| **height-EUR** | UKBB | 1.658928 | 4.16E-02 | 7.60E-08 | 4.34E-09 |
| **height-EUR** | Sibship | 1.116069 | 1.17E-02 | 5.41E-08 | 3.36E-09 |
| **height-EAS** | Chinese | 1.069005 | 8.81E-03 | 8.45E-08 | 6.33E-09 |
| **height-EAS** | BBJ | 1.394916 | 2.41E-02 | 7.81E-08 | 5.31E-09 |
| **Lym-EUR** | UKBB | 1.236483 | 1.66E-02 | 3.00E-08 | 2.37E-09 |
| **Lym-EAS** | BBJ | 1.055371 | 8.18E-03 | 1.47E-08 | 1.96E-09 |
| **WBC-EUR** | UKBB | 1.250661 | 1.79E-02 | 2.85E-08 | 2.07E-09 |
| **WBC-EAS** | BBJ | 1.092382 | 8.88E-03 | 1.76E-08 | 1.93E-09 |
| **Neutro-EUR** | UKBB | 1.217628 | 1.55E-02 | 2.49E-08 | 2.33E-09 |
| **Neutro-EAS** | BBJ | 1.049925 | 7.98E-03 | 2.05E-08 | 2.91E-09 |
| **Mono-EUR** | UKBB | 1.240563 | 1.95E-02 | 3.40E-08 | 4.04E-09 |
| **Mono-EAS** | BBJ | 1.075889 | 8.16E-03 | 1.61E-08 | 2.71E-09 |
| **Eosino-EUR** | UKBB | 1.208816 | 1.78E-02 | 2.52E-08 | 2.37E-09 |
| **Eosino-EAS** | BBJ | 1.067848 | 8.74E-03 | 1.52E-08 | 2.77E-09 |
| **Baso-EUR** | UKBB | 1.113947 | 7.09E-03 | 3.21E-09 | 3.30E-10 |
| **Baso-EAS** | BBJ | 1.100899 | 9.22E-03 | 9.61E-09 | 3.77E-09 |
| **Plt-EUR** | UKBB | 1.27035 | 1.87E-02 | 4.62E-08 | 4.17E-09 |
| **Plt-EAS** | BBJ | 1.120054 | 9.69E-03 | 3.02E-08 | 4.30E-09 |
| **RBC-EUR** | UKBB | 1.257341 | 1.84E-02 | 3.53E-08 | 3.64E-09 |
| **RBC-EAS** | BBJ | 1.098981 | 9.41E-03 | 2.24E-08 | 3.19E-09 |
| **MCH-EUR** | UKBB | 1.176256 | 1.62E-02 | 4.62E-08 | 7.09E-09 |
| **MCH-EAS** | BBJ | 1.102186 | 9.43E-03 | 3.26E-08 | 5.78E-09 |
| **MCHC-EUR** | UKBB | 1.064449 | 9.01E-03 | 9.36E-09 | 1.20E-09 |
| **MCHC-EAS** | BBJ | 1.076843 | 8.91E-03 | 9.70E-09 | 2.09E-09 |
| **MCV-EUR** | UKBB | 1.202512 | 1.67E-02 | 4.60E-08 | 6.46E-09 |
| **MCV-EAS** | BBJ | 1.119433 | 1.03E-02 | 3.56E-08 | 5.76E-09 |
| **Hb-EUR** | UKBB | 1.21788 | 1.86E-02 | 3.32E-08 | 2.90E-09 |
| **Hb-EAS** | BBJ | 1.065907 | 8.12E-03 | 1.23E-08 | 1.45E-09 |

**Table 1:** Estimates of LDSC intercepts and per-SNP heritabilities.

| | Cohort | n | p | Publication | Source |
|---|---|---|---|---|---|
| **LDL-AFR** | GLGC | 92,934 | 25,476,275 | doi.org/10.1038/s41586-021-04064-3 | http://csg.sph.umich.edu/willer/public/glgc-lipids2021/results/ |
| **LDL-EUR** | GLGC | 664,450 | 35,328,891 | doi.org/10.1038/s41586-021-04064-3 | http://csg.sph.umich.edu/willer/public/glgc-lipids2021/results/ |
| **LDL-EUR** | UKBB | 343,621 | 12,515,778 | doi.org/10.1038/s41586-018-0579-z | nealelab.github.io/UKBB_ldsc/index.html |
| **LDL-EAS** | GLGC | 71,150 | 11,569,928 | doi.org/10.1038/s41586-021-04064-3 | http://csg.sph.umich.edu/willer/public/glgc-lipids2021/results/ |
| **height-EUR** | UKBB | 360,388 | 12,515,778 | doi.org/10.1038/s41586-018-0579-z | nealelab.github.io/UKBB_ldsc/index.html |
| **height-EUR** | Sibship | 71,872 | 6,101,836 | doi.org/10.1038/s41588-022-01062-7 | gwas.mrcieu.ac.uk/datasets/ieu-b-4813/ |
| **height-EAS** | Chinese | 32,921 | 3,776,576 | doi.org/10.1016/j.ajhg.2021.03.002 | doi.org/10.1016/j.ajhg.2021.03.002 |
| **height-EAS** | BBJ | 159,095 | 6,310,855 | doi.org/10.1038/s41467-019-12276-5 | http://jenger.riken.jp/en/result |
| **Lym-EUR** | UKBB | 349,856 | 12,515,778 | doi.org/10.1038/s41586-018-0579-z | nealelab.github.io/UKBB_ldsc/index.html |
| **Lym-EAS** | BBJ | 62,076 | 5,961,105 | doi.org/10.1038/s41588-018-0047-6 | http://jenger.riken.jp/en/result |
| **WBC-EUR** | UKBB | 350,470 | 12,515,778 | doi.org/10.1038/s41586-018-0579-z | nealelab.github.io/UKBB_ldsc/index.html |
| **WBC-EAS** | BBJ | 107,964 | 5,961,105 | doi.org/10.1038/s41588-018-0047-6 | http://jenger.riken.jp/en/result |
| **Neutro-EUR** | UKBB | 349,856 | 12,515,778 | doi.org/10.1038/s41586-018-0579-z | nealelab.github.io/UKBB_ldsc/index.html |
| **Neutro-EAS** | BBJ | 62,076 | 5,961,105 | doi.org/10.1038/s41588-018-0047-6 | http://jenger.riken.jp/en/result |
| **Mono-EUR** | UKBB | 349,856 | 12,515,778 | doi.org/10.1038/s41586-018-0579-z | nealelab.github.io/UKBB_ldsc/index.html |
| **Mono-EAS** | BBJ | 62,076 | 5,961,105 | doi.org/10.1038/s41588-018-0047-6 | http://jenger.riken.jp/en/result |
| **Eosino-EUR** | UKBB | 349,856 | 12,515,778 | doi.org/10.1038/s41586-018-0579-z | nealelab.github.io/UKBB_ldsc/index.html |
| **Eosino-EAS** | BBJ | 62,076 | 5,961,105 | doi.org/10.1038/s41588-018-0047-6 | http://jenger.riken.jp/en/result |
| **Baso-EUR** | UKBB | 349,856 | 12,515,778 | doi.org/10.1038/s41586-018-0579-z | nealelab.github.io/UKBB_ldsc/index.html |
| **Baso-EAS** | BBJ | 62,076 | 5,961,105 | doi.org/10.1038/s41588-018-0047-6 | http://jenger.riken.jp/en/result |
| **Plt-EUR** | UKBB | 350,474 | 12,515,778 | doi.org/10.1038/s41586-018-0579-z | nealelab.github.io/UKBB_ldsc/index.html |
| **Plt-EAS** | BBJ | 108,208 | 5,961,105 | doi.org/10.1038/s41588-018-0047-6 | http://jenger.riken.jp/en/result |
| **RBC-EUR** | UKBB | 350,475 | 12,515,778 | doi.org/10.1038/s41586-018-0579-z | nealelab.github.io/UKBB_ldsc/index.html |
| **RBC-EAS** | BBJ | 108,794 | 5,961,105 | doi.org/10.1038/s41588-018-0047-6 | http://jenger.riken.jp/en/result |
| **MCH-EUR** | UKBB | 350,472 | 12,515,778 | doi.org/10.1038/s41586-018-0579-z | nealelab.github.io/UKBB_ldsc/index.html |
| **MCH-EAS** | BBJ | 108,054 | 5,961,105 | doi.org/10.1038/s41588-018-0047-6 | http://jenger.riken.jp/en/result |
| **MCHC-EUR** | UKBB | 350,468 | 12,515,778 | doi.org/10.1038/s41586-018-0579-z | nealelab.github.io/UKBB_ldsc/index.html |
| **MCHC-EAS** | BBJ | 108,728 | 5,961,105 | doi.org/10.1038/s41588-018-0047-6 | http://jenger.riken.jp/en/result |
| **MCV-EUR** | UKBB | 350,473 | 12,515,778 | doi.org/10.1038/s41586-018-0579-z | nealelab.github.io/UKBB_ldsc/index.html |
| **MCV-EAS** | BBJ | 108,256 | 5,961,105 | doi.org/10.1038/s41588-018-0047-6 | http://jenger.riken.jp/en/result |
| **Hb-EUR** | UKBB | 344,182 | 12,515,778 | doi.org/10.1038/s41586-018-0579-z | nealelab.github.io/UKBB_ldsc/index.html |
| **Hb-EAS** | BBJ | 108,769 | 5,961,105 | doi.org/10.1038/s41588-018-0047-6 | http://jenger.riken.jp/en/result |

**Table 2:** GWAS sources

### 2 Supplementary Figures

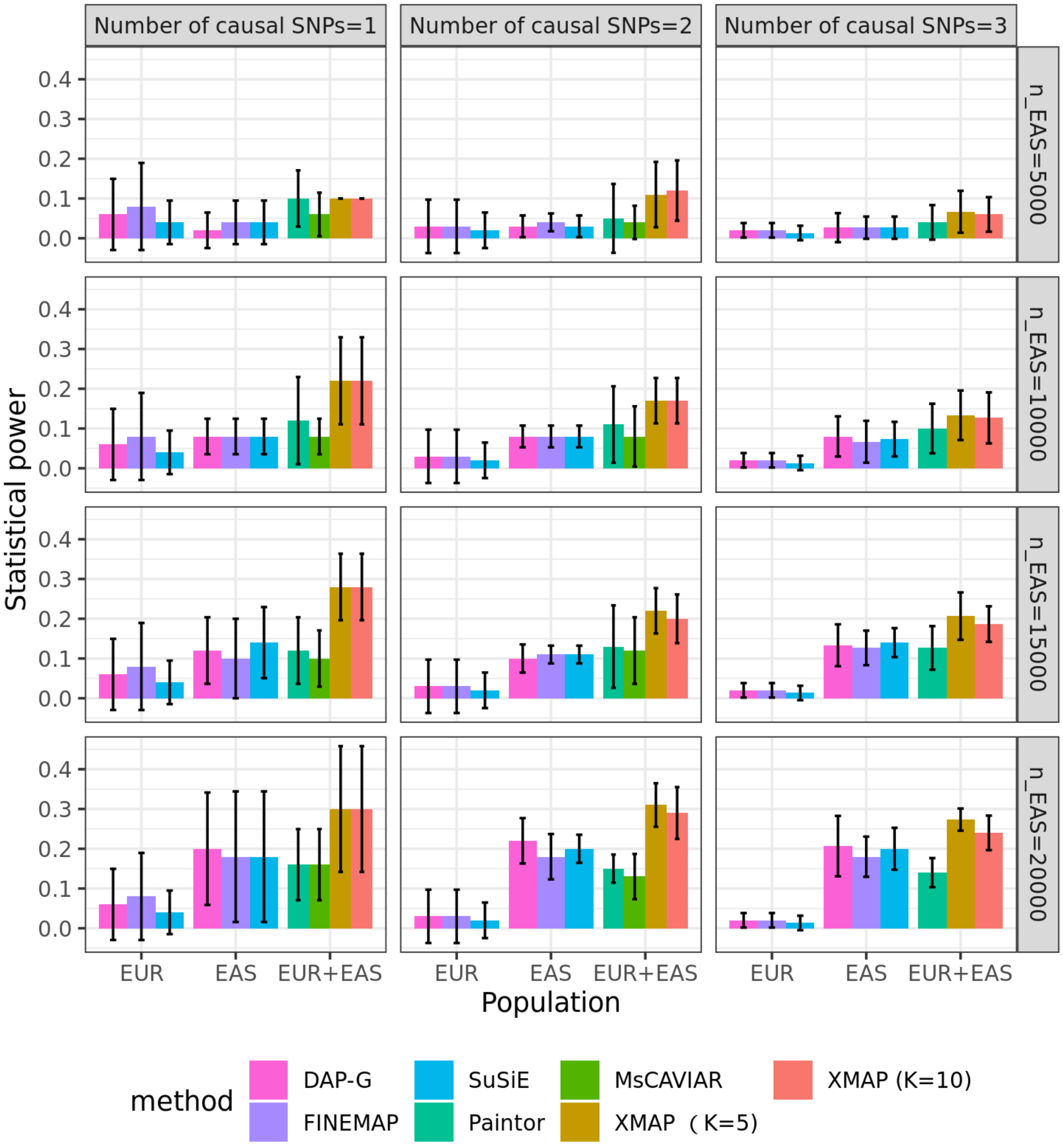

**Supplementary Figure 1:** Comparison of statistical power among DAP-G, FINEMAP, SuSiE, Paintor, MsCAIVAR, and XMAP. We varied $K_{true} \in \{1, 2, 3\}$ and EAS sample size $n_1 \in \{5,000,\ 10,000.\ 15,000,\ 20,000\}$, and set EUR sample size $n_2 = 20,000$. Because MsCAVIAR was intractable when including more then three causal signals, it was excluded from the comparison in the setting of $K_{true} = 3$. Error bars represent the standard errors of statistical powers evaluated on 50 replications.

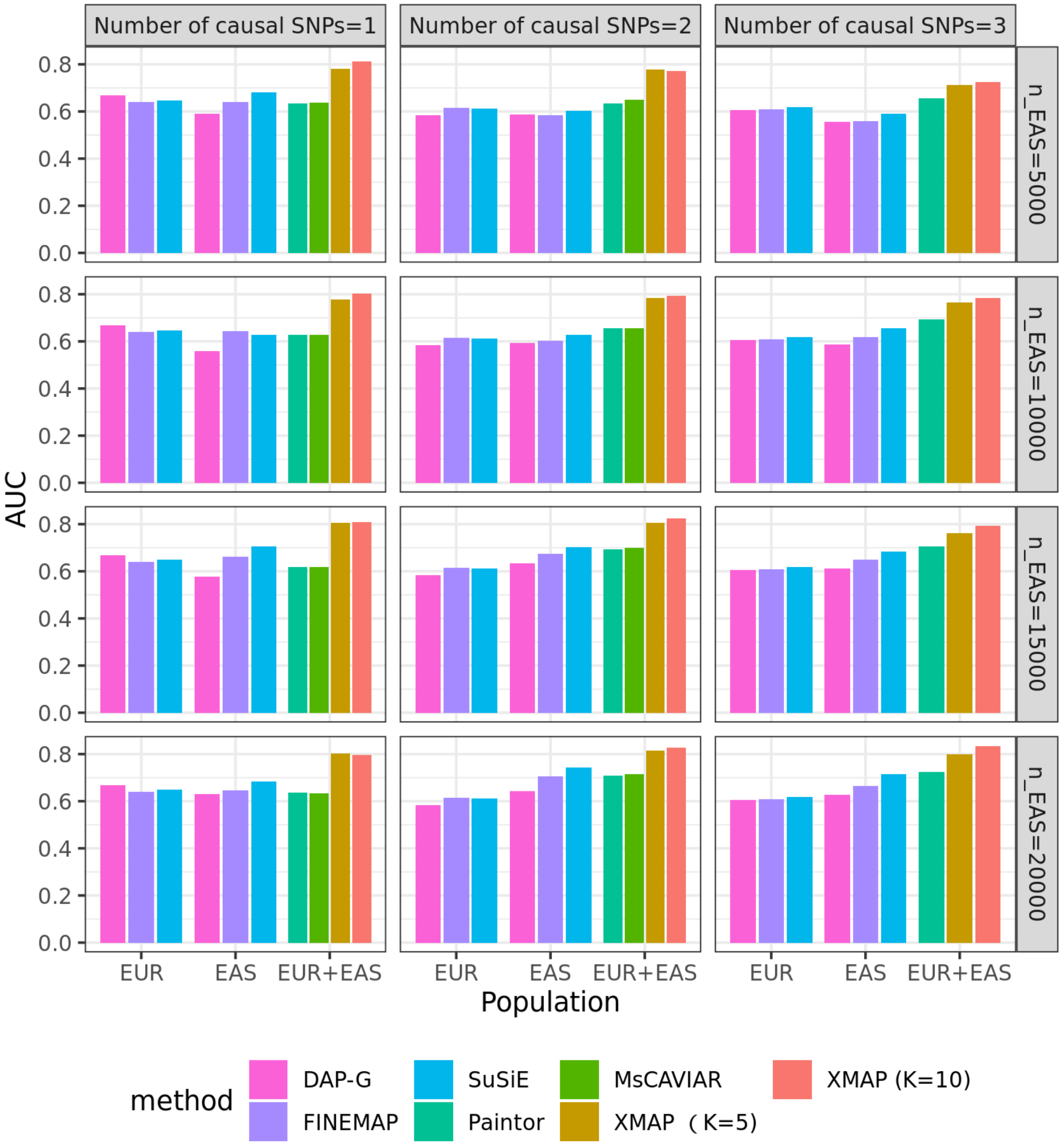

**Supplementary Figure 2:** Comparison of AUC among DAP-G, FINEMAP, SuSiE, Paintor, MsCAIVAR, and XMAP across 50 simulations. We varied $K_{true} \in \{1, 2, 3\}$ and EAS sample size $n_1 \in \{5,000,\ 10,000.\ 15,000,\ 20,000\}$, and set EUR sample size $n_2 = 20,000$. Because MsCAVIAR was intractable when including more then three causal signals, it was excluded from the comparison in the setting of $K_{true} = 3$.

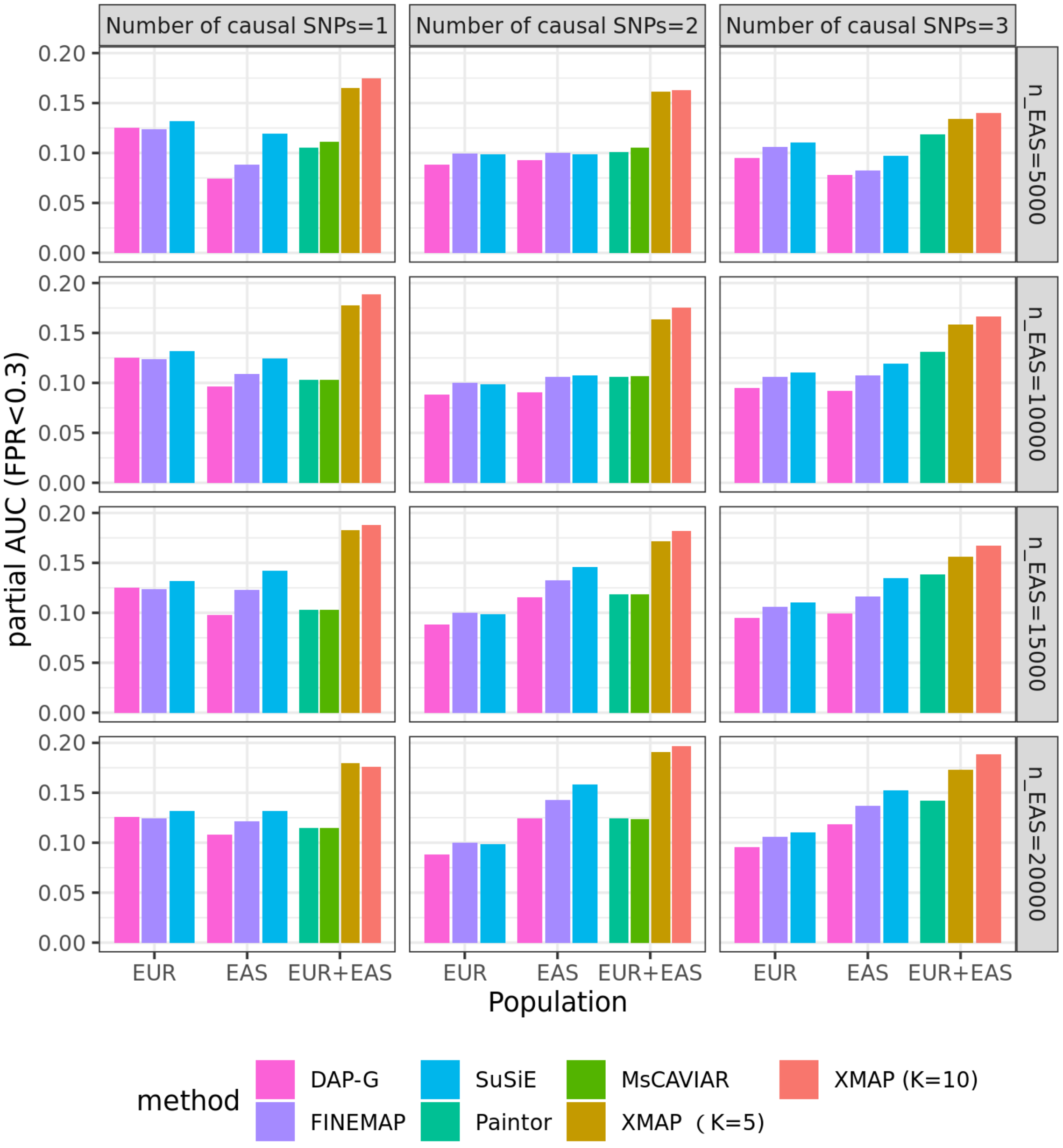

**Supplementary Figure 3:** Comparison of pAUC (FPR$<$ 0.3) among DAP-G, FINEMAP, SuSiE, Paintor, MsCAIVAR, and XMAP across 50 simulations. We varied $K_{true} \in \{1, 2, 3\}$ and EAS sample size $n_1 \in \{5,000,\ 10,000.\ 15,000,\ 20,000\}$, and set EUR sample size $n_2 = 20,000$. Because MsCAVIAR was intractable when including more then three causal signals, it was excluded from the comparison in the setting of $K_{true} = 3$.

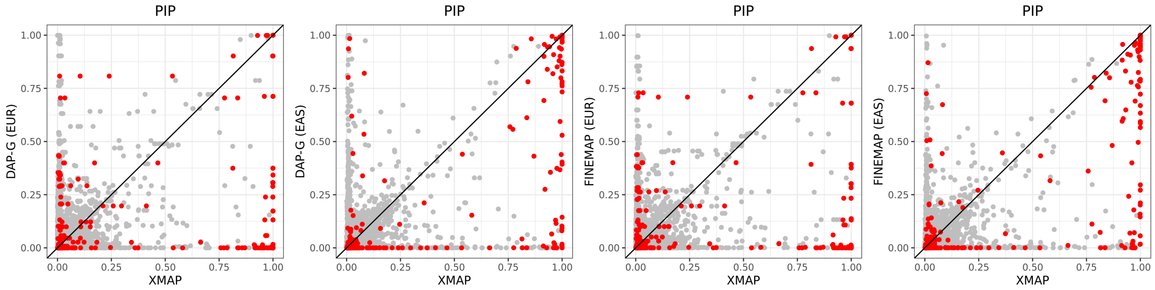

**Supplementary Figure 4:** Pairwise comparisons of PIP obtained by XMAP with those obtained by DAP-G and FINEMAP when $K_{true} = 3$.

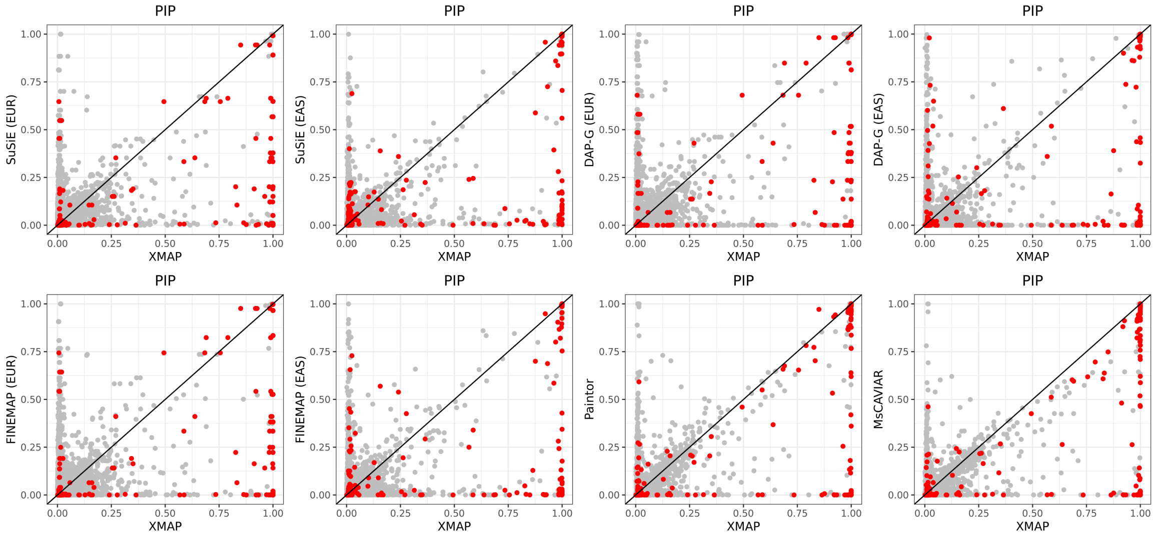

**Supplementary Figure 5:** Pairwise comparisons of PIP obtained by XMAP with those obtained by SuSiE, DAP-G, FINEMAP, PAINTOR and MsCAVIAR when $K_{true} = 2$.

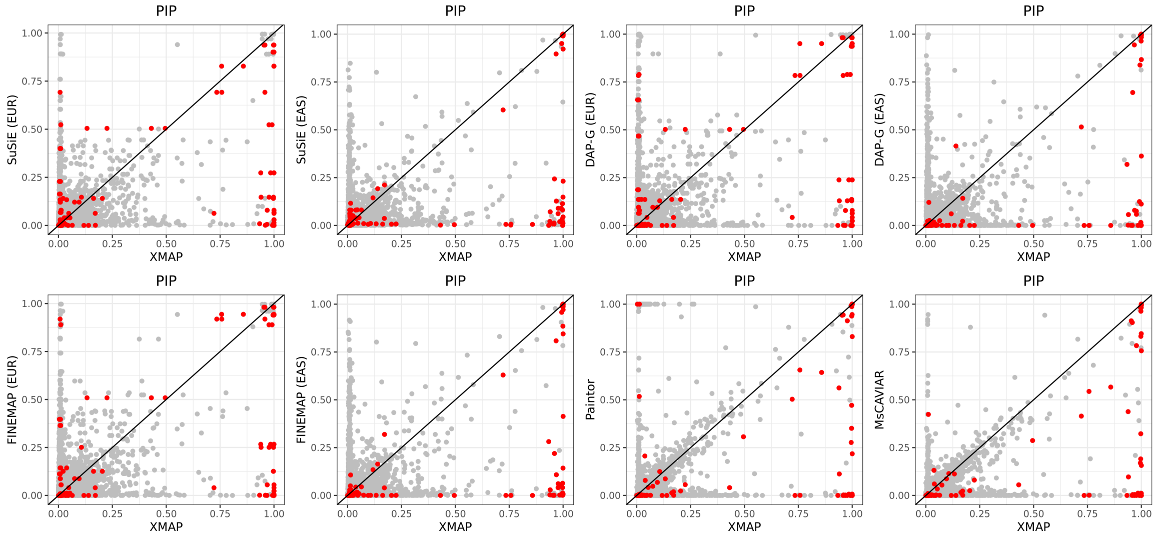

**Supplementary Figure 6:** Pairwise comparisons of PIP obtained by XMAP with those obtained by SuSiE, DAP-G, FINEMAP, PAINTOR and MsCAVIAR when $K_{true} = 1$.

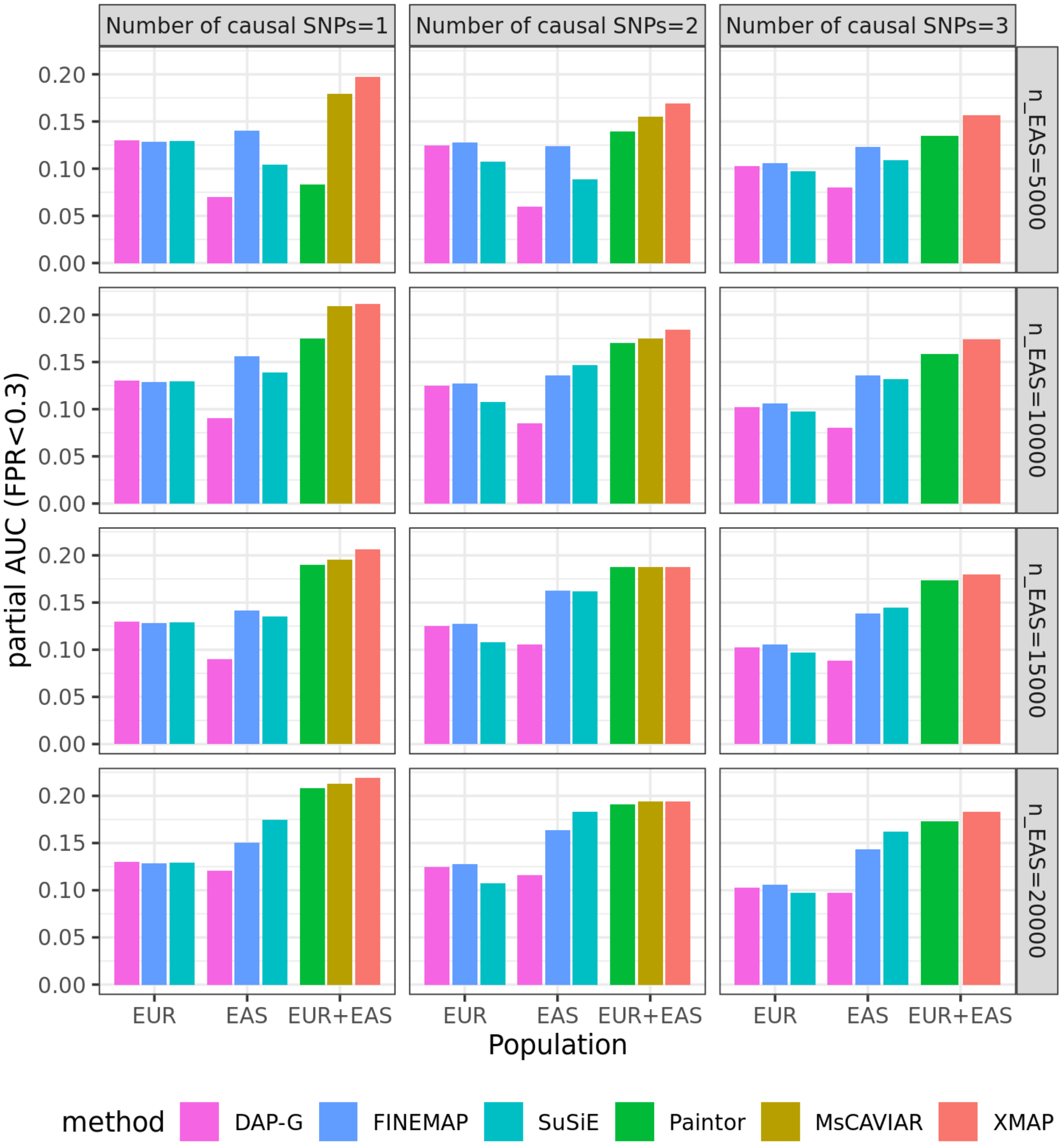

**Supplementary Figure 7:** Comparison of pAUC (FPR< 0.3) among DAP-G, FINEMAP, SuSiE, Paintor, MsCAIVAR, and XMAP in the presence of confounding bias across 50 simulations. We varied $K_{true} \in \{1, 2, 3\}$ and EAS sample size $n_1 \in \{5,000,\ 10,000.\ 15,000,\ 20,000\}$, and set EUR sample size $n_2 = 20,000$. Because MsCAVIAR was intractable when including more then three causal signals, it was excluded from the comparison in the setting of $K_{true} = 3$.

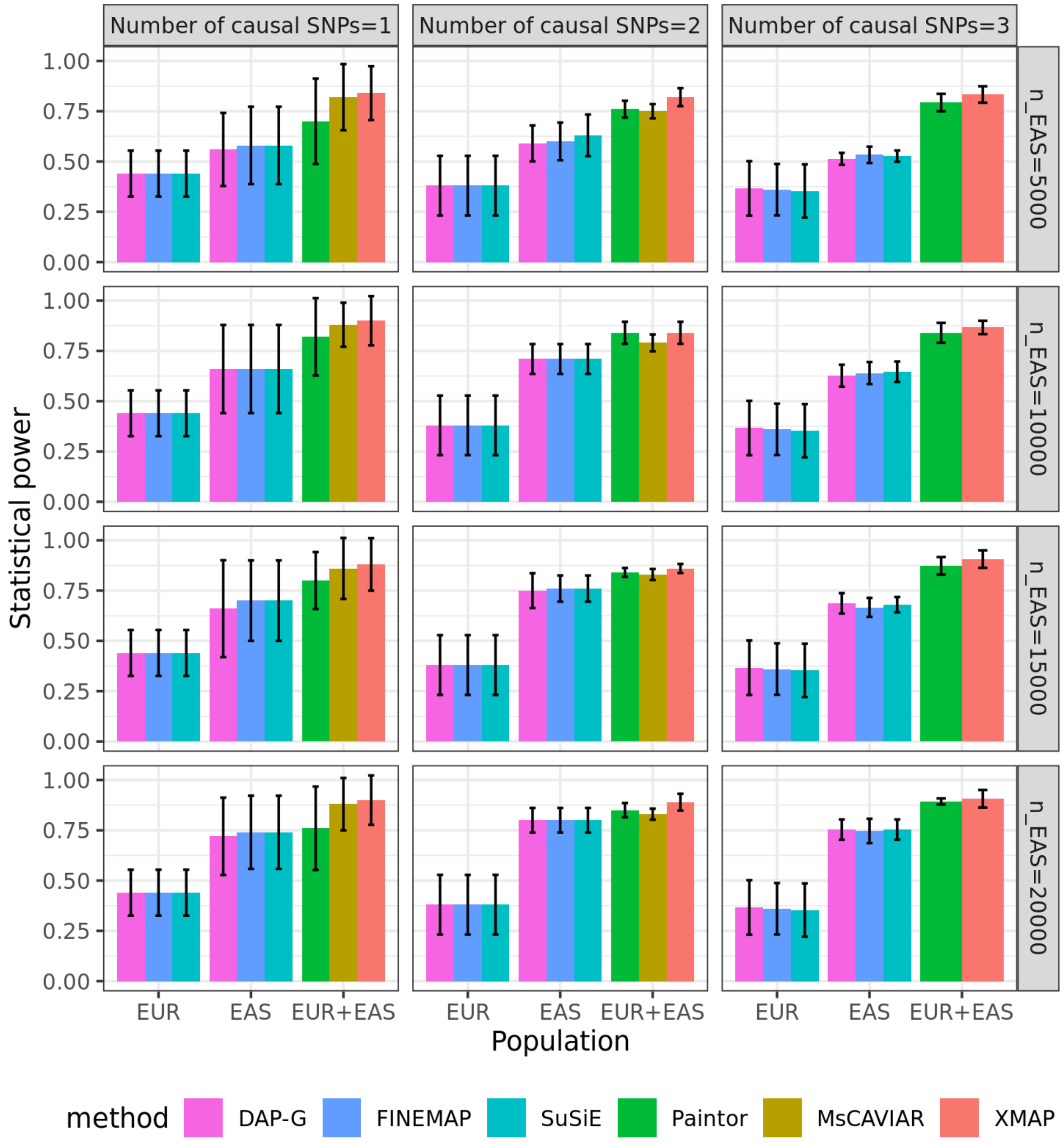

**Supplementary Figure 8:** Comparison of statistical power among DAP-G, FINEMAP, SuSiE, Paintor, MsCAIVAR, and XMAP in the setting without polygenic effects. We varied $K_{true} \in \{1, 2, 3\}$ and EAS sample size $n_1 \in \{5,000,\ 10,000.\ 15,000,\ 20,000\}$, and set EUR sample size $n_2 = 20,000$. Each of the $K_{true}$ causal SNPs explains 1% phenotypic variance. Because MsCAVIAR was intractable when including more then three causal signals, it was excluded from the comparison in the setting of $K_{true} = 3$. Error bars represent the standard errors of statistical powers evaluated on 50 replications.

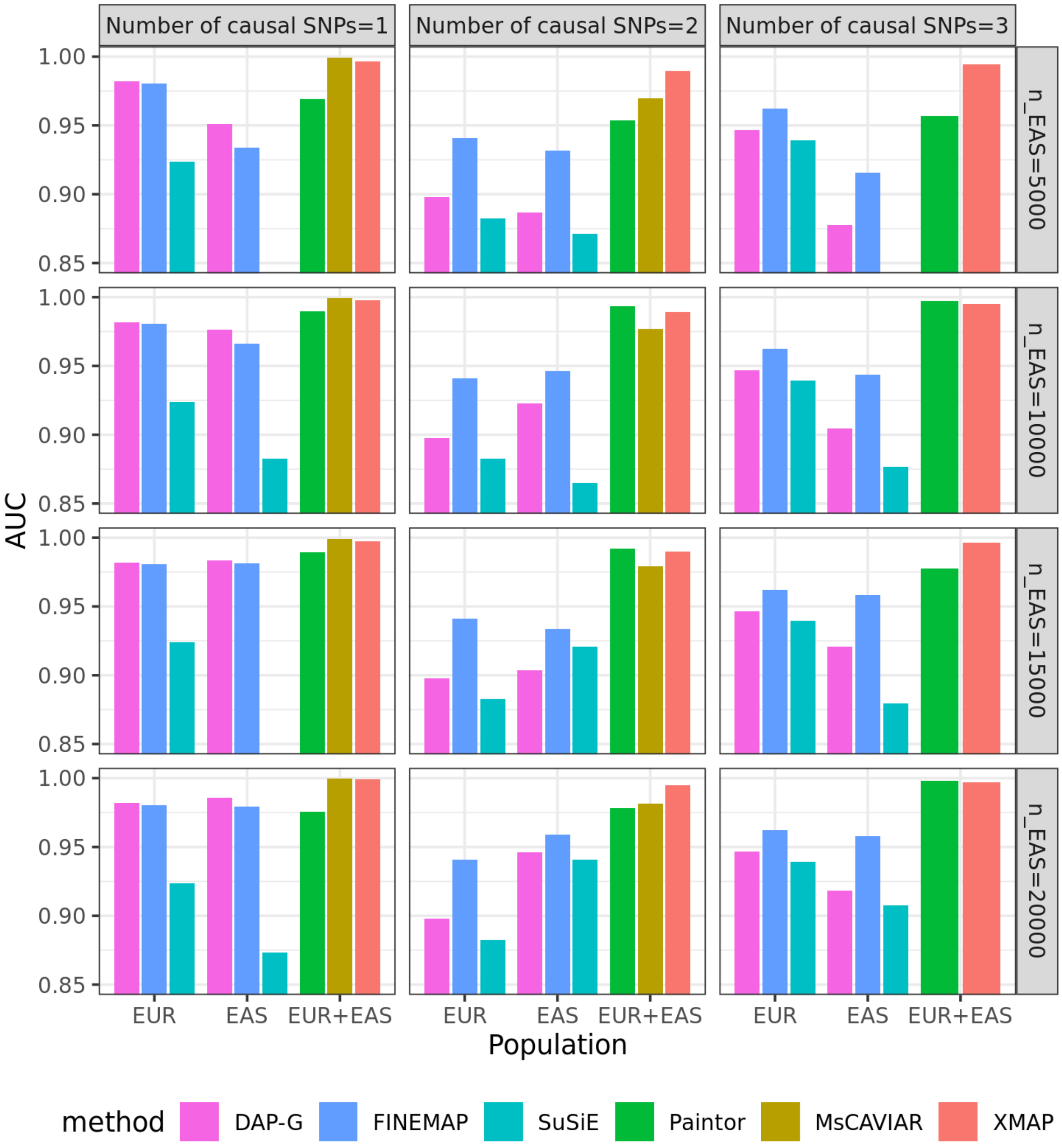

**Supplementary Figure 9:** Comparison of AUC among DAP-G, FINEMAP, SuSiE, Paintor, MsCAIVAR, and XMAP across 50 simulations in the setting without polygenic effects. We varied $K_{true} \in \{1, 2, 3\}$ and EAS sample size $n_1 \in \{5,000,\ 10,000.\ 15,000,\ 20,000\}$, and set EUR sample size $n_2 = 20,000$. Each of the $K_{true}$ causal SNPs explains 1% phenotypic variance. Because MsCAVIAR was intractable when including more then three causal signals, it was excluded from the comparison in the setting of $K_{true} = 3$.

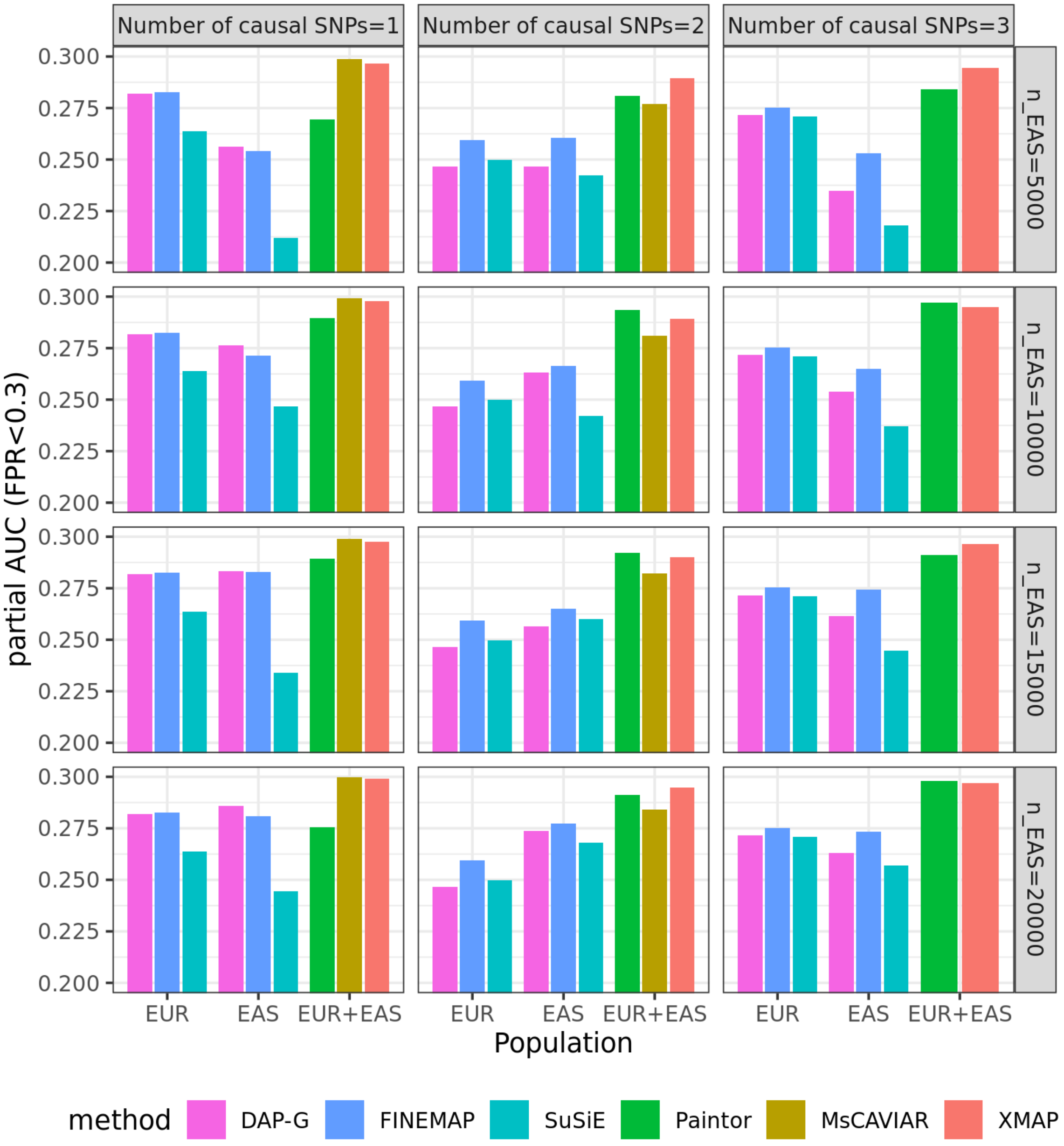

**Supplementary Figure 10:** Comparison of pAUC (FPR$<0.3$) among DAP-G, FINEMAP, SuSiE, Paintor, MsCAIVAR, and XMAP across 50 simulations in the setting without polygenic effects. We varied $K_{true} \in \{1, 2, 3\}$ and EAS sample size $n_1 \in \{5,000,\ 10,000.\ 15,000,\ 20,000\}$, and set EUR sample size $n_2 = 20,000$. Each of the $K_{true}$ causal SNPs explains 1% phenotypic variance. Because MsCAVIAR was intractable when including more then three causal signals, it was excluded from the comparison in the setting of $K_{true} = 3$.

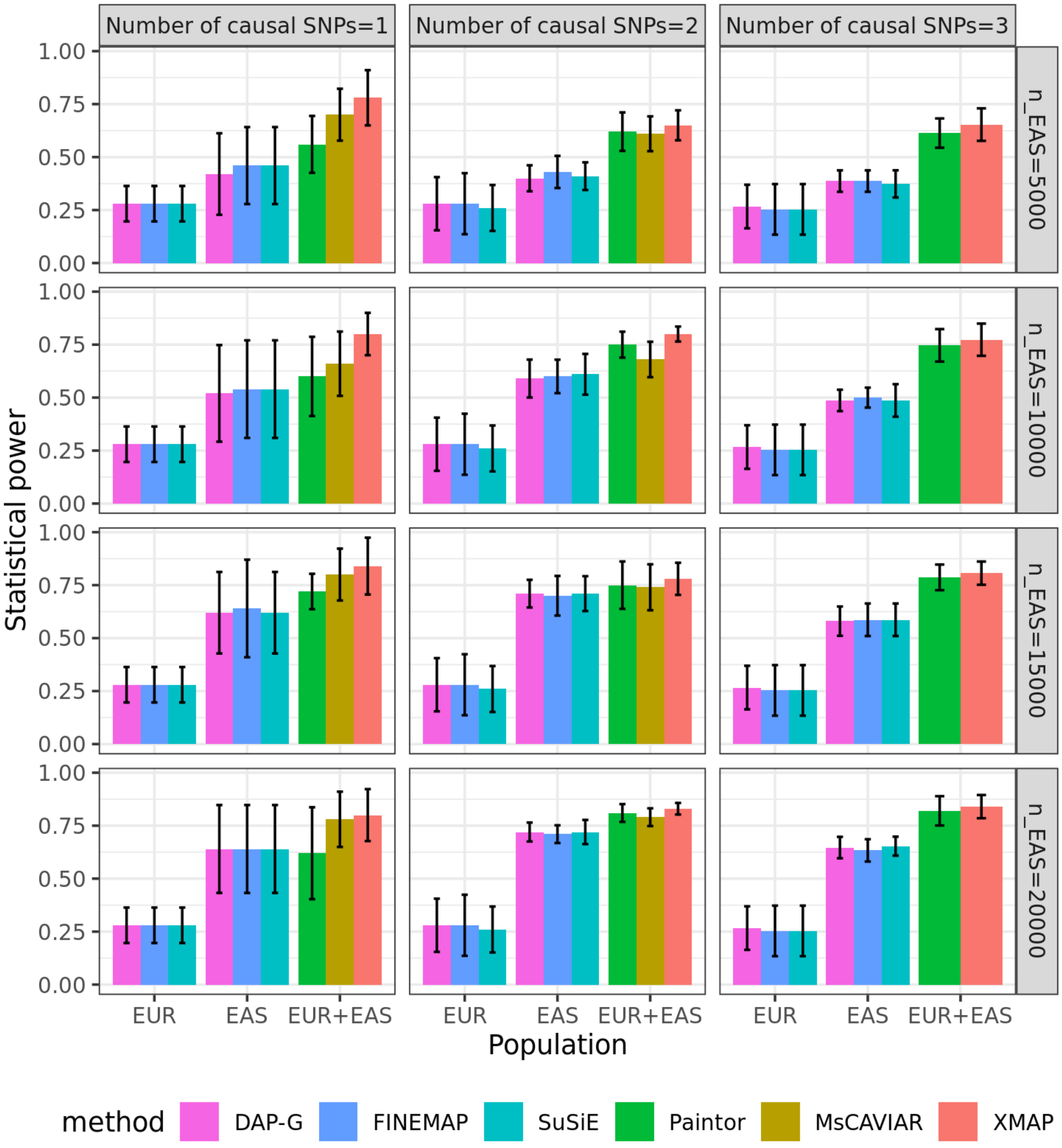

**Supplementary Figure 11:** Comparison of statistical power among DAP-G, FINEMAP, SuSiE, Paintor, MsCAIVAR, and XMAP in the setting without polygenic effects. We varied $K_{true} \in \{1, 2, 3\}$ and EAS sample size $n_1 \in \{5,000,\ 10,000.\ 15,000,\ 20,000\}$, and set EUR sample size $n_2 = 20,000$. Each of the $K_{true}$ causal SNPs explains 0.5% phenotypic variance. Because MsCAVIAR was intractable when including more then three causal signals, it was excluded from the comparison in the setting of $K_{true} = 3$. Error bars represent the standard errors of statistical powers evaluated on 50 replications.

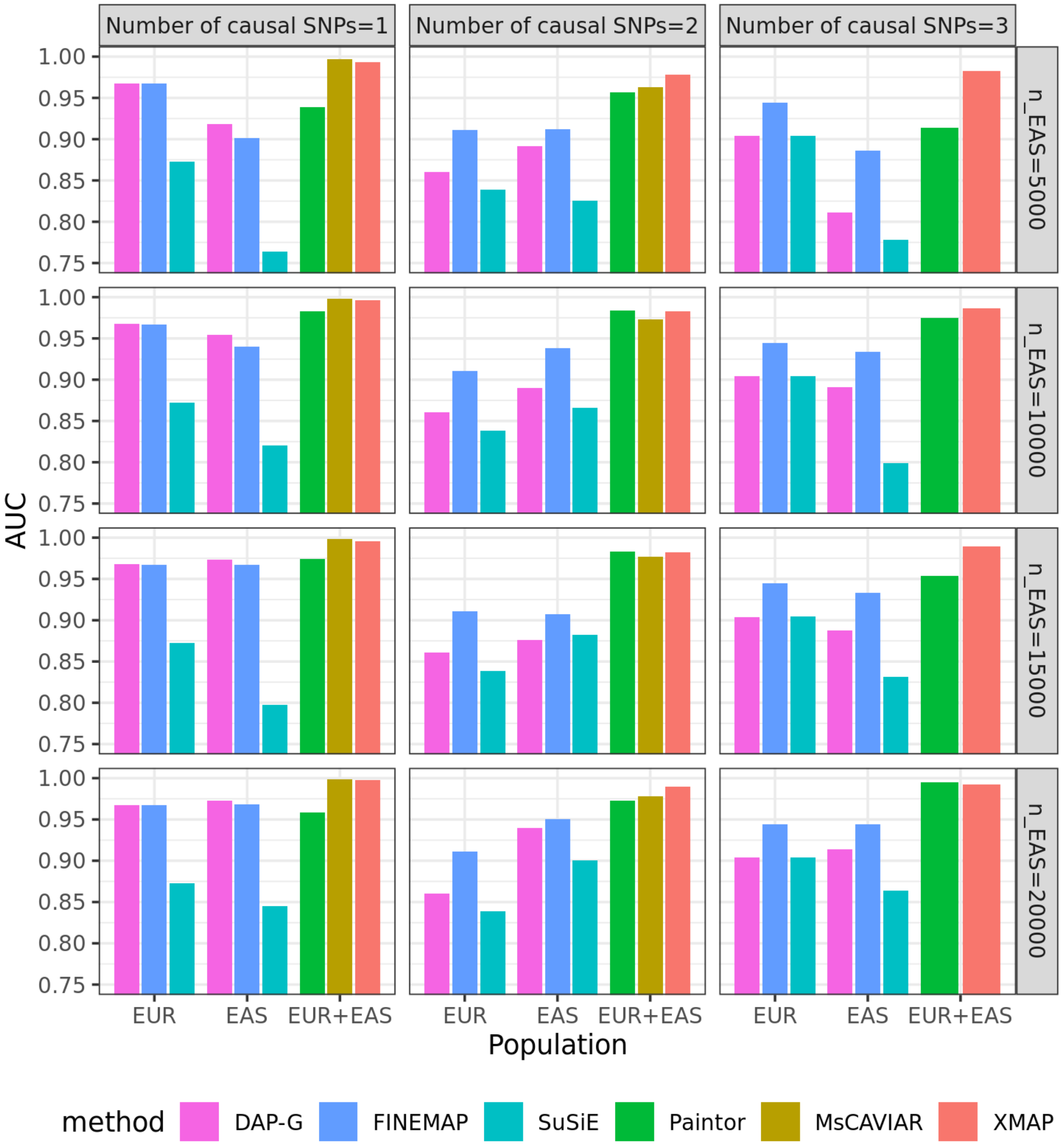

**Supplementary Figure 12:** Comparison of AUC among DAP-G, FINEMAP, SuSiE, Paintor, MsCAIVAR, and XMAP across 50 simulations in the setting without polygenic effects. We varied $K_{true} \in \{1, 2, 3\}$ and EAS sample size $n_1 \in \{5,000,\ 10,000.\ 15,000,\ 20,000\}$, and set EUR sample size $n_2 = 20,000$. Each of the $K_{true}$ causal SNPs explains 0.5% phenotypic variance. Because MsCAVIAR was intractable when including more then three causal signals, it was excluded from the comparison in the setting of $K_{true} = 3$.

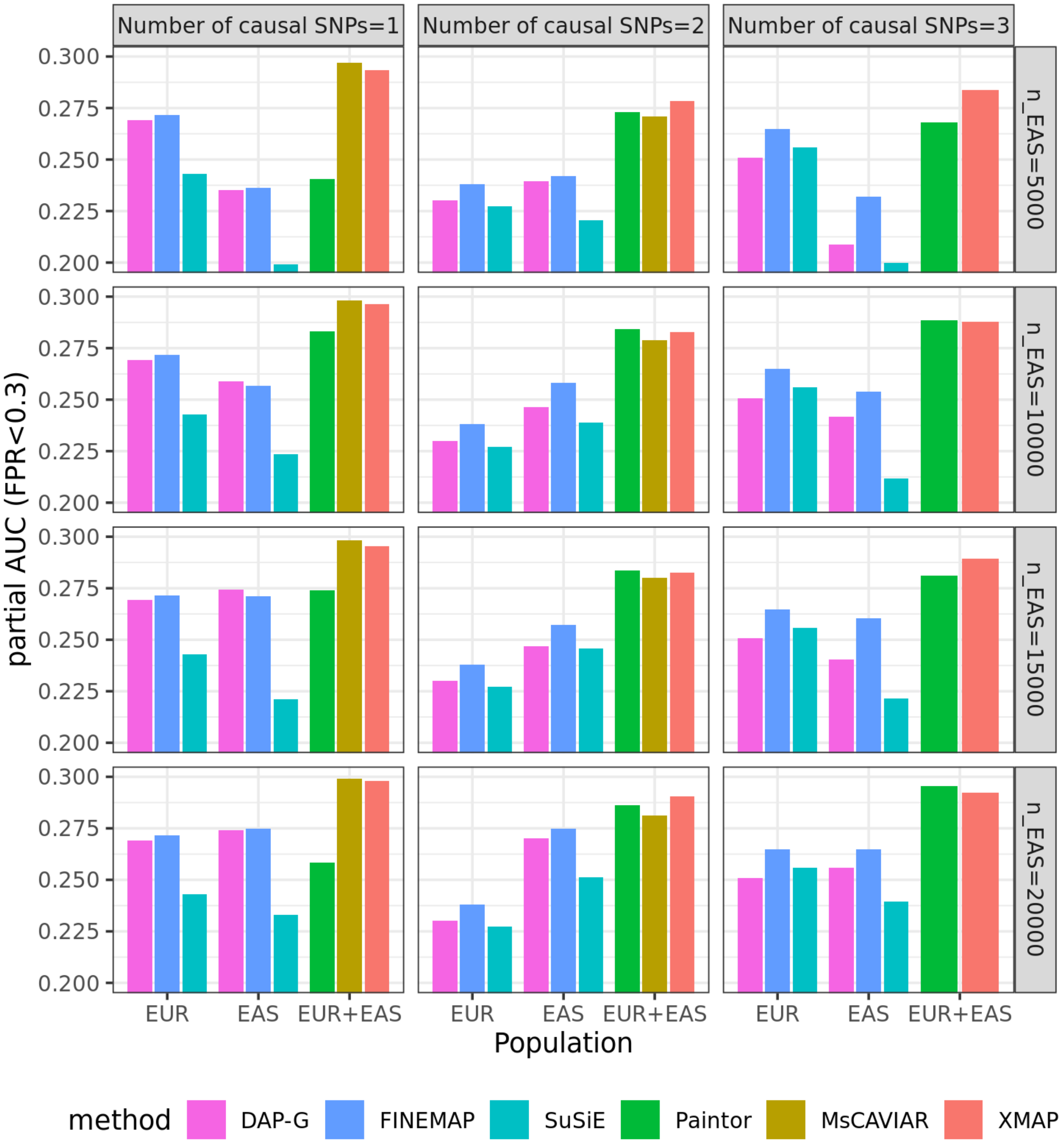

**Supplementary Figure 13:** Comparison of pAUC (FPR$<0.3$) among DAP-G, FINEMAP, SuSiE, Paintor, MsCAIVAR, and XMAP across 50 simulations in the setting without polygenic effects. We varied $K_{true} \in \{1, 2, 3\}$ and EAS sample size $n_1 \in \{5,000,\ 10,000.\ 15,000,\ 20,000\}$, and set EUR sample size $n_2 = 20,000$. Each of the $K_{true}$ causal SNPs explains 0.5% phenotypic variance. Because MsCAVIAR was intractable when including more then three causal signals, it was excluded from the comparison in the setting of $K_{true} = 3$.

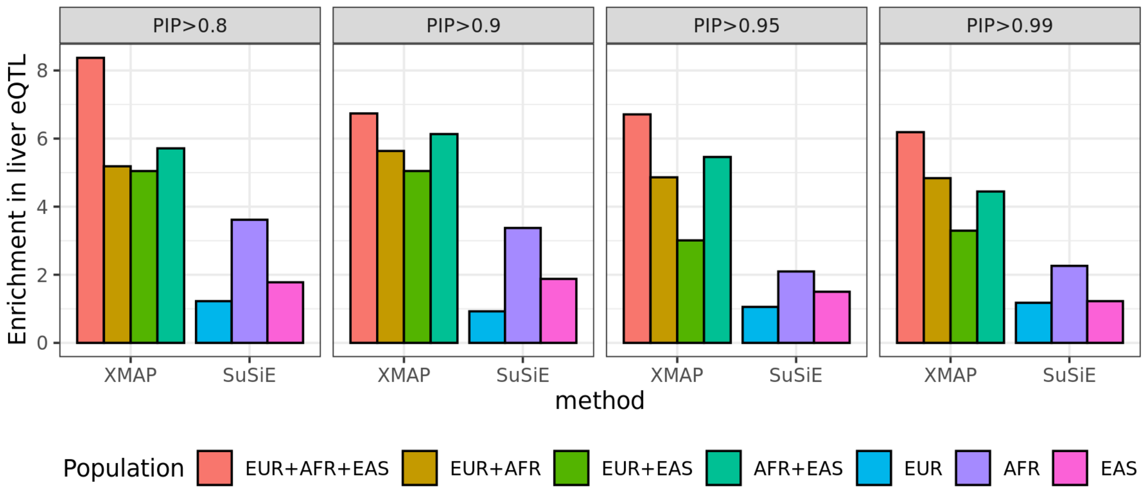

**Supplementary Figure 14:** Enrichment of fine-mapped SNPs in the eQTLs of liver obtained from the Genotype-Tissue Expression (GTEx) project. The enrichment is defined as the proportion of fine-mapped SNPs belonging to liver-eQTLs divided by the proportion of all analyzed SNPs belonging to liver-eQTLs. The SNPs identified by XMAP produced a substantially higher enrichment of LDL in the liver eQTLs, as compared to SNPs identified by SuSiE in a single population GWAS.

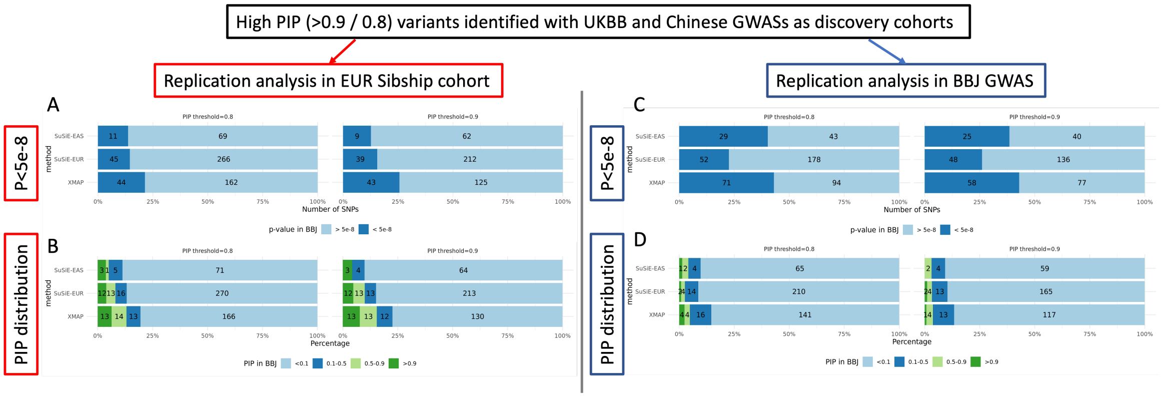

**Supplementary Figure 15:** Distributions of the number of putative causal SNPs identified by XMAP with $K = 15$ under different PIP thresholds.

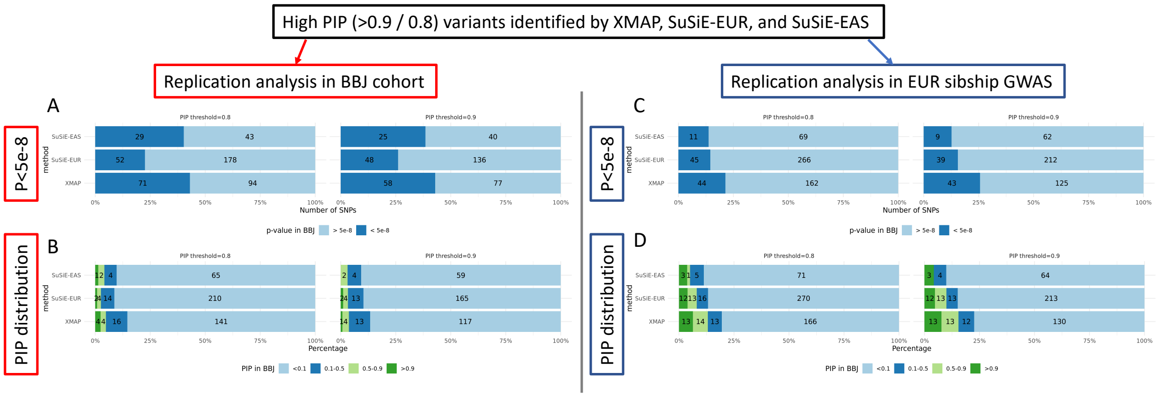

**Supplementary Figure 16:** Replication analysis of XMAP and SuSiE with $K = 15$ on height GWASs. Bar charts are shown for the fraction and number of fine-mapped SNPs with $p$-value$< 5 \times 10^{-8}$ in the replication cohorts of BBJ (top left) and EUR Sibship GWAS (top right), and the PIP distribution of fine-mapped SNPs in the replication cohorts of BBJ (bottom left) and EUR Sibship GWAS (bottom right).

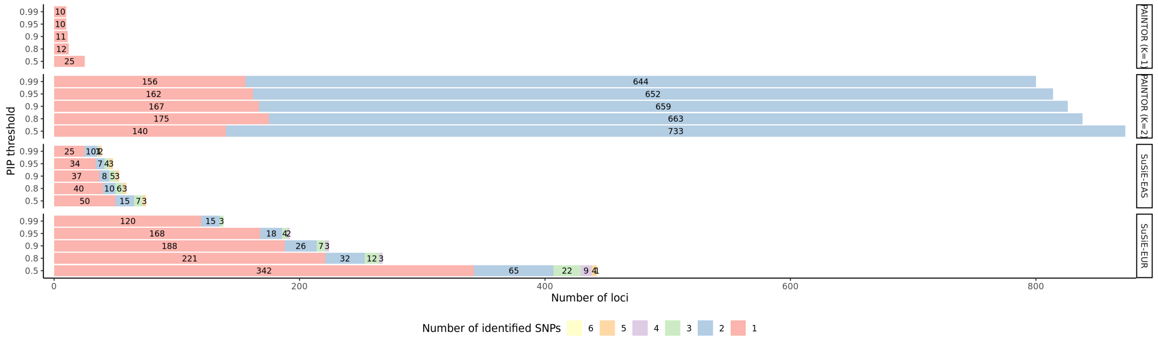

**Supplementary Figure 17:** Performance of PAINTOR (K=1), PAINTOR (K=2), SuSiE applied to EAS GWAS, and SuSiE applied to EUR GWAS in identifying multiple causal variants for height. Bar plots show the distributions of the number of putative causal SNPs under different PIP thresholds. PAINTOR had unstable performance when analyzing loci with thousands of SNPs. When K was set to 1 in PAINTOR (using the falg '-enumerate 1'), many loci had PIP=1 for all SNPs. We removed these loci when summarizing the results. When K was set to 2 in PAINTOR (using the falg '-enumerate 2'), many loci had exactly 2 SNPs with PIP=1, most of which could not be replicated in the Sibship GWAS (see Supplementary Figure18).

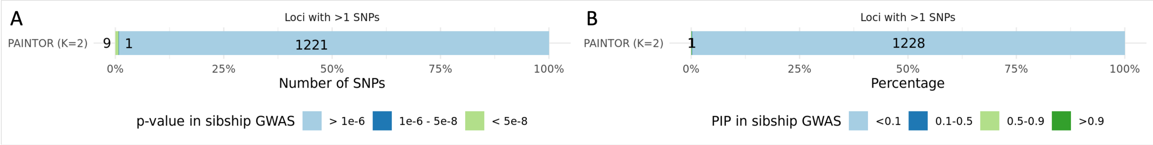

**Supplementary Figure 18:** Replication analysis of PAINTOR with K=2 on height GWASs. Bar charts are shown for the *p*-value (A) and PIP (B) distributions of putative causal SNPs in the replication cohort of EUR Sibship GWAS.

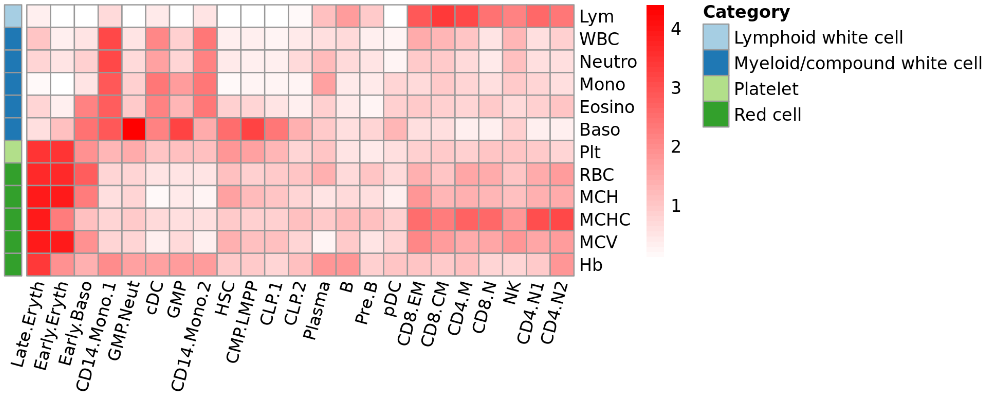

**Supplementary Figure 19:** The heat map showing the median TRS computed by SCAVENGE across 18 cell populations. The 12 blood traits are grouped into four clusters.

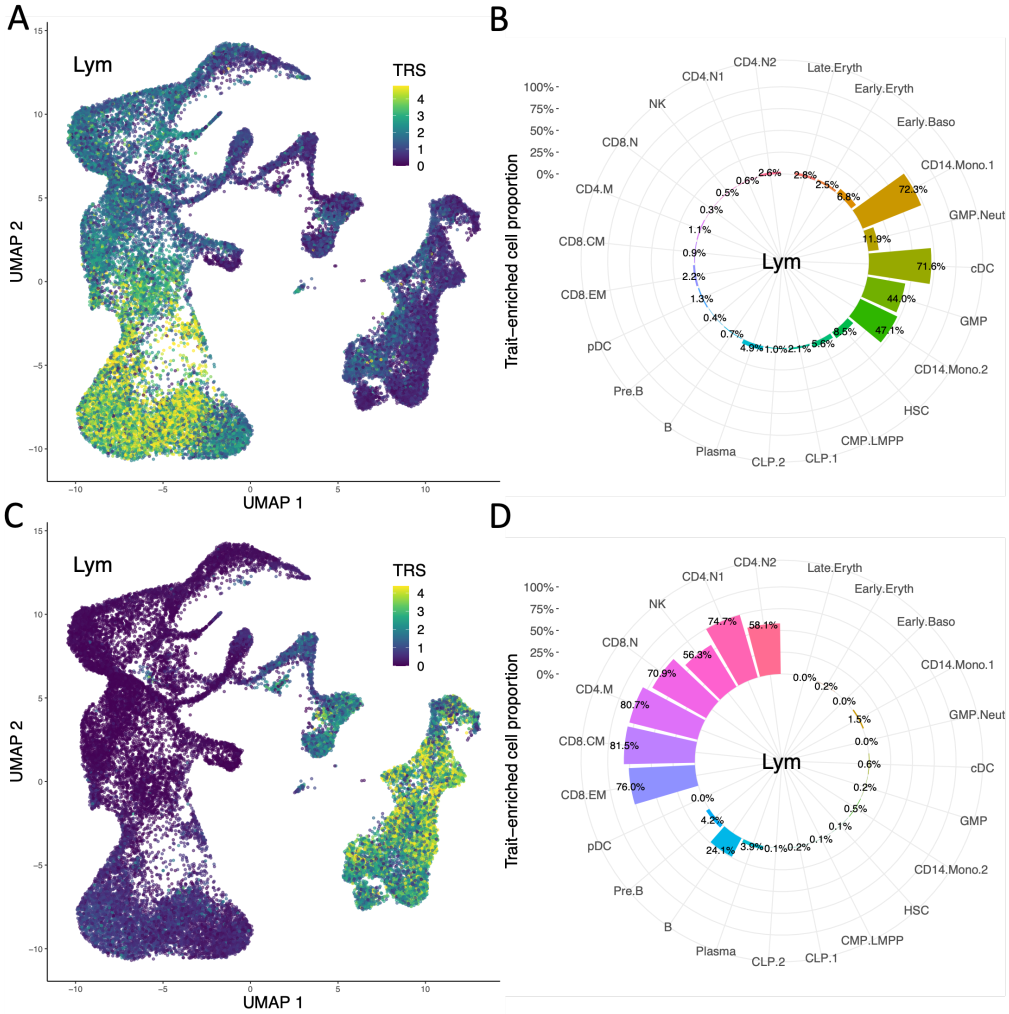

**Supplementary Figure 20:** Enrichment of the lymphocyte count in hematological populations using fine-mapped SNPs as input. The SCAVENGE TRS obtained by using the fine-mapping results of SuSiE in BBJ (A) and UKBB (C) are shown in the UMAP coordinates. The proportions of significantly enriched cells within each population obtained by using the fine-mapping results of SuSiE in BBJ (B) and UKBB (D).

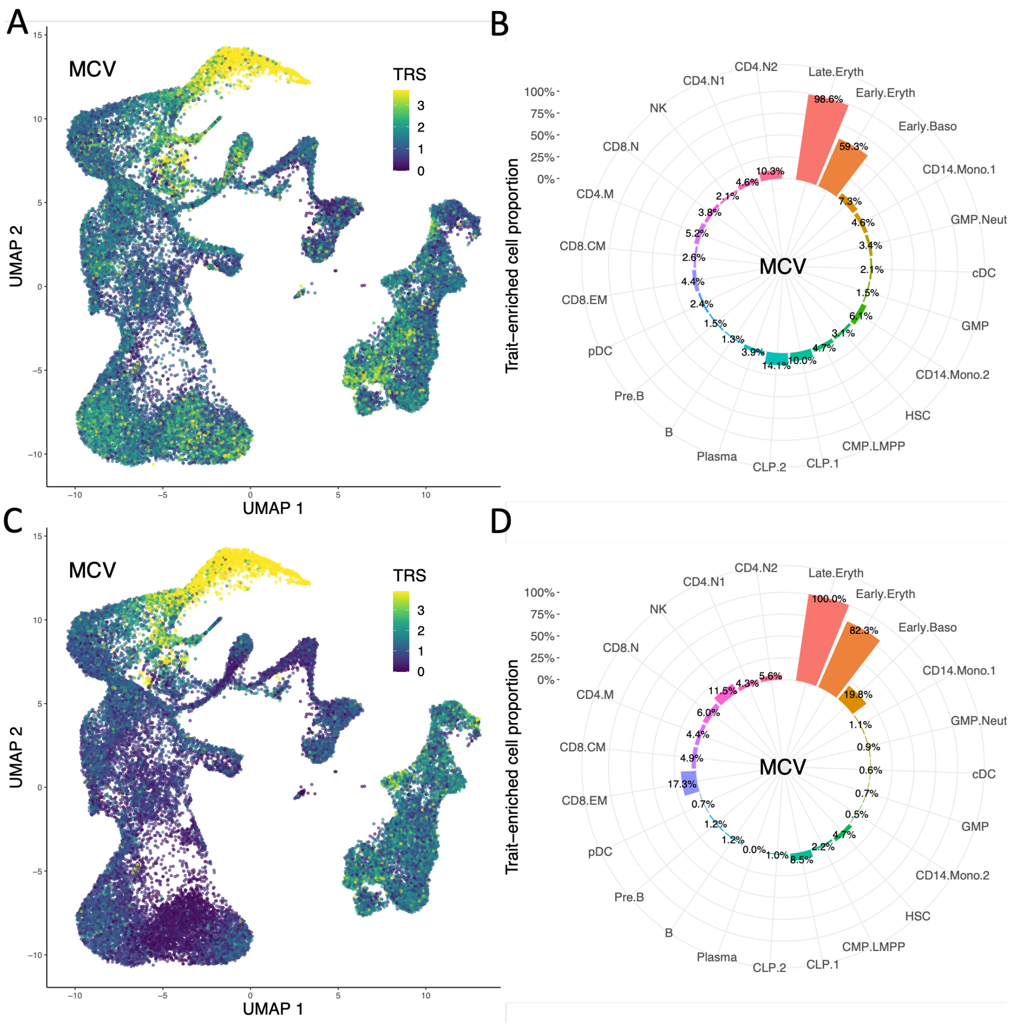

**Supplementary Figure 21:** Enrichment of the mean corpuscular volume in hematological populations using fine-mapped SNPs as input. The SCAVENGE TRS obtained by using the fine-mapping results of SuSiE in BBJ (A) and UKBB (C) are shown in the UMAP coordinates. The proportions of significantly enriched cells within each population obtained by using the fine-mapping results of SuSiE in BBJ (B) and UKBB (D).

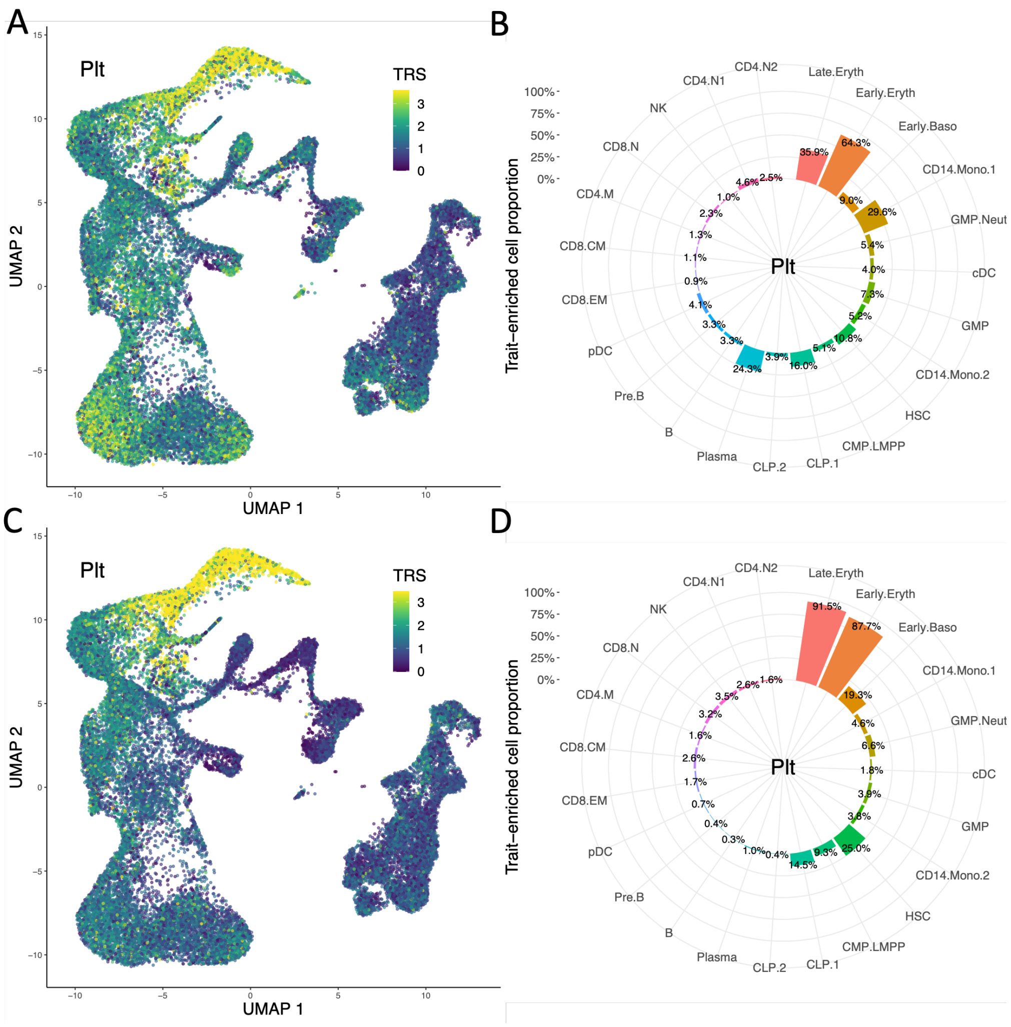

**Supplementary Figure 22:** Enrichment of the platelet count in hematological populations using fine-mapped SNPs as input. The SCAVENGE TRS obtained by using the fine-mapping results of SuSiE in BBJ (A) and UKBB (C) are shown in the UMAP coordinates. The proportions of significantly enriched cells within each population obtained by using the fine-mapping results of SuSiE in BBJ (B) and UKBB (D).

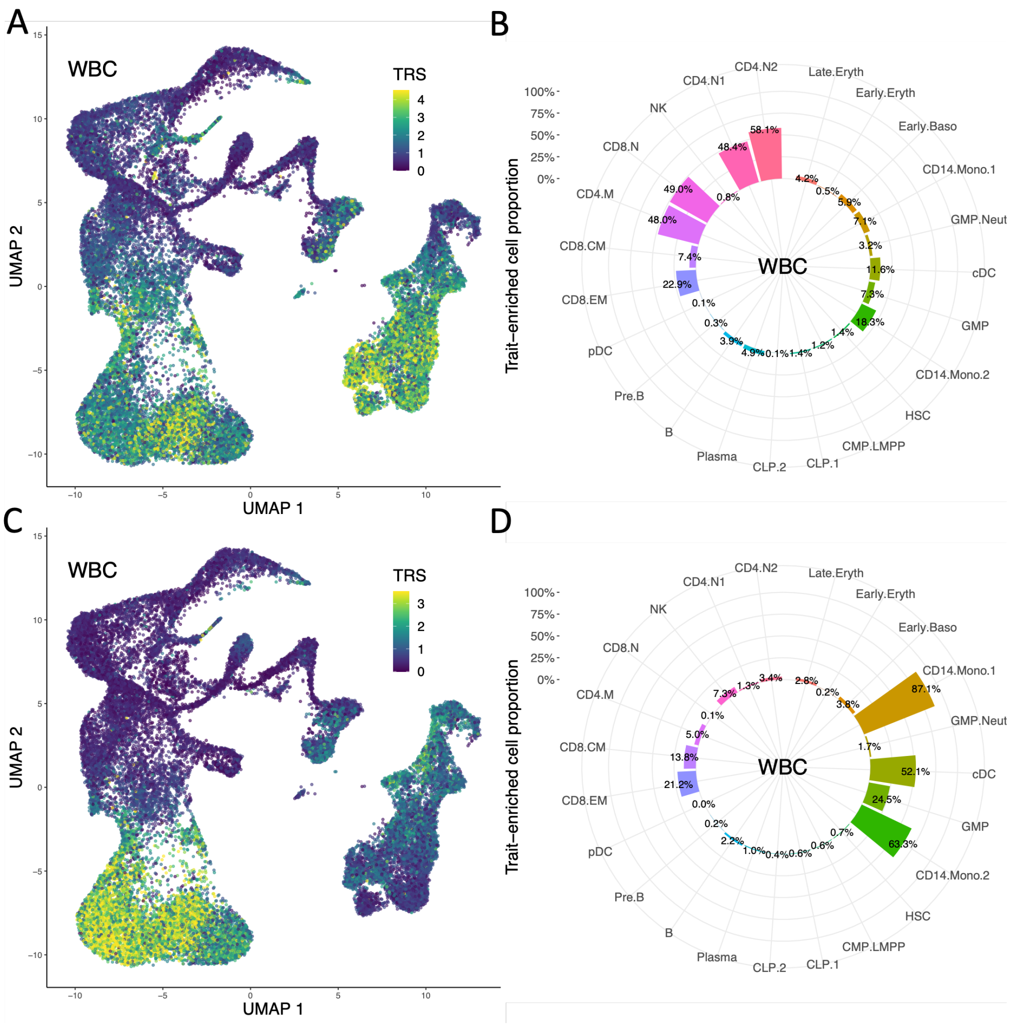

**Supplementary Figure 23:** Enrichment of the white blood cell count in hematological populations using fine-mapped SNPs as input. The SCAVENGE TRS obtained by using the fine-mapping results of SuSiE in BBJ (A) and UKBB (C) are shown in the UMAP coordinates. The proportions of significantly enriched cells within each population obtained by using the fine-mapping results of SuSiE in BBJ (B) and UKBB (D).

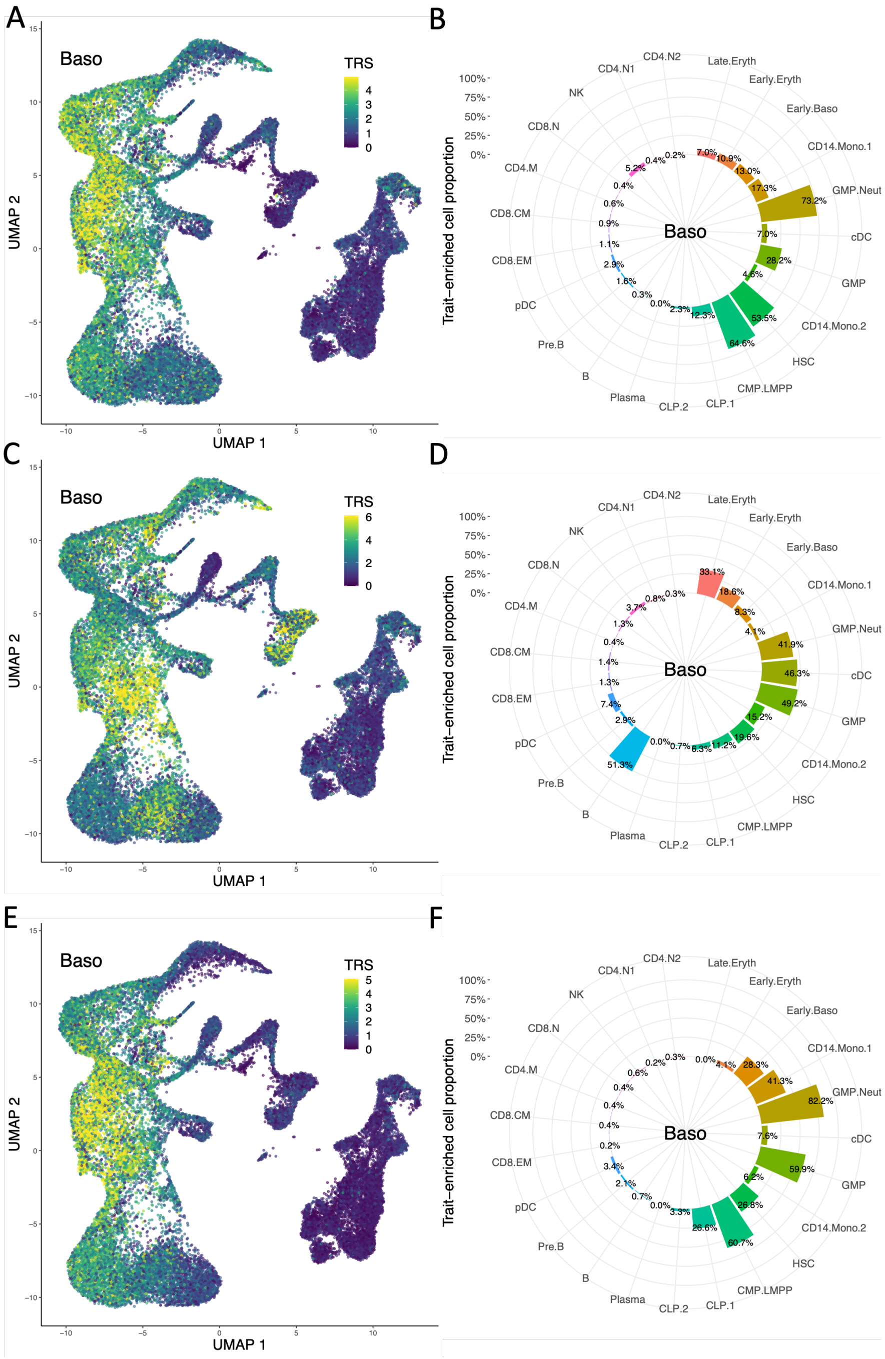

**Supplementary Figure 24:** Enrichment of the basophil count in hematological populations using fine-mapped SNPs as input. The SCAVENGE TRS obtained by using the fine-mapping results of XMAP (A) and SuSiE in BBJ (C) and UKBB (E) are shown in the UMAP coordinates. The proportions of significantly enriched cells within each population obtained by using the fine-mapping results of XMAP (B) and SuSiE in BBJ (D) and UKBB (F).

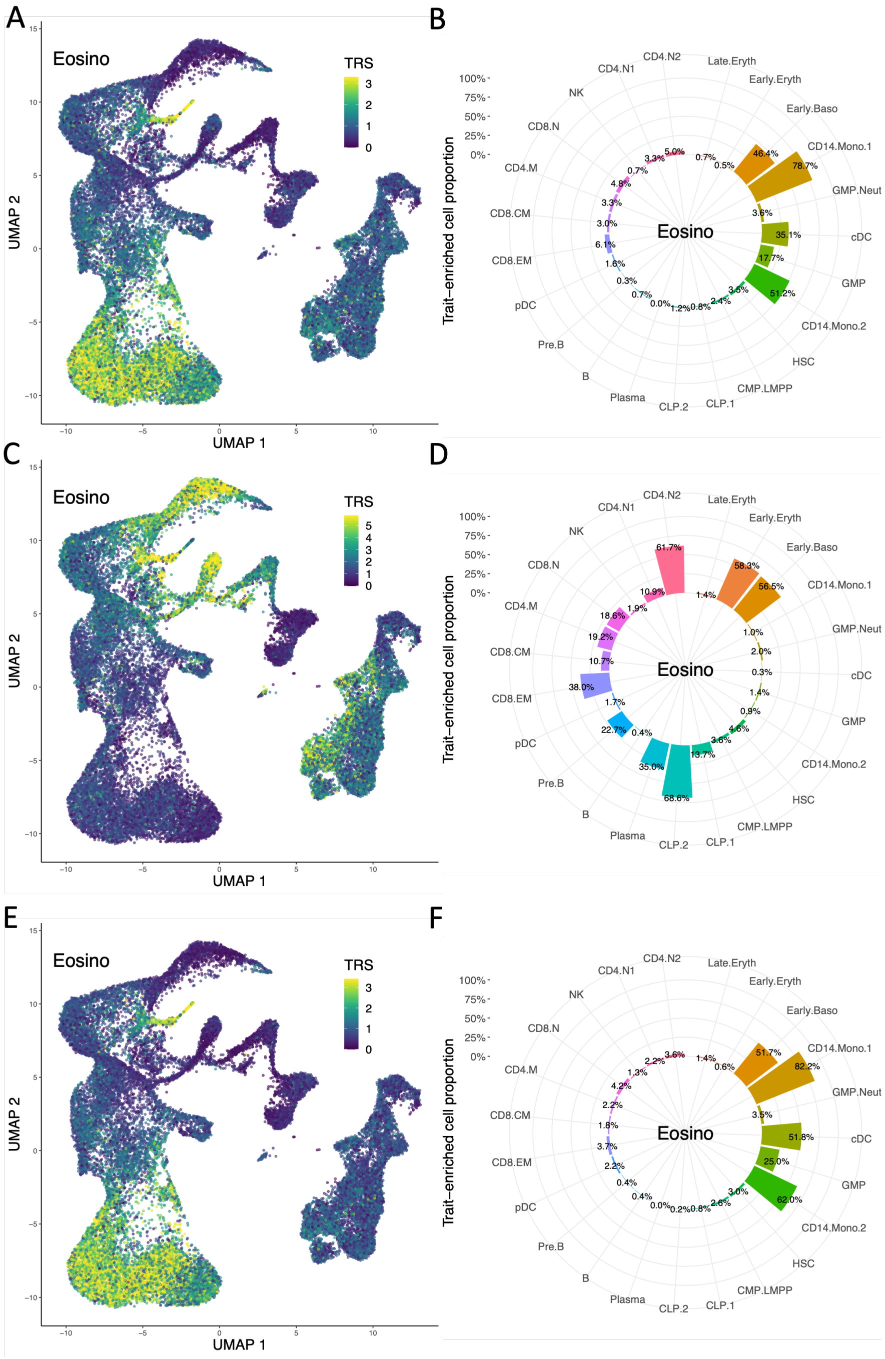

**Supplementary Figure 25:** Enrichment of the eosinophil count in hematological populations using fine-mapped SNPs as input. The SCAVENGE TRS obtained by using the fine-mapping results of XMAP (A) and SuSiE in BBJ (C) and UKBB (E) are shown in the UMAP coordinates. The proportions of significantly enriched cells within each population obtained by using the fine-mapping results of XMAP (B) and SuSiE in BBJ (D) and UKBB (F).

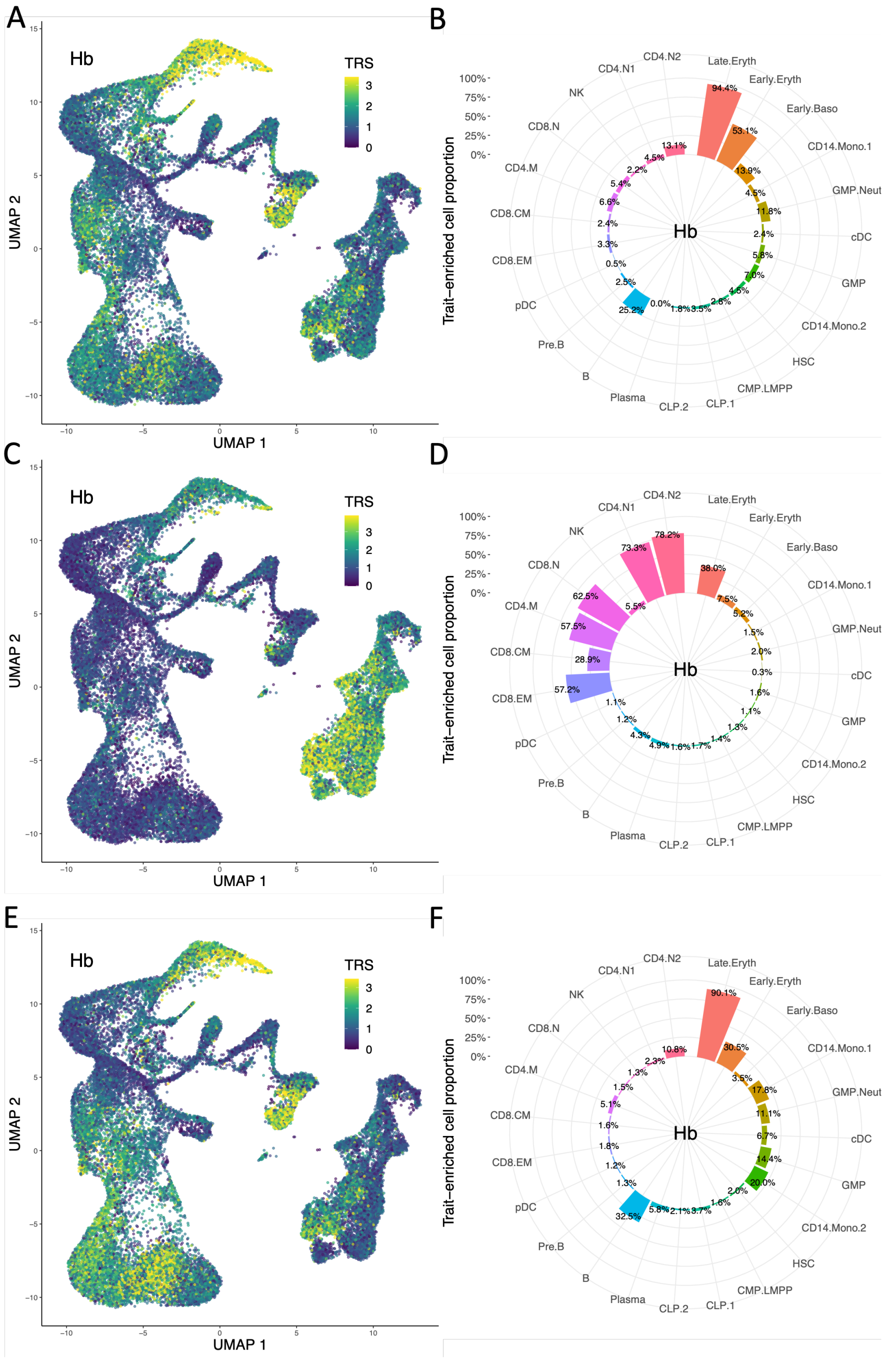

**Supplementary Figure 26:** Enrichment of the hemoglobin in hematological populations using fine-mapped SNPs as input. The SCAVENGE TRS obtained by using the fine-mapping results of XMAP (A) and SuSiE in BBJ (C) and UKBB (E) are shown in the UMAP coordinates. The proportions of significantly enriched cells within each population obtained by using the fine-mapping results of XMAP (B) and SuSiE in BBJ (D) and UKBB (F).

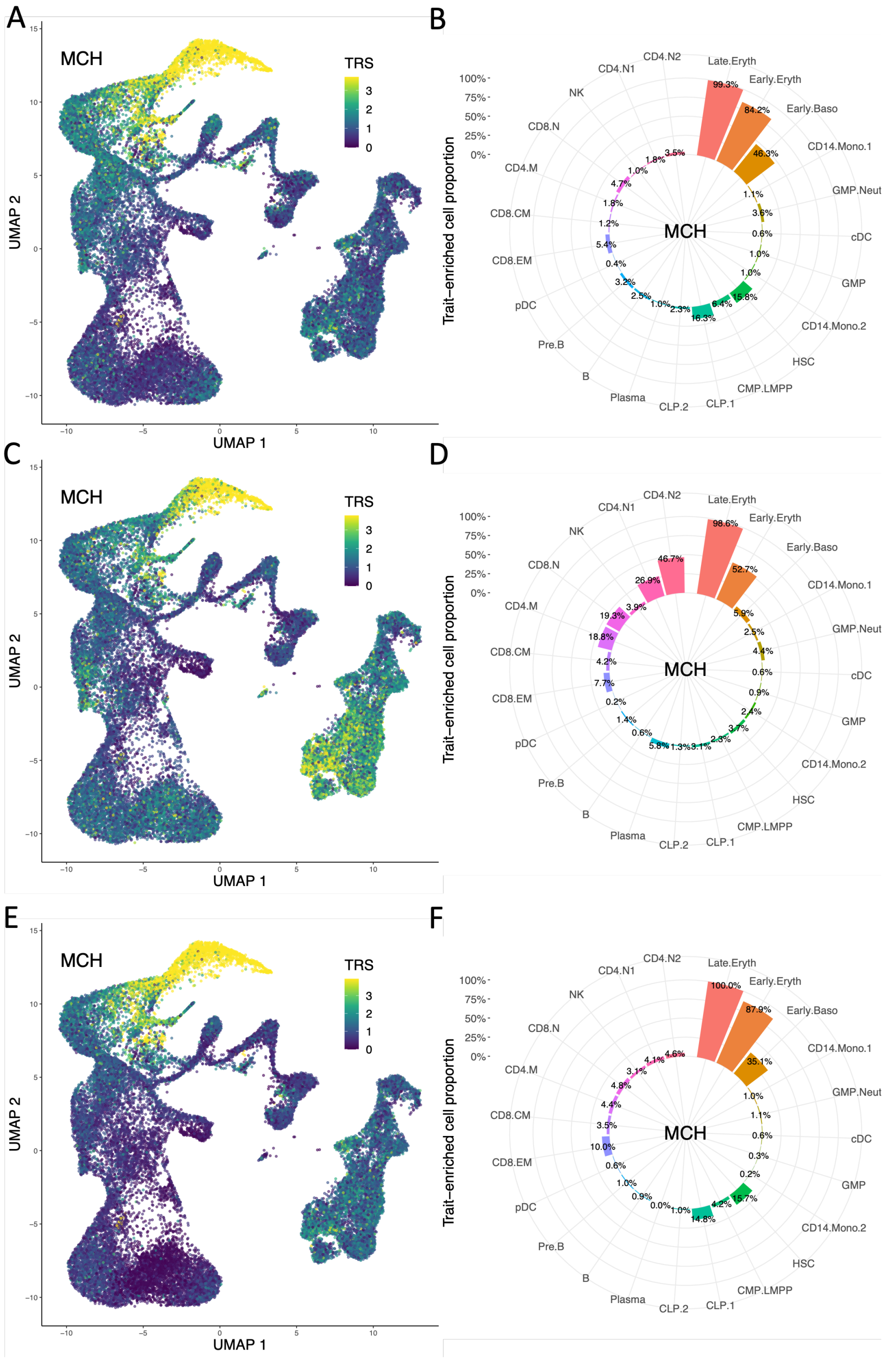

**Supplementary Figure 27:** Enrichment of the mean corpuscular hemoglobin in hematological populations using fine-mapped SNPs as input. The SCAVENGE TRS obtained by using the fine-mapping results of XMAP (A) and SuSiE in BBJ (C) and UKBB (E) are shown in the UMAP coordinates. The proportions of significantly enriched cells within each population obtained by using the fine-mapping results of XMAP (B) and SuSiE in BBJ (D) and UKBB (F).

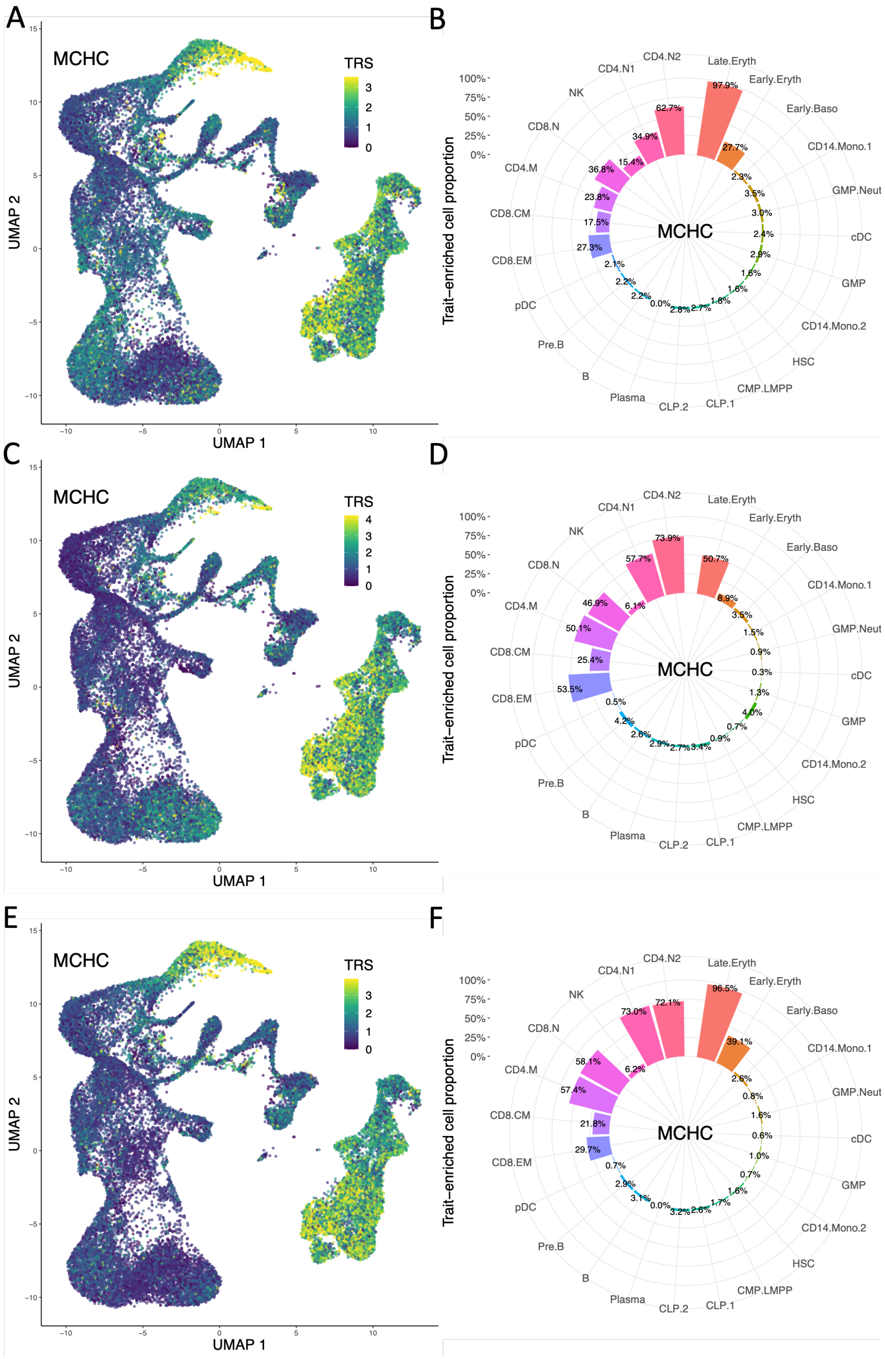

**Supplementary Figure 28:** Enrichment of the mean corpuscular hemoglobin concentration in hematological populations using fine-mapped SNPs as input. The SCAVENGE TRS obtained by using the fine-mapping results of XMAP (A) and SuSiE in BBJ (C) and UKBB (E) are shown in the UMAP coordinates. The proportions of significantly enriched cells within each population obtained by using the fine-mapping results of XMAP (B) and SuSiE in BBJ (D) and UKBB (F).

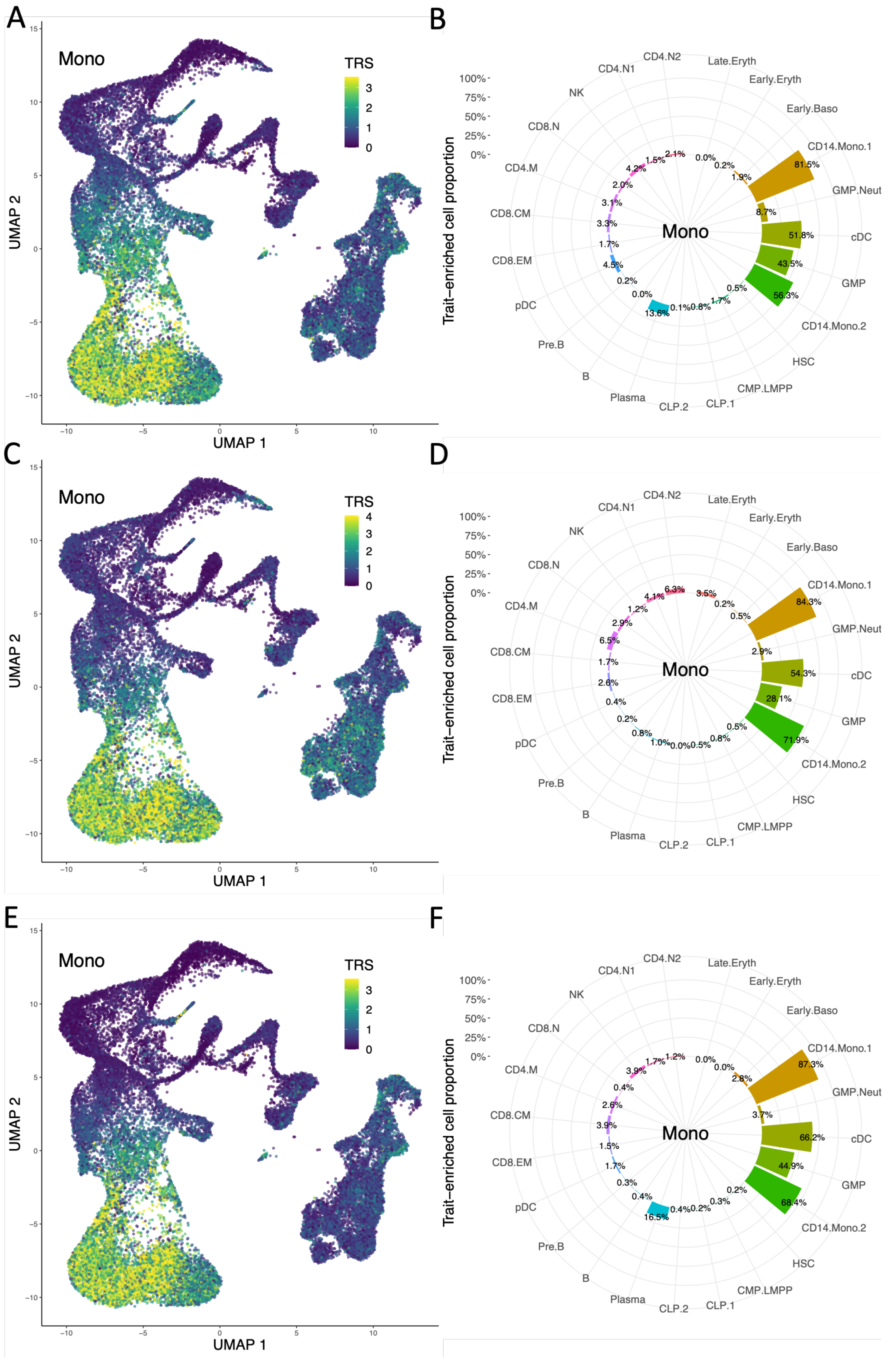

**Supplementary Figure 29:** Enrichment of the monocyte count in hematological populations using fine-mapped SNPs as input. The SCAVENGE TRS obtained by using the fine-mapping results of XMAP (A) and SuSiE in BBJ (C) and UKBB (E) are shown in the UMAP coordinates. The proportions of significantly enriched cells within each population obtained by using the fine-mapping results of XMAP (B) and SuSiE in BBJ (D) and UKBB (F).

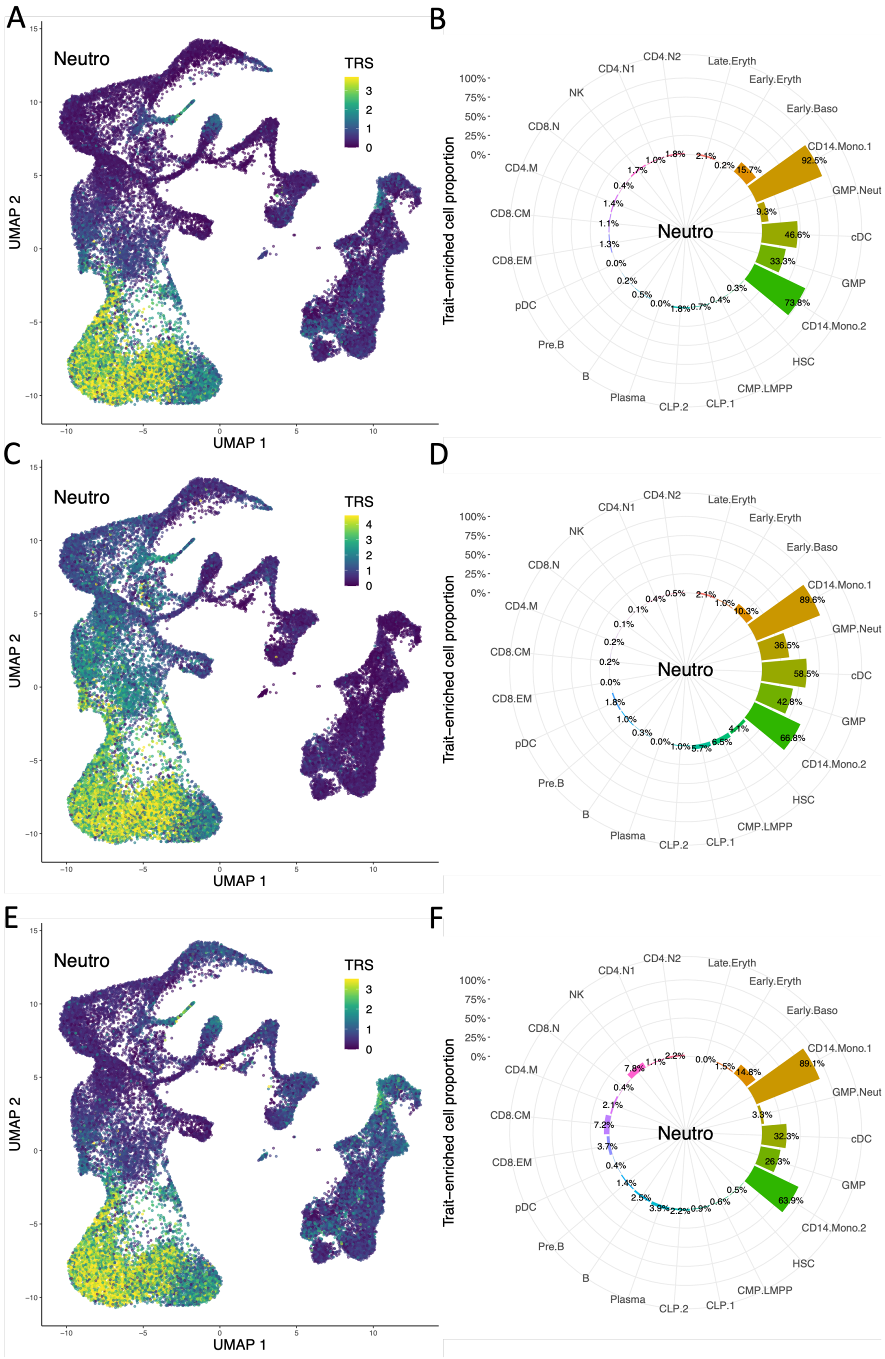

**Supplementary Figure 30:** Enrichment of the neutrophil count in hematological populations using fine-mapped SNPs as input. The SCAVENGE TRS obtained by using the fine-mapping results of XMAP (A) and SuSiE in BBJ (C) and UKBB (E) are shown in the UMAP coordinates. The proportions of significantly enriched cells within each population obtained by using the fine-mapping results of XMAP (B) and SuSiE in BBJ (D) and UKBB (F).

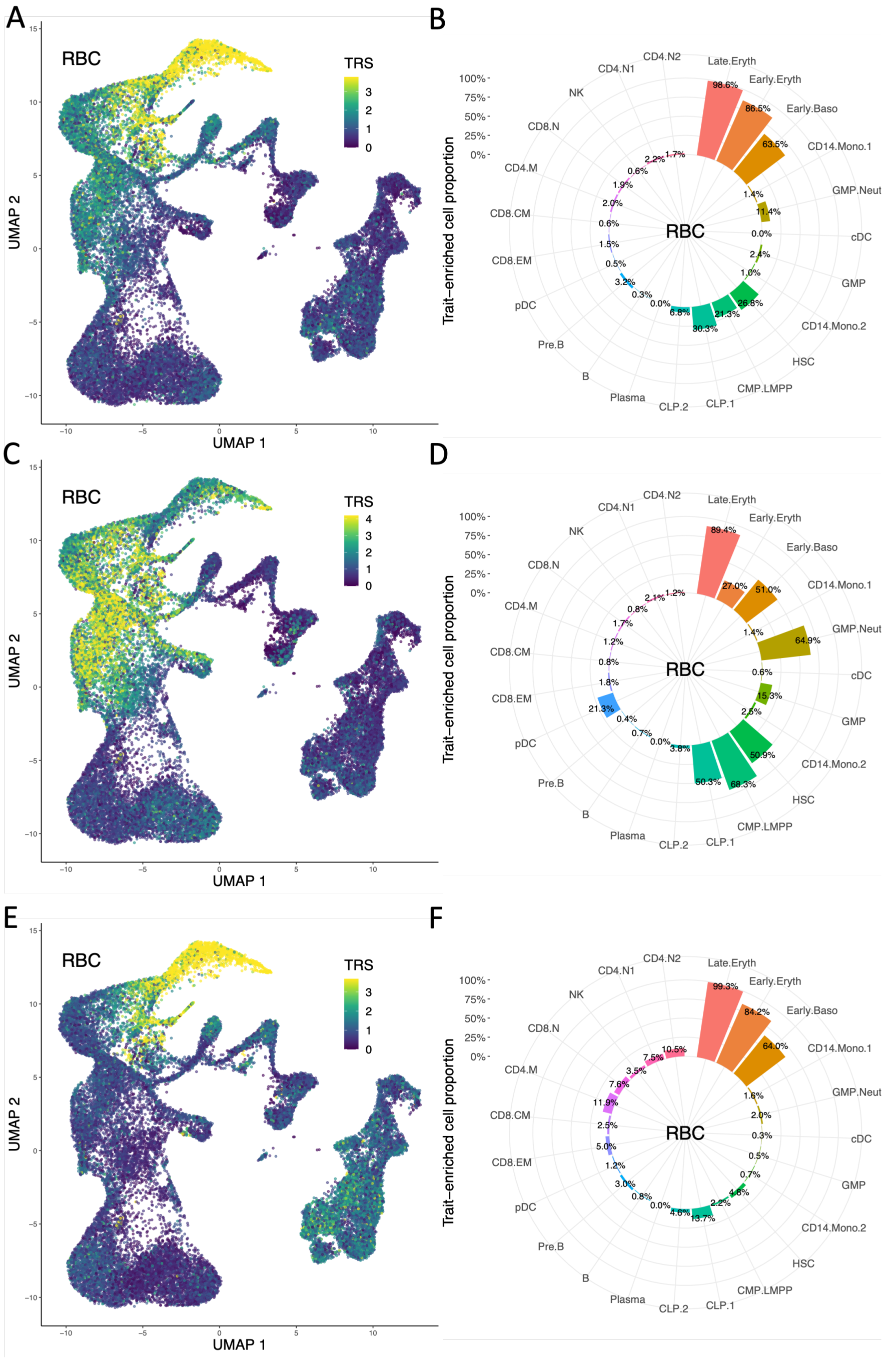

**Supplementary Figure 31:** Enrichment of the red blood cell count in hematological populations using fine-mapped SNPs as input. The SCAVENGE TRS obtained by using the fine-mapping results of XMAP (A) and SuSiE in BBJ (C) and UKBB (E) are shown in the UMAP coordinates. The proportions of significantly enriched cells within each population obtained by using the fine-mapping results of XMAP (B) and SuSiE in BBJ (D) and UKBB (F).

### 3 Supplementary Note

#### 3.1 Simulations without polygenic effects and confounding bias

Besides the settings considered in the main text, we conducted additional simulations to investigate the performance of XMAP in the absence of either polygenic effects and confounding bias. In this simulation, we generated the causal effects with $\beta_{1k} \sim \mathcal{N}(0, \omega_1)$ and $\beta_{2k} \sim \mathcal{N}(0, \omega_2)$ for $k = 1, ..., K_{true}$, where $\omega_1$ and $\omega_2$ are the per-SNP heritabilities explained by the causal SNPs in populations 1 and 2, respectively, and we varied $\omega_1 = \omega_2 \in \{0.01, 0.005\}$. We generated quantitative phenotypes in the two populations with $\mathbf{y}_1 = \sum_{k=1}^{K_{true}} \mathbf{x}_{1[k]}\beta_{1k} + \mathbf{e}_2$ and $\mathbf{y}_2 = \sum_{k=1}^{K_{true}} \mathbf{x}_{2[k]}\beta_{2k} + \mathbf{e}_2$, where $\mathbf{x}_{1[k]}$ and $\mathbf{x}_{2[k]}$ are the columns of $\mathbf{X}_1$ and $\mathbf{X}_2$ corresponding to the $k$-th causal SNP, and $\mathbf{e}_1 \sim \mathcal{N}(\mathbf{0}, (1 - \omega_1 \times K_{true})\mathbf{I}_{n_1})$ and $\mathbf{e}_2 \sim \mathcal{N}(\mathbf{0}, (1 - \omega_2 \times K_{true})\mathbf{I}_{n_2})$ are independent noise in the two populations, respectively. The results are summarized in Supplementary Figures 8-13. Under this setting, the performance of XMAP was comparable to with PAINTOR and MsCAVIAR.

#### 3.2 Compared methods

We compared XMAP with existing fine-mapping approaches in the simulation analysis. In the main analysis of XMAP, we estimated the polygenic parameters $\mathbf{\Omega}$ and inflation constants $c_1$ and $c_2$ with bivariate LDSC, and considered two settings of $K$: $K = 5$ and $K = 10$. For single-population approaches, we considered FINEMAP, Dap-G, and SuSiE. We ran FINEMAP with the shot gun stochastic search algorithm by using the flag '–sss' and set '–n-causal-snps' as the default value 5. For SuSiE, we set the maximum number of causal signals $L = 5$. We also included two cross-population approaches, MsCAVIAR and PAINTOR, in our analysis. We ran MsCAVIAR with the flag '-c $K_{true}$', and ran PAINTOR with the flag '-enumerate $K_{true}$'.

In real data analysis, we used $K = 10$ in XMAP for the main analysis and conducted sensitivity analysis with $K = 15$. For SuSiE, we set $L = 10$ in all populations. When applying PAINTOR to screen all loci on the genome, we considered two settings: '-enumerate 1' and '-enumerate 2'. In the example presented in Figure 6, we used the flags '-enumerate 2' in PAINTOR and '-c 2' in MsCAVIAR.

#### 3.3 Derivation of the covariance of $\boldsymbol{\epsilon}_1$ and $\boldsymbol{\epsilon}_2$

We first derive the first moment of $\mathbf{b}_1$ and $\mathbf{b}_2$:

$$\begin{aligned}
\mathbb{E}[\mathbf{b}_1] &= \mathbb{E}\left[\sum_{k=1}^{K} \boldsymbol{\gamma}_k \beta_{1k}\right] = \sum_{k=1}^{K} \mathbb{E}[\mathbb{E}[\boldsymbol{\gamma}_k \beta_{1k} | \boldsymbol{\gamma}_k]] = \mathbf{0} \\
\mathbb{E}[\mathbf{b}_2] &= \mathbb{E}\left[\sum_{k=1}^{K} \boldsymbol{\gamma}_k \beta_{2k}\right] = \sum_{k=1}^{K} \mathbb{E}[\mathbb{E}[\boldsymbol{\gamma}_k \beta_{2k} | \boldsymbol{\gamma}_k]] = \mathbf{0}.
\end{aligned} \tag{S1}$$

By applying the law of total variance, the second moment of $\mathbf{b}_1$ can be obtained as:

$$\begin{aligned}
&\mathbb{E}\left[\mathbf{b}_1\mathbf{b}_1^T\right] = \mathrm{Var}\left[\mathbf{b}_1\right]\\
=&\mathrm{Var}\left[\sum_{k=1}^{K}\mathbb{E}\left[\boldsymbol{\gamma}_k\beta_{1k}|\boldsymbol{\gamma}_k\right]\right] + \mathbb{E}\left[\sum_{k=1}^{K}\mathrm{Var}\left[\boldsymbol{\gamma}_k\beta_{1k}|\boldsymbol{\gamma}_k\right]\right]\\
=&\mathbf{0} + \frac{1}{p}\sum_{k=1}^{K}\sum_{j=1}^{p}\sigma_{1k}^2\mathbf{I}_p\\
=&\sum_{k=1}^{K}\sigma_{1k}^2\mathbf{I}_p.
\end{aligned} \tag{S2}$$

Similarly, we have $\mathbb{E}\left[\mathbf{b}_2\mathbf{b}_2^T\right] = \mathrm{Var}\left[\mathbf{b}_2\right] = \sum_{k=1}^{K}\sigma_{2k}^2\mathbf{I}_p$. With the above equations, we can obtain the first moments of $\hat{\mathbf{b}}_1$ and $\hat{\mathbf{b}}_2$:

$$\begin{aligned}
\mathbb{E}\left[\hat{\mathbf{b}}_1\right] &= \mathbb{E}\left[\mathbb{E}\left[\hat{\mathbf{b}}_1|\mathbf{b}_1,\boldsymbol{\phi}_1\right]\right] = \mathbb{E}\left[\mathbb{E}\left[\mathbf{R}_1\mathbf{b}_1 + \mathbf{R}_1\boldsymbol{\phi}_1|\mathbf{b}_1,\boldsymbol{\phi}_1\right]\right] = \mathbf{0}\\
\mathbb{E}\left[\hat{\mathbf{b}}_2\right] &= \mathbb{E}\left[\mathbb{E}\left[\hat{\mathbf{b}}_2|\mathbf{b}_2,\boldsymbol{\phi}_2\right]\right] = \mathbb{E}\left[\mathbb{E}\left[\mathbf{R}_2\mathbf{b}_2 + \mathbf{R}_2\boldsymbol{\phi}_2|\mathbf{b}_2,\boldsymbol{\phi}_2\right]\right] = \mathbf{0}.
\end{aligned} \tag{S3}$$

Next, we derive the variance-covariance matrix of of $\hat{\mathbf{b}}_1$. The covariance between the $j$-th and the $j'$-th elements of $\hat{\mathbf{b}}_1$ is given as

$$\begin{aligned}
&\mathrm{Cov}[\hat{\mathbf{b}}_{1j}, \hat{\mathbf{b}}_{1j'}] = \mathbb{E}\left[\hat{\mathbf{b}}_{1j}\hat{\mathbf{b}}_{1j'}\right] \\
=&\mathbb{E}\left[\mathbb{E}\left[\hat{\mathbf{b}}_{1j}\hat{\mathbf{b}}_{1j'}|\mathbf{X}_1\right]\right] \\
=&\mathbb{E}\left[\mathbb{E}\left[\frac{1}{n_1^2}\mathbf{x}_{1j}^T\mathbf{y}_1\mathbf{y}_1^T\mathbf{x}_{1j'}|\mathbf{X}_1\right]\right] \\
=&\mathbb{E}\left[\mathbb{E}\left[\frac{1}{n_1^2}\mathbf{x}_{1j}^T(\mathbf{X}_1\mathbf{b}_1 + \mathbf{X}_1\boldsymbol{\phi}_1 + \mathbf{e}_1)(\mathbf{X}_1\mathbf{b}_1 + \mathbf{X}_1\boldsymbol{\phi}_1 + \mathbf{e}_1)^T\mathbf{x}_{1j'}|\mathbf{X}_1\right]\right] \\
=&\mathbb{E}\left[\mathbb{E}\left[\frac{1}{n_1^2}\mathbf{x}_{1j}^T(\mathbf{X}_1\mathbf{b}_1\mathbf{b}_1^T\mathbf{X}_1^T + \mathbf{X}_1\boldsymbol{\phi}_1\boldsymbol{\phi}_1^T\mathbf{X}_1^T + \mathbf{e}_1\mathbf{e}_1^T + 2\mathbf{X}_1\mathbf{b}_1\boldsymbol{\phi}_1^T\mathbf{X}_1^T + 2\mathbf{X}_1\mathbf{b}_1\mathbf{e}_1^T + 2\mathbf{X}_1\boldsymbol{\phi}_1\mathbf{e}_1^T)\mathbf{x}_{1j'}|\mathbf{X}_1\right]\right] \\
=&\mathbb{E}\left[\frac{1}{n_1^2}\mathbf{x}_{1j}^T\mathbf{X}_1\mathbb{E}\left[\mathbf{b}_1\mathbf{b}_1^T\right]\mathbf{X}_1^T\mathbf{x}_{1j'} + \frac{1}{n_1^2}\mathbf{x}_{1j}^T\mathbf{X}_1\mathbb{E}\left[\boldsymbol{\phi}_1\boldsymbol{\phi}_1^T\right]\mathbf{X}_1^T\mathbf{x}_{1j'} + \frac{1}{n_1^2}\mathbf{x}_{1j}^T\mathbb{E}\left[\mathbf{e}_1\mathbf{e}_1^T\right]\mathbf{x}_{1j'}\right] \\
\overset{①}{=}&\frac{1}{n_1^2}(\sum_{k=1}^K \sigma_{k1}^2)\mathbb{E}\left[\mathbf{x}_{1j}^T\mathbf{X}_1\mathbf{X}_1^T\mathbf{x}_{1j'}\right] + \frac{1}{n_1^2}\omega_1\mathbb{E}\left[\mathbf{x}_{1j}^T\mathbf{X}_1\mathbf{X}_1^T\mathbf{x}_{1j'}\right] + \frac{1}{n_1^2}\sigma_{\mathbf{e}_1}^2\mathbb{E}\left[\mathbf{x}_{1j}^T\mathbf{x}_{1j'}\right] \\
=&(\sum_{k=1}^K \sigma_{k1}^2 + \omega_1)\mathbb{E}\left[\mathbf{x}_{1j}^T\mathbf{X}_1\mathbf{X}_1^T\mathbf{x}_{1j'}/n_1^2\right] + \frac{1}{n_1}\sigma_{\mathbf{e}_1}^2 r_{1jj'} \\
\overset{②}{\approx}&(\sum_{k=1}^K \sigma_{k1}^2 + \omega_1)\sum_{l=1}^p r_{1jl}r_{1j'l} + \frac{p}{n_1}(\sum_{k=1}^K \sigma_{k1}^2 + \omega_1)r_{1jj'} + \frac{1}{n_1}\sigma_{\mathbf{e}_1}^2 r_{1jj'} \\
\overset{③}{=}&(\sum_{k=1}^K \sigma_{k1}^2 + \omega_1)\sum_{l=1}^p r_{1jl}r_{1j'l} + \frac{1}{n_1}r_{1jj'} \\
\overset{④}{\approx}&(\sum_{k=1}^K \sigma_{k1}^2 + \omega_1)\sum_{l=1}^p r_{1jl}r_{1j'l} + \hat{s}_{1j}\hat{s}_{1j'}r_{1jj'},
\end{aligned} \tag{S4}$$

where we have used Equation (S2) for ①, approximated ② with $\mathbb{E}\left[\mathbf{x}_{1j}^T\mathbf{X}_1\mathbf{X}_1^T\mathbf{x}_{1j'}/n_1^2\right] = \mathbb{E}\left[\sum_{l=1}^p \mathbf{x}_{1j}^T\mathbf{x}_{1l}\mathbf{x}_{1l}^T\mathbf{x}_{1j'}/n_1^2\right] \approx \sum_{l=1}^p r_{1jl}r_{1j'l} + r_{1jj'}/n_1$, obtained ③ from $\mathrm{Var}(y_{1i}) = p\sum_{k=1}^K \sigma_{k1}^2 + p\omega_1 + \sigma_{\mathbf{e}_1}^2 = 1$, and the approximation ④ is granted by Equations (6). Similarly, we have $\mathrm{Cov}[\hat{\mathbf{b}}_{2j}, \hat{\mathbf{b}}_{2j'}] \approx (\sum_{k=1}^K \sigma_{k2}^2 + \omega_2)\sum_{l=1}^p r_{2jl}r_{2j'l} + \hat{s}_{2j}\hat{s}_{2j'}r_{2jj'}$. The covariance between elements of $\hat{\mathbf{b}}_1$ and $\hat{\mathbf{b}}_2$ can be derived as

$$
\begin{aligned}
&\mathrm{Cov}[\hat{\mathbf{b}}_{1j}, \hat{\mathbf{b}}_{2j'}] = \mathbb{E}\left[\hat{\mathbf{b}}_{1j}\hat{\mathbf{b}}_{2j'}\right] \\
=&\mathbb{E}\left[\mathbb{E}\left[\hat{\mathbf{b}}_{1j}\hat{\mathbf{b}}_{2j'}|\mathbf{X}_1, \mathbf{X}_2\right]\right] \\
=&\mathbb{E}\left[\mathbb{E}\left[\frac{1}{n_1n_2}\mathbf{x}_{1j}^T\mathbf{y}_1\mathbf{y}_2^T\mathbf{x}_{2j'}|\mathbf{X}_1, \mathbf{X}_2\right]\right] \\
=&\mathbb{E}\left[\mathbb{E}\left[\frac{1}{n_1n_2}\mathbf{x}_{1j}^T(\mathbf{X}_1\mathbf{b}_1 + \mathbf{X}_1\boldsymbol{\phi}_1 + \mathbf{e}_1)(\mathbf{X}_2\mathbf{b}_2 + \mathbf{X}_2\boldsymbol{\phi}_2 + \mathbf{e}_2)^T\mathbf{x}_{2j'}|\mathbf{X}_1, \mathbf{X}_2\right]\right] \\
=&\mathbb{E}\left[\mathbb{E}\left[\frac{1}{n_1n_2}\mathbf{x}_{1j}^T(\mathbf{X}_1\mathbf{b}_1\mathbf{b}_2^T\mathbf{X}_2^T + \mathbf{X}_1\boldsymbol{\phi}_1\boldsymbol{\phi}_2^T\mathbf{X}_2^T + \mathbf{e}_1\mathbf{e}_2^T|\mathbf{X}_1, \mathbf{X}_2\right]\right] \\
=&\mathbb{E}\left[\frac{1}{n_1n_2}\mathbf{x}_{1j}^T\mathbf{X}_1\mathbb{E}\left[\mathbf{b}_1\mathbf{b}_2^T\right]\mathbf{X}_2^T\mathbf{x}_{2j'} + \frac{1}{n_1n_2}\mathbf{x}_{1j}^T\mathbf{X}_1\mathbb{E}\left[\boldsymbol{\phi}_1\boldsymbol{\phi}_2^T\right]\mathbf{X}_2^T\mathbf{x}_{2j'} + \frac{1}{n_1n_2}\mathbf{x}_{1j}^T\mathbb{E}\left[\mathbf{e}_1\mathbf{e}_2^T\right]\mathbf{x}_{2j'}\right] \\
\overset{①}{=}&\frac{1}{n_1n_2}(\sum_{k=1}^K\sigma_{k12}^2)\mathbb{E}\left[\mathbf{x}_{1j}^T\mathbf{X}_1\mathbf{X}_2^T\mathbf{x}_{2j'}\right] + \frac{1}{n_1n_2}\omega_{12}\mathbb{E}\left[\mathbf{x}_{1j}^T\mathbf{X}_1\mathbf{X}_2^T\mathbf{x}_{2j'}\right] \\
=&(\sum_{k=1}^K\sigma_{k12}^2 + \omega_{12})\mathbb{E}\left[\mathbf{x}_{1j}^T\mathbf{X}_1\mathbf{X}_2^T\mathbf{x}_{2j'}/(n_1n_2)\right] \\
=&(\sum_{k=1}^K\sigma_{k12}^2 + \omega_{12})\sum_{l=1}^p r_{1jl}r_{2j'l},
\end{aligned}
\tag{S5}
$$

where the equation ① is obtained given the fact that $\mathbb{E}\left[\mathbf{e}_1\mathbf{e}_2^T\right] = \mathbf{0}$ because GWAS samples from two populations are independent. With above relationships, the variance-covariance matrices of $\hat{\mathbf{b}}_1$ and $\hat{\mathbf{b}}_2$ can be obtained as

$$
\mathrm{Var}\begin{bmatrix}\hat{\mathbf{b}}_1\\ \hat{\mathbf{b}}_2\end{bmatrix} = \mathcal{N}\left(\mathbf{0}, \begin{bmatrix}(\sum_{k=1}^K\sigma_{k1}^2 + \omega_1)\mathbf{R}_1^2 + \hat{\mathbf{S}}_1\mathbf{R}_1\hat{\mathbf{S}}_1 & (\sum_{k=1}^K\sigma_{k12}^2 + \omega_{12})\mathbf{R}_1\mathbf{R}_2 \\ (\sum_{k=1}^K\sigma_{k12}^2 + \omega_{12})\mathbf{R}_1\mathbf{R}_2 & (\sum_{k=1}^K\sigma_{k2}^2 + \omega_2)\mathbf{R}_2^2 + \hat{\mathbf{S}}_2\mathbf{R}_2\hat{\mathbf{S}}_2\end{bmatrix}\right). \tag{S6}
$$

Based on model (8), the variance-covariance matrices of $\boldsymbol{\epsilon}_1$ and $\boldsymbol{\epsilon}_2$ can be obtained as:

$$
\begin{aligned}
\mathrm{Var}\begin{bmatrix}\boldsymbol{\epsilon}_1\\ \boldsymbol{\epsilon}_2\end{bmatrix} =&\mathrm{Var}\begin{bmatrix}\hat{\mathbf{b}}_1\\ \hat{\mathbf{b}}_2\end{bmatrix} - \mathrm{Var}\begin{bmatrix}\mathbf{R}_1(\sum_{k=1}^K\gamma_k\beta_{1k} + \boldsymbol{\phi}_1)\\ \mathbf{R}_2(\sum_{k=1}^K\gamma_k\beta_{2k} + \boldsymbol{\phi}_2)\end{bmatrix} \\
=&\mathrm{Var}\begin{bmatrix}\hat{\mathbf{b}}_1\\ \hat{\mathbf{b}}_2\end{bmatrix} - \begin{bmatrix}(\sum_{k=1}^K\sigma_{k1}^2 + \omega_1)\mathbf{R}_1^2 & (\sum_{k=1}^K\sigma_{k12}^2 + \omega_{12})\mathbf{R}_1\mathbf{R}_2 \\ (\sum_{k=1}^K\sigma_{k12}^2 + \omega_{12})\mathbf{R}_1\mathbf{R}_2 & (\sum_{k=1}^K\sigma_{k2}^2 + \omega_2)\mathbf{R}_2^2\end{bmatrix} \\
=&\begin{bmatrix}\hat{\mathbf{S}}_1\mathbf{R}_1\hat{\mathbf{S}}_1 & \mathbf{0}\\ \mathbf{0} & \hat{\mathbf{S}}_2\mathbf{R}_2\hat{\mathbf{S}}_2\end{bmatrix}.
\end{aligned}
\tag{S7}
$$

Considering the large sample size of GWASs, we can obtain the asymptotic normal distribution in Equation (9).

##### 3.4 The XMAP model accounting for sample structure

Here, we derive XMAP under the genetic drift model in Equation (10). To model the population stratification, we assume that the samples from population 1 are constructed by a 50:50 mixture of sub-population $1a$ and sub-population $1b$, and the samples from population 2 are constructed by a 50:50 mixture of sub-population $2a$ and sub-population $2b$. We use $F_{ST,1}$ to denote the allele frequency difference between sub-population $1a$ and sub-population $1b$ and use $\sigma_{d_1}$ to denote the mean phenotype difference between sub-population $1a$ and sub-population $1b$. Similarly, we use $F_{ST,2}$ and $\sigma_{d_2}$ to denote the allele frequency difference and mean phenotype difference between the two sub-populations of population 2, respectively. To account for the sample structures, we consider an extension of model (1) as follows:

$$\begin{aligned} \mathbf{y}_1 &= \mathbf{X}_1\mathbf{b}_1 + \mathbf{X}_1\boldsymbol{\phi}_1 + \mathbf{d}_1 + \mathbf{e}_1, \\ \mathbf{y}_2 &= \mathbf{X}_2\mathbf{b}_2 + \mathbf{X}_2\boldsymbol{\phi}_2 + \mathbf{d}_2 + \mathbf{e}_2, \end{aligned} \tag{S8}$$

where $\mathbf{d}_1 \sin \mathbb{R}^{n_1}$ and $\mathbf{d}_2 \sin \mathbb{R}^{n_2}$ are the environmental stratification terms defined as

$$\begin{aligned} d_{1,i} &= \begin{cases} \sigma_{d_1}, & i \in \text{ sub-population } 1a \\ -\sigma_{d_1}, & i \in \text{ sub-population } 1b \end{cases}, \quad i = 1, \cdots n_1 \ , \\ d_{2,i} &= \begin{cases} \sigma_{d_2}, & i \in \text{ sub-population } 2a \\ -\sigma_{d_2}, & i \in \text{ sub-population } 2b \end{cases}, \quad i = 1, \cdots n_2 \ . \end{aligned} \tag{S9}$$

We also assume that the $\mathbf{d}_1$ is independent of $\mathbf{e}_1$ and $\mathbf{d}_2$ is independent of $\mathbf{e}_1$, and that $\mathrm{Var}(y_{1i}) = p\sum_{k=1}^K \sigma_{k1}^2 + p\omega_1 + \sigma_{d_1}^2 + \sigma_{\mathbf{e}_1}^2 = 1$ and $\mathrm{Var}(y_{2i}) = p\sum_{k=1}^K \sigma_{k2}^2 + p\omega_2 + \sigma_{d_2}^2 + \sigma_{\mathbf{e}_2}^2 = 1$. Using the results of bivariate LDSC [1, 2], we have

$$\begin{aligned} \mathrm{Cov}[\hat{\mathbf{b}}_{1j}, \hat{\mathbf{b}}_{1j'}] &= (\sum_{k=1}^K \sigma_{k1}^2 + \omega_1) \sum_{l=1}^p r_{1jl} r_{1j'l} + \underbrace{(1 + n_1 F_{ST,1}(h_1^2 F_{ST,1} + \sigma_{d_1}^2))}_{c_1} \hat{s}_{b,1j}\hat{s}_{b,1j'} r_{1jj'} \\ \mathrm{Cov}[\hat{\mathbf{b}}_{2j}, \hat{\mathbf{b}}_{2j'}] &= (\sum_{k=1}^K \sigma_{k2}^2 + \omega_2) \sum_{l=1}^p r_{2jl} r_{2j'l} + \underbrace{(1 + n_2 F_{ST,2}(h_2^2 F_{ST,2} + \sigma_{d_2}^2))}_{c_2} \hat{s}_{b,2j}\hat{s}_{b,2j'} r_{2jj'}, \end{aligned} \tag{S10}$$

where $h_1^2 = p\sum_{k=1}^K \sigma_{k1}^2 + p\omega_1$ and $h_2^2 = p\sum_{k=1}^K \sigma_{k2}^2 + p\omega_2$ are heritabilities of $\mathbf{y}_1$ and $\mathbf{y}_2$, respectively. Then, we can update Equation (S7) as

$$\mathrm{Var}\begin{bmatrix} \boldsymbol{\epsilon}_1 \\ \boldsymbol{\epsilon}_2 \end{bmatrix} = \begin{bmatrix} c_1\hat{\mathbf{S}}_1\mathbf{R}_1\hat{\mathbf{S}}_1 & \mathbf{0} \\ \mathbf{0} & c_2\hat{\mathbf{S}}_2\mathbf{R}_2\hat{\mathbf{S}}_2 \end{bmatrix}. \tag{S11}$$

As we can observe, the inflation constants $c_1$ and $c_2$ can be greater than one in the presence of population stratification ($F_{ST,1} \neq 0$ and $F_{ST,2} \neq 0$, respectively).

##### 3.5 Derivation of the variational EM algorithm of XMAP

We derive the variational EM algorithm to obtain the estimate of parameters $\boldsymbol{\Sigma}$ and the approximate posterior $q(\boldsymbol{\gamma}, \boldsymbol{\beta}, \boldsymbol{\phi})$. For simplicity of notation, we suppress the pre-estimated parameters $\{\hat{\boldsymbol{\Omega}}, \hat{c}_1, \hat{c}_2\}$ in the following derivation.

The complete-data log-likelihood is given as

$$
\begin{aligned}
&\Pr(\hat{\mathbf{b}}_1, \hat{\mathbf{b}}_2, \boldsymbol{\gamma}, \boldsymbol{\beta}, \boldsymbol{\phi}|\boldsymbol{\Sigma}) \\
=&\log[\Pr(\hat{\mathbf{b}}_1|\boldsymbol{\gamma}, \boldsymbol{\beta}, \boldsymbol{\phi}) \Pr(\hat{\mathbf{b}}_2|\boldsymbol{\gamma}, \boldsymbol{\beta}, \boldsymbol{\phi}) \prod_k^K \Pr(\beta_{1k}, \beta_{2k}) \prod_k^K \Pr(\boldsymbol{\gamma}_k) \Pr(\boldsymbol{\phi}_1, \boldsymbol{\phi}_2)] \\
=&-\frac{1}{2\hat{c}_1}(\hat{\mathbf{b}}_1 - \sum_k^K \mathbf{R}_1\boldsymbol{\gamma}_k\beta_{1k} - \mathbf{R}_1\boldsymbol{\phi}_1)^T(\hat{\mathbf{S}}_1\mathbf{R}_1\hat{\mathbf{S}}_1)^{-1}(\hat{\mathbf{b}}_1 - \sum_k^K \mathbf{R}_1\boldsymbol{\gamma}_k\beta_{1k} - \mathbf{R}_1\boldsymbol{\phi}_1) \\
&-\frac{1}{2\hat{c}_2}(\hat{\mathbf{b}}_2 - \sum_k^K \mathbf{R}_2\boldsymbol{\gamma}_k\beta_{2k} - \mathbf{R}_2\boldsymbol{\phi}_2)^T(\hat{\mathbf{S}}_2\mathbf{R}_2\hat{\mathbf{S}}_2)^{-1}(\hat{\mathbf{b}}_2 - \sum_k^K \mathbf{R}_2\boldsymbol{\gamma}_k\beta_{2k} - \mathbf{R}_2\boldsymbol{\phi}_2)' \\
&-\frac{1}{2}\sum_k^K \log(2\pi)^2|\boldsymbol{\Sigma}_k| - \frac{1}{2}\sum_k^K \begin{bmatrix}\beta_{1k} & \beta_{2k}\end{bmatrix} \boldsymbol{\Sigma}_k^{-1} \begin{bmatrix}\beta_{1k} \\ \beta_{2k}\end{bmatrix} + \sum_j^p\sum_k^K \gamma_{kj}\log\frac{1}{p} \\
&-\frac{p}{2}\log|2\pi\hat{\boldsymbol{\Omega}}| - \frac{1}{2}\sum_{j=1}^p \begin{bmatrix}\phi_{1j} & \phi_{2j}\end{bmatrix} \hat{\boldsymbol{\Omega}}^{-1} \begin{bmatrix}\phi_{1j} \\ \phi_{2j}\end{bmatrix} + \text{constant}
\end{aligned}
\tag{S12}
$$

where the constant term do not depend on $\mathbf{b}$ and $\boldsymbol{\phi}$. In practice, the LD matrices $\mathbf{R}_1$ and $\mathbf{R}_2$ can be estimated with population-matched reference samples. However, the LD matrices may not be invertible when some SNPs are in perfect LD or the number of individuals in the reference panel is less than $p$. To address this difficulty, we define the XMAP likelihood by discarding the terms that do not depend on $\mathbf{b}$ and $\boldsymbol{\phi}$:

$$
\begin{aligned}
&\mathcal{L}(\boldsymbol{\Sigma}) \\
=&-\frac{1}{2\hat{c}_1}(-2(\sum_k^K \boldsymbol{\gamma}_k\beta_{1k} + \boldsymbol{\phi}_1)^T\hat{\mathbf{S}}_1^{-2}\hat{\mathbf{b}}_1 + (\sum_k^K \boldsymbol{\gamma}_k\beta_{1k} + \boldsymbol{\phi}_1)^T\hat{\mathbf{S}}_1^{-1}\mathbf{R}_1\hat{\mathbf{S}}_1^{-1}(\sum_k^K \boldsymbol{\gamma}_k\beta_{1k} + \boldsymbol{\phi}_1)) \\
&-\frac{1}{2\hat{c}_2}(-2(\sum_k^K \boldsymbol{\gamma}_k\beta_{2k} + \boldsymbol{\phi}_2)^T\hat{\mathbf{S}}_2^{-2}\hat{\mathbf{b}}_2 + (\sum_k^K \boldsymbol{\gamma}_k\beta_{2k} + \boldsymbol{\phi}_2)^T\hat{\mathbf{S}}_2^{-1}\mathbf{R}_2\hat{\mathbf{S}}_2^{-1}(\sum_k^K \boldsymbol{\gamma}_k\beta_{2k} + \boldsymbol{\phi}_2)) \\
&-\frac{1}{2}\sum_k^K \log(2\pi)^2|\boldsymbol{\Sigma}_k| - \frac{1}{2}\sum_k^K \begin{bmatrix}\beta_{1k} & \beta_{2k}\end{bmatrix} \boldsymbol{\Sigma}_k^{-1} \begin{bmatrix}\beta_{1k} \\ \beta_{2k}\end{bmatrix} + \sum_j^p\sum_k^K \boldsymbol{\gamma}_{kj}\log\frac{1}{p} \\
&-\frac{p}{2}\log|2\pi\hat{\boldsymbol{\Omega}}| - \frac{1}{2}\sum_{j=1}^p \begin{bmatrix}\phi_{1j} & \phi_{2j}\end{bmatrix} \hat{\boldsymbol{\Omega}}^{-1} \begin{bmatrix}\phi_{1j} \\ \phi_{2j}\end{bmatrix}.
\end{aligned}
\tag{S13}
$$

As we can observe, this definition of likelihood allows our algorithm to handle non-invertible LD matrices because it does not depend on $\mathbf{R}_1^{-1}$ and $\mathbf{R}_2^{-1}$.

**E-step** Based on the mean field assumption for the variational distribution (15), we can derive the approximated posterior $q(\boldsymbol{\phi})$:

$$\begin{aligned}\log q\left(\begin{bmatrix}\boldsymbol{\phi}_1\\ \boldsymbol{\phi}_2\end{bmatrix}\right) &= \begin{bmatrix}\boldsymbol{\phi}_1 & \boldsymbol{\phi}_2\end{bmatrix}\begin{bmatrix}\frac{1}{\hat{c}_1}\hat{\mathbf{S}}_1^{-2}\hat{\mathbf{b}}_1 - \frac{1}{\hat{c}_1}\hat{\mathbf{S}}_b^{-1}\mathbf{R}_1\hat{\mathbf{S}}_1^{-1}\sum_{k=1}^K \mathbb{E}_{q_k}(\boldsymbol{\gamma}_k\beta_{1k})\\ \frac{1}{\hat{c}_2}\hat{\mathbf{S}}_2^{-2}\hat{\mathbf{b}}_2 - \frac{1}{\hat{c}_2}\hat{\mathbf{S}}_2^{-1}\mathbf{R}_2\hat{\mathbf{S}}_2^{-1}\sum_{k=1}^K \mathbb{E}_{q_k}(\boldsymbol{\gamma}_k\beta_{2k})\end{bmatrix}\\ &\quad - \frac{1}{2}\begin{bmatrix}\boldsymbol{\phi}_1 & \boldsymbol{\phi}_2\end{bmatrix}\left(\begin{bmatrix}\frac{1}{\hat{c}_1}\hat{\mathbf{S}}_{b,1}^{-1}\mathbf{R}_1\hat{\mathbf{S}}_{b,1}^{-1} & \mathbf{0}\\ \mathbf{0} & \frac{1}{\hat{c}_2}\hat{\mathbf{S}}_2^{-1}\mathbf{R}_2\hat{\mathbf{S}}_2^{-1}\end{bmatrix} + \hat{\boldsymbol{\Omega}}^{-1}\otimes\mathbf{I}_p\right)\begin{bmatrix}\boldsymbol{\phi}_1\\ \boldsymbol{\phi}_2\end{bmatrix}\\ &\quad + \text{constant},\end{aligned} \tag{S14}$$

where $\otimes$ denotes the Kronecker product, and the expectation is taken under the distribution $q(\boldsymbol{\gamma}_k)$ and $q(\beta_{1k}, \beta_{2k}|\boldsymbol{\gamma}_k)$ for $k = 1, .., K$. From the quadratic form of Equation (S14), we know that $q(\boldsymbol{\phi})$ follows the normal distribution:

$$\begin{bmatrix}\boldsymbol{\phi}_1\\ \boldsymbol{\phi}_2\end{bmatrix} \sim \mathcal{N}(\tilde{\boldsymbol{\nu}}, \tilde{\boldsymbol{\Lambda}}), \tag{S15}$$

where

$$\begin{aligned}\tilde{\boldsymbol{\Lambda}} &= \left(\begin{bmatrix}\frac{1}{\hat{c}_1}\hat{\mathbf{S}}_1^{-1}\mathbf{R}_1\hat{\mathbf{S}}_1^{-1} & \mathbf{0}\\ \mathbf{0} & \frac{1}{\hat{c}_2}\hat{\mathbf{S}}_2^{-1}\mathbf{R}_2\hat{\mathbf{S}}_2^{-1}\end{bmatrix} + \hat{\boldsymbol{\Omega}}^{-1}\otimes\mathbf{I}_p\right)^{-1},\\ \tilde{\boldsymbol{\nu}} &= \tilde{\boldsymbol{\Lambda}}\begin{bmatrix}\frac{1}{\hat{c}_1}\hat{\mathbf{S}}_1^{-2}\hat{\mathbf{b}}_1 - \frac{1}{\hat{c}_1}\hat{\mathbf{S}}_1^{-1}\mathbf{R}_1\hat{\mathbf{S}}_1^{-1}\sum_{k=1}^K \mathbb{E}_{q_k}(\boldsymbol{\gamma}_k\beta_{1k})\\ \frac{1}{\hat{c}_2}\hat{\mathbf{S}}_2^{-2}\hat{\mathbf{b}}_2 - \frac{1}{\hat{c}_2}\hat{\mathbf{S}}_2^{-1}\mathbf{R}_2\hat{\mathbf{S}}_2^{-1}\sum_{k=1}^K \mathbb{E}_{q_k}(\boldsymbol{\gamma}_k\beta_{2k})\end{bmatrix}.\end{aligned} \tag{S16}$$

Similarly, $\log q(\beta_{1k}, \beta_{2k}|\gamma_{kj} = 1)$ can be obtained as

$$\begin{aligned}\log q\left(\begin{bmatrix}\beta_{1k}\\ \beta_{2k}\end{bmatrix}|\gamma_{kj} = 1\right) &= \begin{bmatrix}\beta_{1k} & \beta_{2k}\end{bmatrix}\begin{bmatrix}\frac{\hat{\mathbf{b}}_{1j}}{\hat{c}_1\hat{s}_{1j}^2} - \frac{1}{\hat{c}_1\hat{s}_{1j}^2}\mathbf{R}_{1j}^T(\sum_{k'\neq 1}^K \mathbb{E}_{q_{k'}}(\boldsymbol{\gamma}_{k'}\beta_{1k'}) + \mathbb{E}_{q_\phi}(\boldsymbol{\phi}_1))\\ \frac{\hat{\mathbf{b}}_{2j}}{\hat{c}_2\hat{s}_{2j}^2} - \frac{1}{\hat{c}_2\hat{s}_{2j}^2}\mathbf{R}_{2j}^T(\sum_{k'\neq 1}^K \mathbb{E}_{q_{k'}}(\boldsymbol{\gamma}_{k'}\beta_{2k'}) + \mathbb{E}_{q_\phi}(\boldsymbol{\phi}_2))\end{bmatrix}\\ &\quad - \frac{1}{2}\begin{bmatrix}\beta_{1k} & \beta_{2k}\end{bmatrix}\left(\begin{bmatrix}\frac{r_{1jj}}{\hat{c}_1\hat{s}_{1j}^2} & \mathbf{0}\\ \mathbf{0} & \frac{r_{2jj}}{\hat{c}_2\hat{s}_{2j}^2}\end{bmatrix} + \boldsymbol{\Sigma}_k^{-1}\right)\begin{bmatrix}\beta_{1k}\\ \beta_{2k}\end{bmatrix}\\ &\quad + \text{constant},\end{aligned} \tag{S17}$$

where $\mathbf{R}_{1j} = [r_{1j1}, ..., r_{1jp}]^T$ and $\mathbf{R}_{2j} = [r_{2j1}, ..., r_{2jp}]^T$, the expectation $\mathbb{E}_{q_{k'}}$ is taken under the distributions $q(\boldsymbol{\gamma}_{k'})$ and $q(\beta_{1k'}, \beta_{2k'}|\boldsymbol{\gamma}_k)$ for $k' \neq k$, and the expectation $\mathbb{E}_\phi$ is taken under the distribution $q(\boldsymbol{\phi})$. The expression in Equation (S17) indicates that $q(\beta_{1k}, \beta_{2k}|\gamma_{kj} = 1)$ follows the normal distribution:

$$\begin{bmatrix}\beta_{1k}\\ \beta_{2k}\end{bmatrix}|\gamma_{kj} = 1 \sim \mathcal{N}(\tilde{\boldsymbol{\mu}}_{kj}, \tilde{\boldsymbol{\Sigma}}_{kj}), \tag{S18}$$

where

$$\begin{aligned}\tilde{\boldsymbol{\Sigma}}_{kj} &= \begin{bmatrix}\tilde{\sigma}_{kj,1}^2 & \tilde{\sigma}_{kj,12}^2\\ \tilde{\sigma}_{kj,2}^2 & \tilde{\sigma}_{kj,2}^2\end{bmatrix} = \left(\begin{bmatrix}\frac{r_{1jj}}{\hat{c}_1\hat{s}_{b,1j}^2} & \mathbf{0}\\ \mathbf{0} & \frac{r_{2jj}}{\hat{c}_2\hat{s}_{b,2j}^2}\end{bmatrix} + \boldsymbol{\Sigma}_k^{-1}\right)^{-1},\\ \tilde{\boldsymbol{\mu}}_{kj} &= \begin{bmatrix}\tilde{\mu}_{kj,1}\\ \tilde{\mu}_{kj,2}\end{bmatrix} = \tilde{\boldsymbol{\Sigma}}_{kj}\begin{bmatrix}\frac{\hat{\mathbf{b}}_{1j}}{\hat{c}_1\hat{s}_{1j}^2} - \frac{1}{\hat{c}_1\hat{s}_{1j}^2}\mathbf{R}_{1j}^T(\sum_{k'\neq 1}^K \mathbb{E}_{q_{k'}}(\boldsymbol{\gamma}_{k'}\beta_{1k'}) + \mathbb{E}_{q_\phi}(\boldsymbol{\phi}_1))\\ \frac{\hat{\mathbf{b}}_{2j}}{\hat{c}_2\hat{s}_{2j}^2} - \frac{1}{\hat{c}_2\hat{s}_{2j}^2}\mathbf{R}_{2j}^T(\sum_{k'\neq 1}^K \mathbb{E}_{q_{k'}}(\boldsymbol{\gamma}_{k'}\beta_{2k'}) + \mathbb{E}_{q_\phi}(\boldsymbol{\phi}_2))\end{bmatrix}.\end{aligned} \tag{S19}$$

Let $q(\boldsymbol{\gamma}_k) = \tilde{\boldsymbol{\pi}} = [\tilde{\pi}_{k1}, ..., \tilde{\pi}_{kp}]^T$, where $\sum_{j=1}^{p} \tilde{\pi}_{kj} = 1$ for $k = 1, .., K$. We can obtain $q(\boldsymbol{\gamma}, \boldsymbol{\beta}, \boldsymbol{\phi})$ as

$$q(\boldsymbol{\gamma}, \boldsymbol{\beta}, \boldsymbol{\phi}) = \prod_{k}^{K} q(\boldsymbol{\gamma}_k) q(\beta_{1k}, \beta_{2k} | \boldsymbol{\gamma}_k) q(\boldsymbol{\phi}_1, \boldsymbol{\phi}_2) = \prod_{k}^{K} \prod_{j}^{p} \left[ \tilde{\boldsymbol{\pi}}_{kj} \mathcal{N}(\tilde{\boldsymbol{\mu}}_{kj}, \tilde{\boldsymbol{\Sigma}}_{kj}) \right]^{\gamma_{kj}} \mathcal{N}(\tilde{\boldsymbol{\nu}}, \tilde{\boldsymbol{\Lambda}}). \quad \text{(S20)}$$

Then, we can evaluate the lower bound given by Equation (13)

$$\begin{aligned}
&\mathcal{L}_q(\boldsymbol{\Sigma}) = \mathbb{E}_q[\mathcal{L}(\boldsymbol{\Sigma})] - \mathbb{E}_q[q(\boldsymbol{\gamma}, \boldsymbol{\beta}, \boldsymbol{\phi}))] \\
&= \left( \sum_{k}^{K} \tilde{\boldsymbol{\mu}}_{kj} \otimes \tilde{\boldsymbol{\pi}}_k + \tilde{\boldsymbol{\nu}} \right)^T \begin{bmatrix} \frac{\hat{\mathbf{S}}_1^{-2} \hat{\mathbf{b}}_1}{\hat{c}_1} \\ \frac{\hat{\mathbf{S}}_2^{-2} \hat{\mathbf{b}}_2}{\hat{c}_2} \end{bmatrix} - \frac{1}{2} \left( \sum_{k}^{K} \tilde{\boldsymbol{\mu}}_{kj} \otimes \tilde{\boldsymbol{\pi}}_k + \tilde{\boldsymbol{\nu}} \right)^T \begin{bmatrix} \frac{\hat{\mathbf{S}}_1^{-1} \mathbf{R}_1 \hat{\mathbf{S}}_1^{-1}}{\hat{c}_1} & \mathbf{0} \\ \mathbf{0} & \frac{\hat{\mathbf{S}}_2^{-1} \mathbf{R}_2 \hat{\mathbf{S}}_2^{-1}}{\hat{c}_2} \end{bmatrix} \left( \sum_{k}^{K} \tilde{\boldsymbol{\mu}}_{kj} \otimes \tilde{\boldsymbol{\pi}}_k + \tilde{\boldsymbol{\nu}} \right) \\
&\quad - \sum_{j}^{p} \frac{1}{2\hat{c}_1 \hat{s}_{1j}^2} r_{1jj} \sum_{k}^{K} \tilde{\pi}_{kj} (\tilde{\mu}_{kj,1}^2 + \tilde{\sigma}_{kj,1}^2) - \sum_{j}^{p} \frac{1}{2\hat{c}_2 \hat{s}_{2j}^2} r_{2jj} \sum_{k}^{K} \tilde{\pi}_{kj} (\tilde{\mu}_{kj,2}^2 + \tilde{\sigma}_{kj,2}^2) \\
&\quad + \frac{1}{2} \sum_{k}^{K} \left( (\tilde{\boldsymbol{\mu}}_{kj} \otimes \tilde{\boldsymbol{\pi}}_k)^T \begin{bmatrix} \frac{\hat{\mathbf{S}}_1^{-1} \mathbf{R}_1 \hat{\mathbf{S}}_1^{-1}}{\hat{c}_1} & \mathbf{0} \\ \mathbf{0} & \frac{\hat{\mathbf{S}}_2^{-1} \mathbf{R}_2 \hat{\mathbf{S}}_2^{-1}}{\hat{c}_2} \end{bmatrix} (\tilde{\boldsymbol{\mu}}_{kj} \otimes \tilde{\boldsymbol{\pi}}_k) \right) \\
&\quad - \frac{1}{2} \sum_{k}^{K} \log(2\pi)^2 |\boldsymbol{\Sigma}_k| - \frac{1}{2} \sum_{k} \sum_{j} \tilde{\pi}_{kj} \mathrm{Tr}(\boldsymbol{\Sigma}_k^{-1} (\tilde{\boldsymbol{\Sigma}}_{kj} + \tilde{\boldsymbol{\mu}}_{kj} \tilde{\boldsymbol{\mu}}_{kj}^T)) + \sum_{j}^{p} \sum_{k}^{K} \tilde{\pi}_{kj} \log \frac{1}{p} \\
&\quad - \frac{p}{2} \log |2\pi \hat{\boldsymbol{\Omega}}| - \frac{1}{2} \tilde{\boldsymbol{\nu}}^T (\hat{\boldsymbol{\Omega}}^{-1} \otimes \mathbf{I}_p) \tilde{\boldsymbol{\nu}} - \frac{1}{2} \mathrm{Tr} \left( \left( \begin{bmatrix} \frac{1}{\hat{c}_1} \hat{\mathbf{S}}_1^{-1} \mathbf{R}_1 \hat{\mathbf{S}}_1^{-1} & \mathbf{0} \\ \mathbf{0} & \frac{1}{\hat{c}_2} \hat{\mathbf{S}}_2^{-1} \mathbf{R}_2 \hat{\mathbf{S}}_2^{-1} \end{bmatrix} + \hat{\boldsymbol{\Omega}}^{-1} \otimes \mathbf{I}_p \right) \tilde{\boldsymbol{\Lambda}} \right) \\
&\quad - \sum_{j}^{p} \sum_{k}^{K} \tilde{\pi}_{kj} \log \tilde{\pi}_{kj} + \frac{1}{2} \sum_{j}^{p} \sum_{k}^{K} \tilde{\pi}_{kj} \log |\tilde{\boldsymbol{\Sigma}}_{kj}| + K + K \log(2\pi) + \frac{1}{2} \log |\tilde{\boldsymbol{\Lambda}}| + p + p \log 2\pi \\
&= \left( \sum_{k}^{K} \tilde{\boldsymbol{\mu}}_{kj} \otimes \tilde{\boldsymbol{\pi}}_k + \tilde{\boldsymbol{\nu}} \right)^T \begin{bmatrix} \frac{\hat{\mathbf{S}}_1^{-2} \hat{\mathbf{b}}_1}{\hat{c}_1} \\ \frac{\hat{\mathbf{S}}_2^{-2} \hat{\mathbf{b}}_2}{\hat{c}_2} \end{bmatrix} - \frac{1}{2} \left( \sum_{k}^{K} \tilde{\boldsymbol{\mu}}_{kj} \otimes \tilde{\boldsymbol{\pi}}_k + \tilde{\boldsymbol{\nu}} \right)^T \begin{bmatrix} \frac{\hat{\mathbf{S}}_1^{-1} \mathbf{R}_1 \hat{\mathbf{S}}_1^{-1}}{\hat{c}_1} & \mathbf{0} \\ \mathbf{0} & \frac{\hat{\mathbf{S}}_2^{-1} \mathbf{R}_2 \hat{\mathbf{S}}_2^{-1}}{\hat{c}_2} \end{bmatrix} \left( \sum_{k}^{K} \tilde{\boldsymbol{\mu}}_{kj} \otimes \tilde{\boldsymbol{\pi}}_k + \tilde{\boldsymbol{\nu}} \right) \\
&\quad - \sum_{j}^{p} \frac{1}{2\hat{c}_1 \hat{s}_{1j}^2} r_{1jj} \sum_{k}^{K} \tilde{\pi}_{kj} (\tilde{\mu}_{kj,1}^2 + \tilde{\sigma}_{kj,1}^2) - \sum_{j}^{p} \frac{1}{2\hat{c}_2 \hat{s}_{2j}^2} r_{2jj} \sum_{k}^{K} \tilde{\pi}_{kj} (\tilde{\mu}_{kj,2}^2 + \tilde{\sigma}_{kj,2}^2) \\
&\quad + \frac{1}{2} \sum_{k}^{K} \left( (\tilde{\boldsymbol{\mu}}_{kj} \otimes \tilde{\boldsymbol{\pi}}_k)^T \begin{bmatrix} \frac{\hat{\mathbf{S}}_1^{-1} \mathbf{R}_1 \hat{\mathbf{S}}_1^{-1}}{\hat{c}_1} & \mathbf{0} \\ \mathbf{0} & \frac{\hat{\mathbf{S}}_2^{-1} \mathbf{R}_2 \hat{\mathbf{S}}_2^{-1}}{\hat{c}_2} \end{bmatrix} (\tilde{\boldsymbol{\mu}}_{kj} \otimes \tilde{\boldsymbol{\pi}}_k) \right) \\
&\quad - \frac{1}{2} \sum_{k} \sum_{j} \boldsymbol{\gamma}_{kj} \mathrm{Tr}(\boldsymbol{\Sigma}_k^{-1} (\tilde{\boldsymbol{\Sigma}}_{kj} + \tilde{\boldsymbol{\mu}}_{kj} \tilde{\boldsymbol{\mu}}_{kj}^T)) \\
&\quad - \frac{p}{2} \log |2\pi \hat{\boldsymbol{\Omega}}| - \frac{1}{2} \tilde{\boldsymbol{\nu}}^T (\hat{\boldsymbol{\Omega}}^{-1} \otimes \mathbf{I}_p) \tilde{\boldsymbol{\nu}} - \frac{1}{2} \mathrm{Tr} \left( \left( \begin{bmatrix} \frac{1}{\hat{c}_1} \hat{\mathbf{S}}_1^{-1} \mathbf{R}_1 \hat{\mathbf{S}}_1^{-1} & \mathbf{0} \\ \mathbf{0} & \frac{1}{\hat{c}_2} \hat{\mathbf{S}}_2^{-1} \mathbf{R}_2 \hat{\mathbf{S}}_2^{-1} \end{bmatrix} + \hat{\boldsymbol{\Omega}}^{-1} \otimes \mathbf{I}_p \right) \tilde{\boldsymbol{\Lambda}} \right) \\
&\quad + \sum_{j}^{p} \sum_{k}^{K} \tilde{\pi}_{kj} \log \frac{1}{p} - \sum_{j}^{p} \sum_{k}^{K} \tilde{\pi}_{kj} \log \tilde{\pi}_{kj} + \frac{1}{2} \sum_{j}^{p} \sum_{k}^{K} [\tilde{\pi}_{kj} (\log |\tilde{\boldsymbol{\Sigma}}_{kj}| - \log |\boldsymbol{\Sigma}_k|)] + \frac{1}{2} \log |\tilde{\boldsymbol{\Lambda}}| \\
&\quad + \text{constant.}
\end{aligned} \quad \text{(S21)}$$

By setting the derivative of the lower bound w.r.t $\tilde{\pi}_{kj}$ as zero, we can get the update of variational parameter $\gamma_{kj}$:

$$\tilde{\pi}_{kj} = softmax(\log\frac{1}{p} + \frac{1}{2}\log|\tilde{\mathbf{\Sigma}}_{kj}| + \frac{1}{2}\tilde{\boldsymbol{\mu}}_{kj}^T\tilde{\mathbf{\Sigma}}_{kj}^{-1}\tilde{\boldsymbol{\mu}}_{kj}), \tag{S22}$$

where $softmax$ is the softmax function to make sure $\sum_{j=1}^{p}\tilde{\pi}_{kj} = 1$. Combining, we can obtain the updating equations of the variational parameters in Equation (16).

**M-step** At M-step, we set $\frac{\partial\mathcal{L}_q}{\partial\mathbf{\Sigma}_k} = 0$ to obtain the update equation of $\mathbf{\Sigma}_k$:

$$\mathbf{\Sigma}_k = \sum_{j}^{p}\tilde{\pi}_{kj}(\tilde{\boldsymbol{\mu}}_{kj}\tilde{\boldsymbol{\mu}}_{kj}^T + \tilde{\mathbf{\Sigma}}_{kj}). \tag{S23}$$
